# High Resolution Genome Wide Expression Analysis of Single Myofibers Using SMART-Seq

**DOI:** 10.1101/724393

**Authors:** Darren M. Blackburn, Felicia Lazure, Aldo H. Corchado, Theodore J. Perkins, Hamed S. Najafabadi, Vahab D. Soleimani

**Author notes:** These authors made equal contributions (Darren M. Blackburn, Felicia Lazure). To whom correspondence should be addressed: Vahab D. Soleimani: Department of Human Genetics, McGill University, 3640 rue University, Montreal, QC, H3A 0C7, Canada.

## Abstract

Skeletal muscle is a heterogeneous tissue. Individual myofibers that make up muscle tissue exhibit variation in their metabolic and contractile properties. Although there are biochemical and histological assays to study myofiber heterogeneity, efficient methods to analyze the whole transcriptome of individual myofibers are lacking. We have developed single myofiber RNA-Seq (smfRNA-Seq) to analyze the whole transcriptome of individual myofibers by combining single fiber isolation with Switching Mechanisms at 5’ end of RNA Template (SMART) technology. Our method provides high-resolution genome wide expression profiles of single myofibers. Using smfRNA-Seq, we have analyzed the differences in the transcriptome of young and old myofibers to validate the effectiveness of this new method. Using smfRNA-Seq, we performed comparative gene expression analysis between single myofibers from young and old mice. Our data suggests that aging leads to significant changes in the expression of metabolic and structural genes in myofibers. Our data suggests that smfRNA-Seq is a powerful tool to study developmental, disease and age-related dynamics in the composition of skeletal muscle.

## INTRODUCTION

Skeletal muscle is composed of a variety of different cell types, including endothelial cells, fibro/adipogenic cells (FAPs), adipocytes, mesenchymal cells and fibroblasts, among others(1–4). Further heterogeneity of skeletal muscle is also manifested by the diversity in the composition of myofiber types which constitute muscles (5). Skeletal muscle fiber types are often categorized based on their contractile properties, giving two broad categories: fast twitch muscles and slow twitch muscles(5, 6). These fiber types can be further subcategorized based on metabolic properties and myosin heavy chain (MyHC) isoforms(5, 7).

Methods of investigating changes in fiber type in response to different stimuli rely on staining and biochemical analyses of individual fibers, biochemical analyses of the whole muscle or sequencing of entire muscles(8). Standard bulk RNA sequencing is not suited for the analysis of myofibers at high resolution, as it captures the entirety of the muscle tissue, resulting in the pooling of different fiber types in addition to non-myogenic cells (8). Consequently, in such studies, the myofiber-specific gene signature cannot be inferred based on RNA sequencing of whole muscle tissue. The emergence of single cell technology provides ample opportunity for further investigation into the heterogeneity of single muscle fibers at the transcriptome level. Here, we combine single myofiber isolation of the Extensor Digitorum Longus (EDL) in *Mus musculus* with Switching Mechanism at the 5’ end of RNA Template (SMART) technology to analyze the whole transcriptome of individual myofibers (Figure 1)(9), in a method called Single MyoFiber RNA-Seq (smfRNA-Seq).

**Figure 1:**
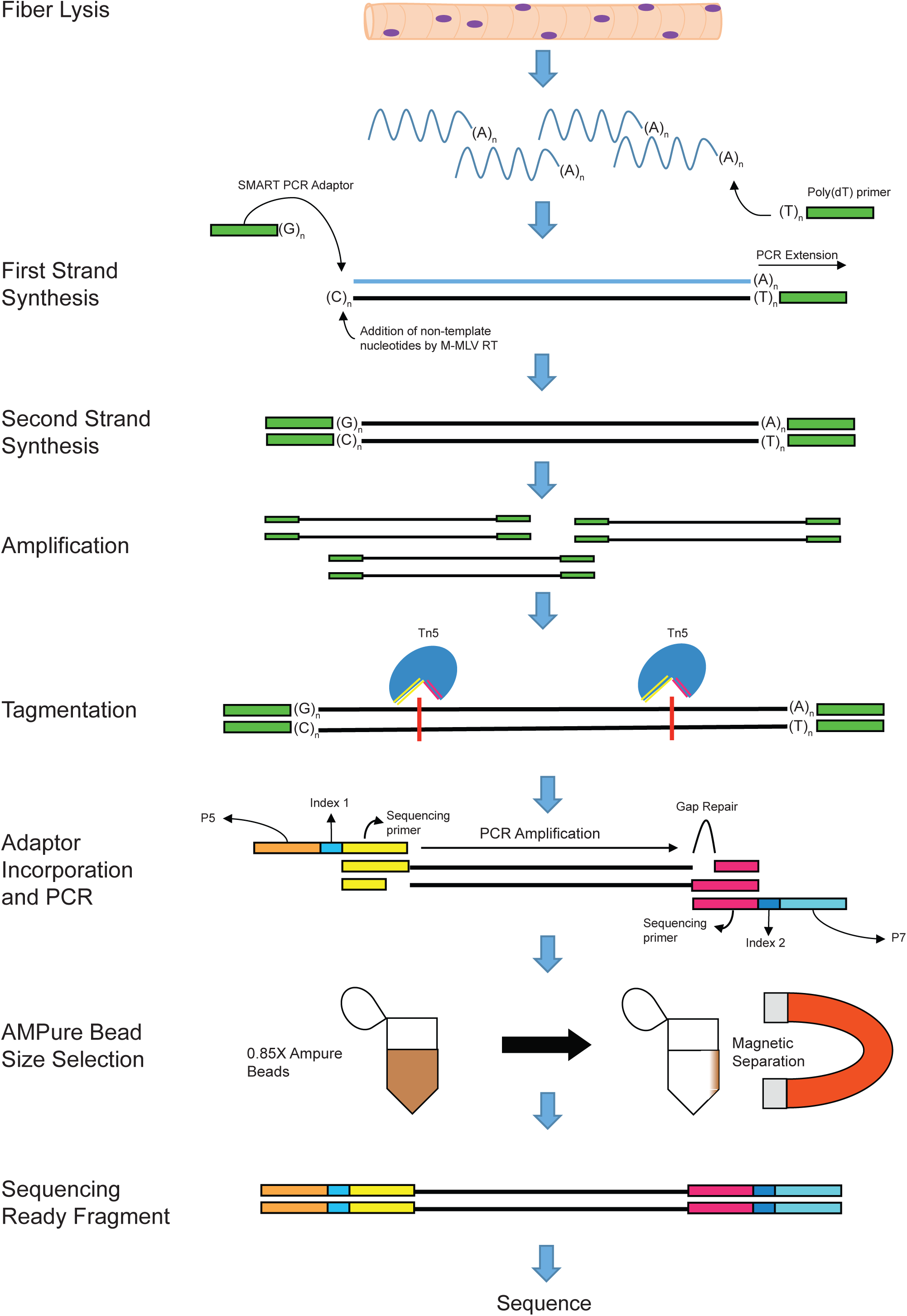
SMART technology and incorporation of Illumina adaptors to the fiber mRNA. Schematic displaying the steps and biochemical reactions involved in the generation of sequence ready cDNA fragments

We describe a robust method to extract RNA from a single myofiber followed by generation of sequencing ready libraries and whole transcriptome analysis. Using this technique, we first determined the genes that are found in the whole muscle that are not produced by the myofiber and are instead produced by non-myogenic cell types. To demonstrate the effectiveness of this technique, we next went on to analyze the differences in the whole transcriptome between myofibers isolated from young and old mice. smfRNA-Seq proved to be a useful tool in determining gene expression changes that occur in myofibers between different conditions.

## RESULTS

### Isolation of high quality mRNA from single myofibers

One of the intentions of our novel method is to give researchers the ability to sequence RNA from a single myofiber without the confounding presence of other cell types. A potential source of unwanted signal in the sequencing of myofiber RNA is the presence of muscle stem cells, also known as satellite cells, that are physically associated with the fibers. Using our method, we have found that satellite cells can be almost completely removed from the fibers with the addition of trypsin to the collagenase digestion buffer at a final concentration of 0.25%. This process does not damage the fiber itself, with a proper EDL isolation yielding over 200 myofibers per mouse (Figure 2A). However, this buffer effectively strips the myofibers of their satellite cells as shown by the reduction in the number of PAX7^+^ cells per fiber (Figure 2B-D).

**Figure 2:**
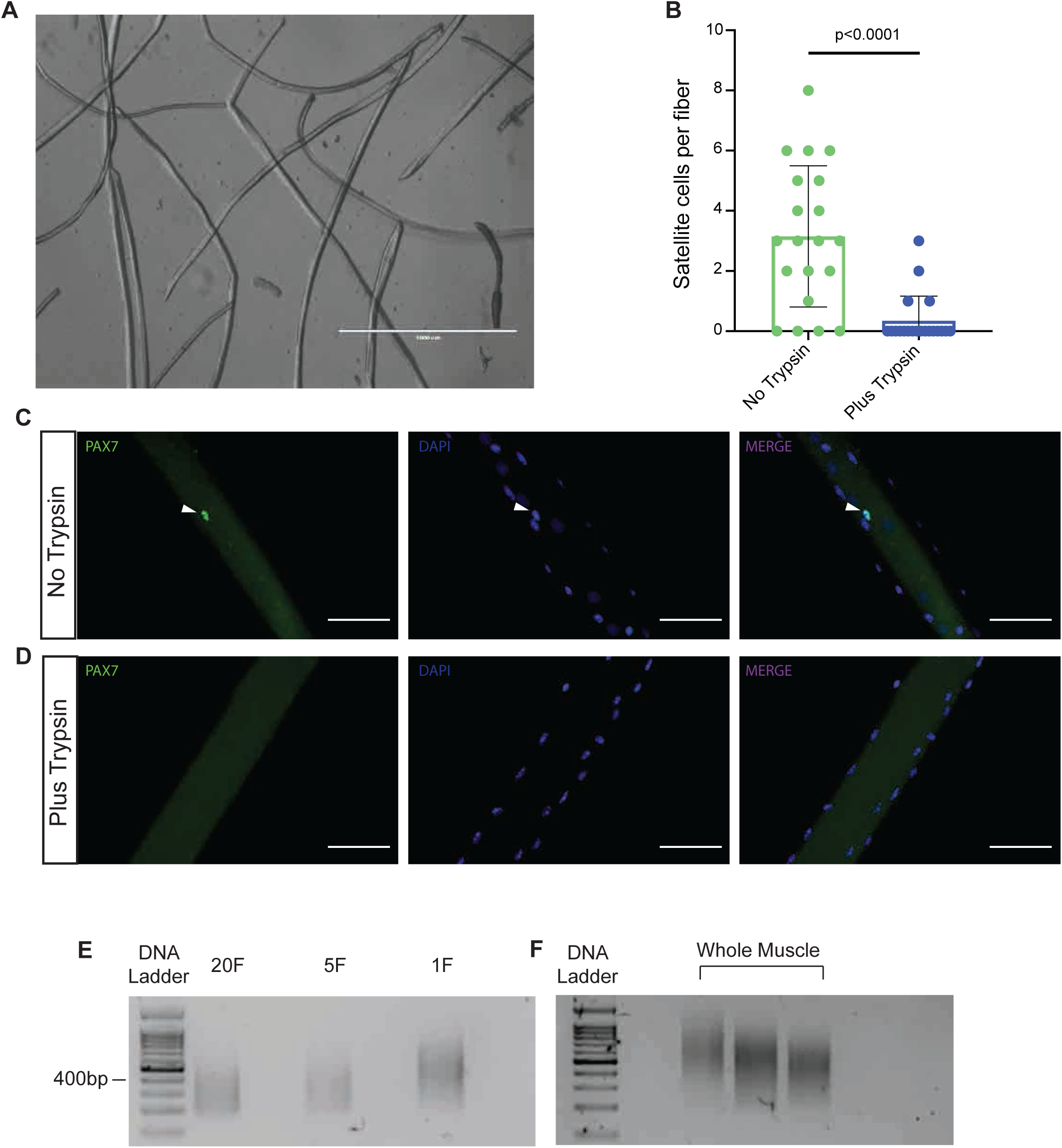
Isolation of total RNA from a single myofiber. (A) Isolated myofibers after a properly performed dissection and digestion. (B) Counts of the number of satellite cells per fiber in myofibers treated with and without trypsin. (C) Representative picture of a myofiber stained for PAX7 and counterstained with DAPI after no treatment with trypsin. (D) Representative picture of a myofiber stained for PAX7 and counterstained with DAPI after treatment with trypsin. (E) Representative image of cDNA from 20 fibers (20F), 5 fibers (5F), and single fiber (1F) on an agarose gel after library preparation and size selection. (F) Representative picture of cDNA from whole muscle on an agarose gel after library preparation and size selection.

Since muscle fibers are very tough and do not readily breakdown under normal lysing conditions, extracting the RNA from a single myofiber can prove challenging. With whole muscle, a method to overcome this is by freezing the muscle in liquid nitrogen and grinding it into a powder with mortar and pestle(10). However, this method cannot realistically be done to a single fiber while still collecting all of the RNA. Therefore, we lysed the fiber with lysis buffer in RNAse free water, utilizing osmotic pressure and gentle pipetting to break down the fiber and retrieve the intact RNA. This method proved effective as more than an adequate amount of RNA was recovered, even from a single myofiber, for use with SMART-Seq technology (Table 1).

**Table 1:** Total cDNA after SMART cDNA synthesis and amplification, and after incorporation of Illumina adaptors and size selection.

| **Samples** | **Total cDNA (ng) after SMART reaction and amplification** | **Total cDNA (ng) after adding Illumina adapters and size selection** | **Average fragment size after size selection (bp)** |
| --- | --- | --- | --- |
| Single Fiber 1 | 36.38 | 98.25 | 313 |
| Single Fiber 2 | 130.56 | 66.9 | 355 |
| Single Fiber 3 | 89.76 | 64.5 | 329 |
| Five Fibers 1 | 22.78 | 61.2 | 351 |
| Five Fibers 2 | 48.62 | 161.85 | 382 |
| Five Fibers 3 | 40.97 | 54.15 | 382 |
| Twenty Fibers 1 | 24.65 | 46.8 | 486 |
| Twenty Fibers 2 | 36.89 | 66.75 | 483 |
| Twenty Fibers 3 | 29.07 | 86.55 | 484 |
| Whole Muscle 1 | 3.74 | 34.65 | 617 |
| Whole Muscle 2 | 4.76 | 63.45 | 528 |
| Whole Muscle 3 | 5.1 | 53.25 | 456 |

From the extracted RNA, we successfully generated sequencing-ready cDNA libraries using the DNA SMART-Seq HT kit (Takara Biosciences) in combination with the Nextera XT DNA Library Preparation Kit (Illumina). Final single myofiber sequencing-ready libraries were of adequate quantity and of ideal size for sequencing, with fragments being of comparable average size as those generated from traditional whole muscle RNA-Seq libraries (Figure 2E-F). This implies that our fiber RNA extraction procedure generated high quality starting material that was compatible with low-input library preparation technologies.

### Comparative analysis of whole muscle and single myofiber RNA-Sequencing

By evaluating the number of unique reads obtained from smfRNA-Seq samples, we found that sequencing depth of single myofiber libraries was comparable to whole muscle RNA sequencing (Figure 3A). We obtained an average of 24 million unique reads from single myofiber samples when multiplexing 12 samples per lane on a NextSeq500, and an average of 35 million unique reads when multiplexing 10 samples per lane. This corresponds to an average overall alignment of 82.42%, with an average unique alignment percentage of 64.47% (Table S3). The overall expression profile of the single myofibers is also very similar to that of the whole muscle, indicating a similar high resolution (Figure 3A). When performing Principal Component Analysis (PCA), single fiber samples cluster together on both axes, but away from the whole muscle on the PCA2 axis (Figure 3B). To compare the efficiency of single cell technologies with increasing input, we also generated RNA-Seq libraries from five and twenty myofibers. When the number of fibers is increased, samples become less similar to both the whole muscle and the single myofiber samples, with higher variation between technical replicates (Figure 3A,B). This is most likely due to the excessive quantity of sample and could be resolved by scaling up the protocol.

**Figure 3:**
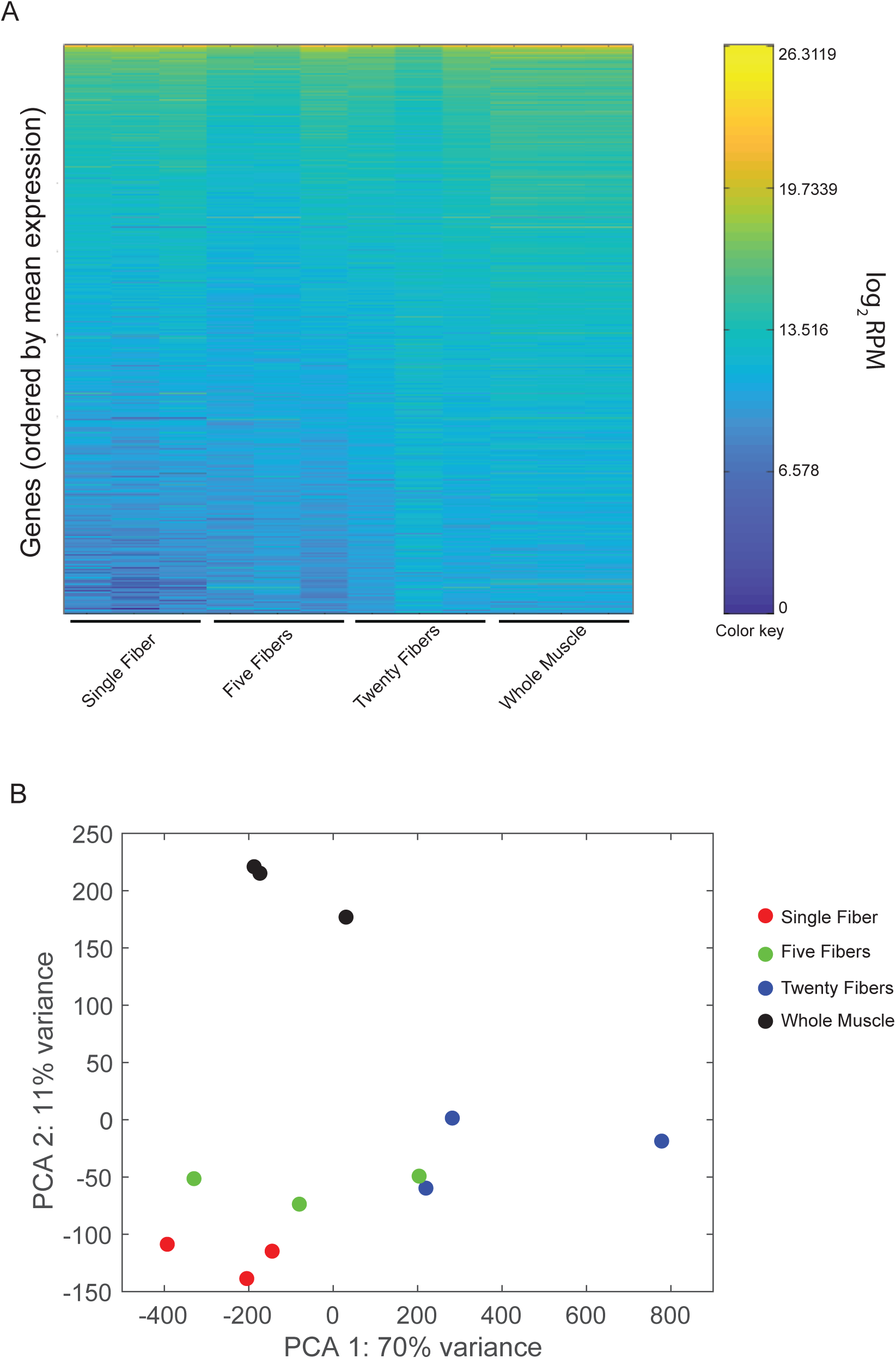
Comparative analysis of whole transcriptome from single myofibers and whole muscle. (A) Heatmap of gene expression in single fibers, groups of five fibers, groups of twenty fibers, and whole muscle, each with three replicates. Colors represent mean gene expression within each sample, from highest expression (yellow), to lowest expression (dark blue). Genes are ordered from top to bottom by their average expression across all samples. (B) Projection of samples along first two principal components found by PCA applied to log reads-per-million gene expression.

When looking more in depth at individual genes we see many similarities between the single myofiber and the whole muscle, but also crucial differences. Of particular note, we see that muscle specific genes have similar numbers of reads between the single fiber and the whole muscle samples (Figure 4A-C). The *Myh* cluster codes for a variety of myosin heavy chain proteins (MyHC), which are the motor proteins of muscle whose various isoforms are the basis of the different fiber types(7, 11). The similar expression between the single fiber and the whole muscle conclusively shows that the RNA sequenced came from a myofiber alone. (Figure 4A). For further confirmation, we also display *Ckm*, the muscle specific creatine kinase(12) and *ACTA1*, which codes for skeletal muscle alpha actin(13). These genes have the same pattern as is seen with *Myh* (Figure 4B,C).

**Figure 4:**
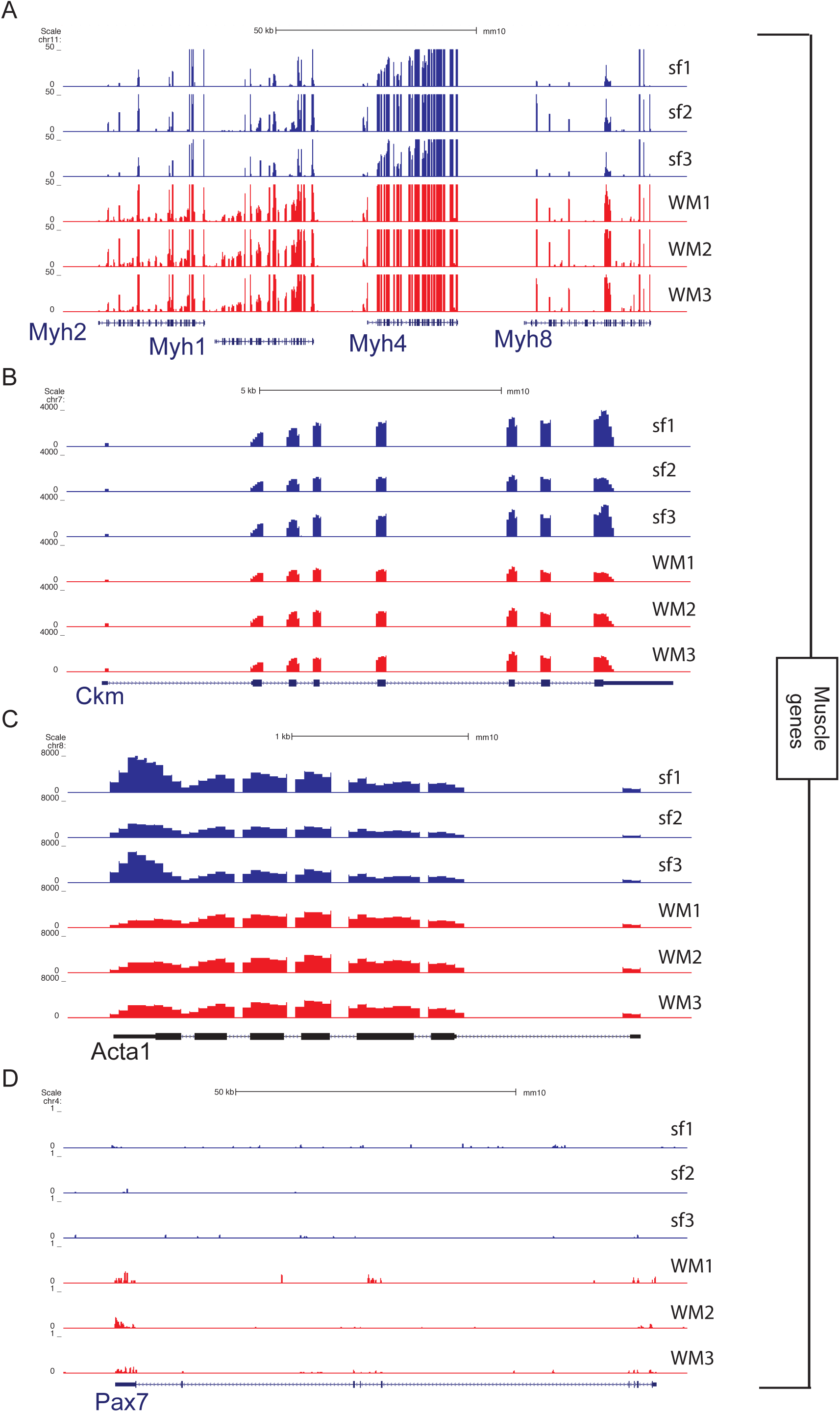
UCSC snapshots showing expression of myogenic genes in single myofibers and whole muscle. (A) Part of the myosin heavy chain (*Myh*) gene cluster located on chromosome 11. (B) The muscle creatine kinase (*Ckm*). (C) Actin alpha 1 (*Acta1*) gene. (D) Paired Box 7 (*Pax7*) gene expressed in the associated satellite cells.

Differences between the single myofiber and whole muscle transcriptomes arise from non-muscle cells. Using smfRNA-Seq, we are capable of completely removing these unwanted cell types to sequence only the myofiber. To demonstrate the removal of cell populations other than the myofiber, we looked at cell specific genes of a variety of different muscle-resident cell types. We first looked at the expression of the satellite cell marker *Pax7,* and see that there is no expression of *Pax7* in the single fiber transcriptome despite its presence in whole muscle samples (Figure 4D). We also analyzed markers for fibroblasts, namely *Col1a1* and *Thy1(14,15)*. As expected for these genes, they are expressed in the whole muscle, but not expressed in the single fiber (Figure 5A,B). Endothelial cells are also depleted in single fiber samples, since their markers *Kdr* and *PECAM1* are expressed in whole muscle, but not in the single fiber(16, 17)(Figure 5C,D). We also analyzed *Retn* to identify the presence of adipocytes(18), *Cd34* for hematopoietic cells(19), *Ly6a* as a marker for FAPs(2), and *ADGRE1* for macrophages(20) (Figure 5E-J). As expected, in the single fiber transcriptome there is no expression for any of these genes, all of which were present in the whole muscle. These results clearly demonstrate the purity of smfRNA-Seq in sequencing only the myofiber without confounding cell types.

**Figure 5:**
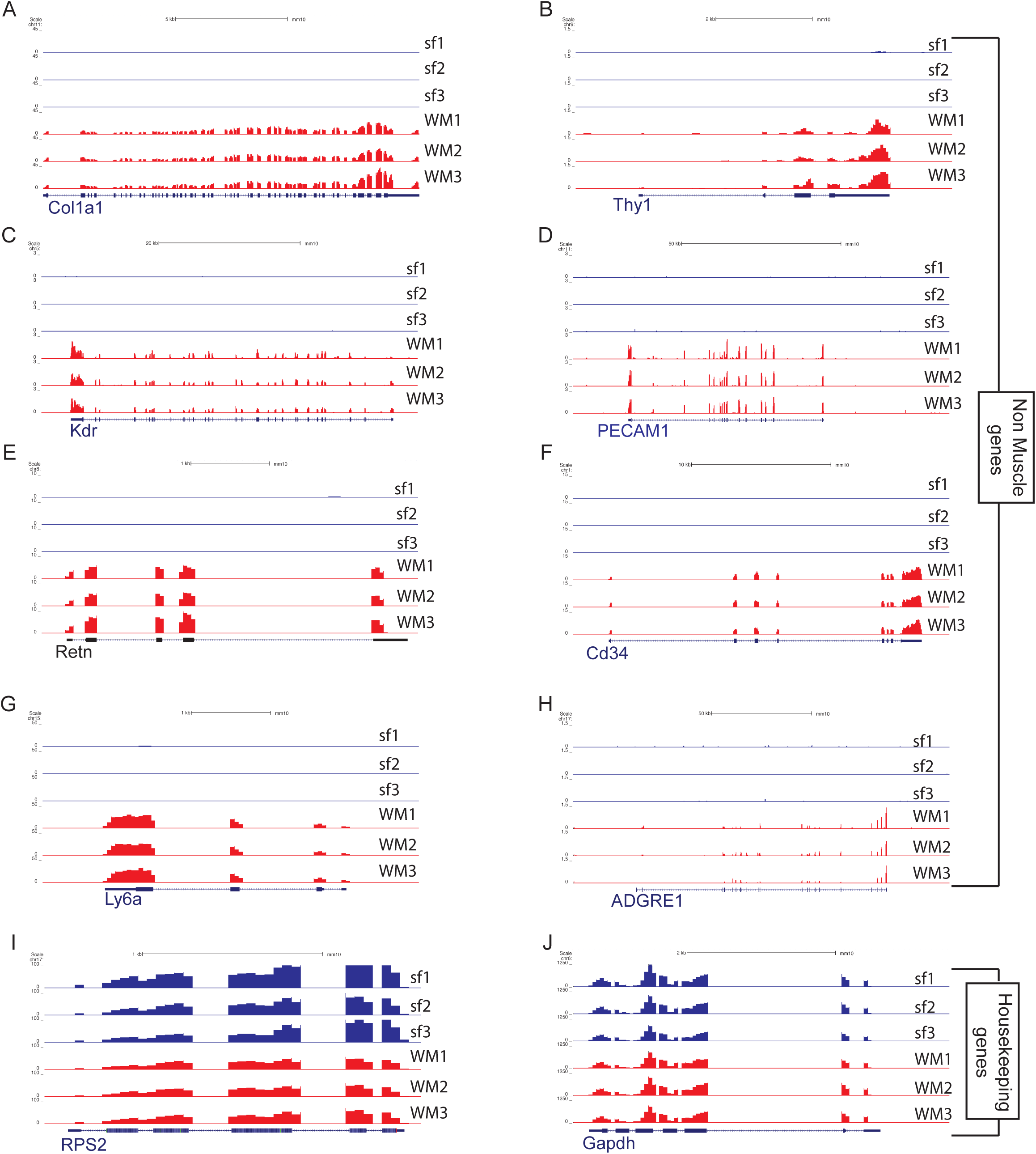
UCSC snapshots showing expression of non-myogenic genes between single myofibers and whole muscle. (A) Genes collagen type 1 alpha 1 chain (*Col1a1*) and (B) CD90 (*Thy1*) expressed in fibroblasts. (C) The genes kinase insert domain receptor (*Kdr*) and (D) CD31 (*PECAM1*) as markers for endothelial cells. (E) Resistin (*Retn*) as a marker for adipocytes. (F) *Cd34* as a marker for hematopoietic cells. (G) *Ly6a* to detect the presence of fibro/adipogenic progenitors (FAPs). (H) Adhesion G protein-coupled receptor E1 (*ADGRE1*) gene for macrophages. (I) Housekeeping genes *RPS2* and (J) *Gapdh*

From this data, we performed a GO term analysis of genes that are expressed solely in the WM samples, defined as having an RPM value of at least 10 in WM and 0 in SF, indicating that they originate from non-myogenic cells. Using these criteria, we identified 445 genes that were solely expressed in the WM muscle samples (Table S1). Some of the more upregulated genes were involved in the formation of the extracellular matrix or immunity (Table S1).

We also attempted to identify genes that were exclusively produced in the myofiber. We defined these genes as being more highly expressed in the SF samples, with a q value < 0.01, being expressed at more than 10 RPM and being expressed at least 10 RPM more than in the WM samples. We identified 622 genes that matched these criteria and performed a Gene Ontology (GO) analysis (Table S2). From this, we see that some of the top pathways associated with these genes are Ribosomal proteins and the Respiratory chain. However, this is not an exhaustive list, due to the difficulty in identifying genes expressed solely in myofibers when the WM samples are composed primarily of myofibers. For example, *Myh4,* which is expressed exclusively in muscle fibers, does not pass the criteria laid down and is not part of the list of genes expressed solely in myofibers. On the other hand, dystrophin (*DMD*), another myofiber specific gene does pass our criteria.

### Age effect on the transcriptome of single myofibers

In order to verify the utility of our new method to analyze variation in the myofiber transcriptome under different conditions, we performed smfRNA-Seq on EDL myofibers isolated from young (1 and 3 months-old) and aged (19 months-old) mice.

Using our technique, we see clear differences in the transcriptome of young and old myofibers. In the old myofibers we see the deregulation of a number of genes, with 181 genes being significantly (padj <0.05) differentially expressed in old compared to young myofibers (Figure 6A,B). Principal Component Analysis (PCA) demonstrates how young myofibers, isolated from 1 month old mice, cluster together away from the old myofibers on the PC2 axis (Figure 6E). Furthermore, the myofibers isolated from 3 month-old mice begin to resemble the 19 month-old myofibers, while also maintaining similarities with the young 1 month-old myofibers, demonstrating the gradual change in transcriptome of the myofibers as the mouse ages (Figure 6E). GO Term analysis of the genes with the highest variation between young and old myofibers shows that deregulated pathways in aging include the transport of small molecules such as salts and metal ions, and the synthesis of collagens. Additional examples of genes significantly deregulated in aging include *Actc1*, *Myl1*, *Dkk3*, *Atf3*, *H19*, *Nos1* and *Ndn* (Figure 7A-H). These are genes involved in skeletal muscle structure, metabolism, growth and maintenance of homeostasis.

**Figure 6:**
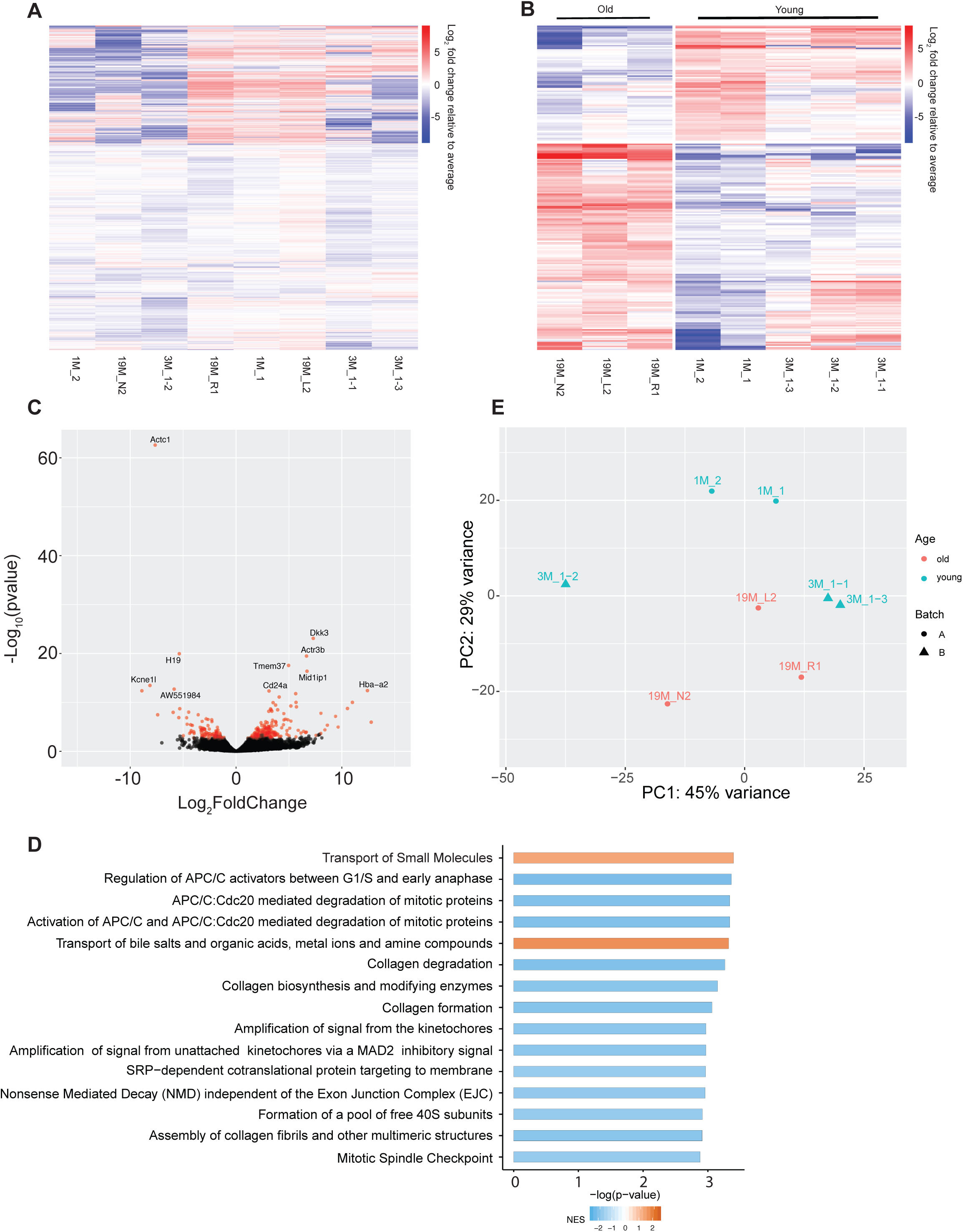
Comparative analysis of whole transcriptome between single myofibers from young and old mice. (A) Heatmap of all genes expressed in old and young and (B) heatmap of differentially expressed genes between the old and young myofibers. Colors indicate Log_2_ fold change relative to average per gene with red indicating a higher expression and blue as a lower expression between young (1 and 3 months) and old (19 months) myofibers. (C) Volcano plot of the differentially expressed genes between young and old myofibers. Points in red indicate there is a significant difference between the two groups. (D) Go term analysis of the top 15 differentially regulated pathways. (E) Projection of samples along first two principal components found by PCA applied to log reads-per-million gene expression of young (1 and 3 months) and old (19 months) myofibers.

**Figure 7:**
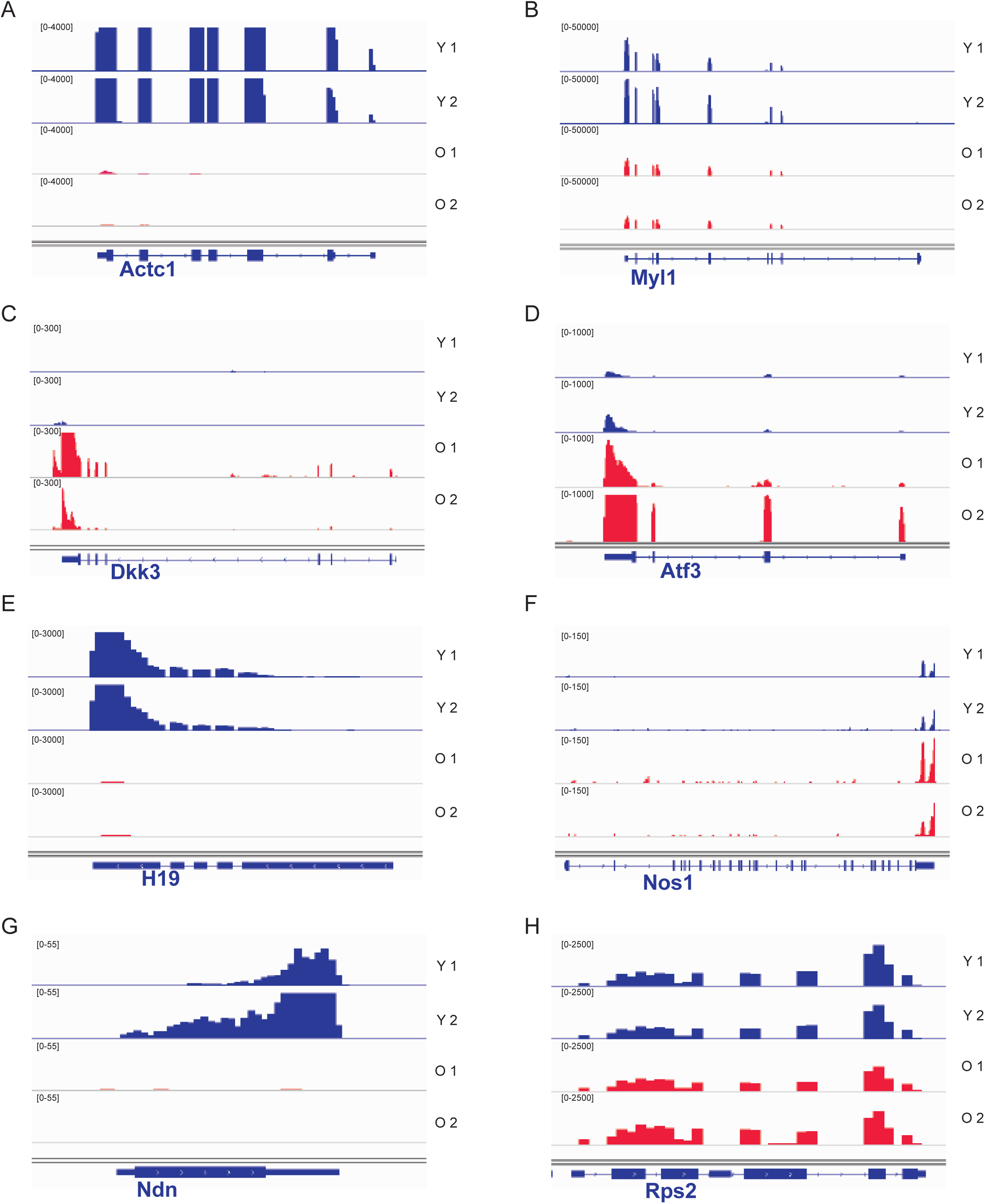
IGV snapshots showing the expression of selected differentially expressed genes between young and old myofibers. Young myofiber tracks are labeled in blue and old are in red. (A) Actin alpha cardiac muscle 1 (*Actc1*). (B) Myosin light chain 1 (*Myl1*). (C) dickkopf WNT signaling pathway inhibitor 3 (*Dkk3*). (D) Activating transcription factor 3 (*Atf3*). (E) H19, imprinted maternally expressed transcript (*H19*). (F) Nitric oxide synthase 1 (*Nos1*). (G) Necdin (*Ndn*). (H) housekeeping gene ribosomal protein S2 (*Rps2*).

## DISCUSSION

smfRNA-Seq is a powerful new technique that allows high resolution whole transcriptome sequencing of a single myofiber. Here, we have demonstrated that RNA can be successfully isolated from a single myofiber at a suitable concentration and quality to be used with SMART-Seq technology, generating a high-quality cDNA library for sequencing.

What distinguishes smfRNA-Seq from single cell RNA-Seq is that sequencing depth is comparable to large-scale bulk RNA-Seq (Figure 3). However, as shown by PCA, single myofiber samples are very similar to whole muscle samples with regards to PCA1, but dissimilar with regards to PCA2. By analyzing individual genes, we can interpret the similarities as being due to muscle specific genes, while the lack of other cell types allows the single fibers to form a distinct cluster away from the whole muscle. Indeed, comparison of single myofiber to whole muscle samples identified a list of genes expressed only in the whole muscle, which are likely responsible for this distinction (Table S1). This dissociation of the myofiber transcriptome from the transcriptome of other muscle-resident cell types will be of critical importance to tease out the contribution of the myofiber when a gene of interest is expressed in multiple cell types, or when a treatment condition has a whole muscle effect.

One condition that leads to changes in the numbers and in the gene expression profile of various muscle-resident cell types is aging (21, 22). Due to the changes occurring in multiple cell types, including immune cells (23, 24), FAPs (25), and satellite cells(26, 27), whole muscle RNA-Seq would be inappropriate to distinguish age-induced changes in myofibers alone. Using smfRNA-Seq, we were able to uncover differences in the transcriptome of old and young myofibers, without any added noise from age-related alterations in other cell types. Among the genes deregulated in aging myofibers are genes involved in muscle growth and structure, suggesting that some of these altered genes may be responsible for age-related muscle atrophy, also called sarcopenia. Of note, we see that *Actc1* and *Myl1* are greatly downregulated in old compared to young myofibers (Figure 7A,B). Despite not reaching significance, there is also a moderate decrease in *Acta1* with a log_2_fold change of -0.84. Since actins and myosins play a role in myofiber structure, force and contractile properties, we suspect that the downregulation of these genes may play a role in the weakening in contractile strength in aged muscles (28, 29). A gene that is significantly upregulated in aged myofibers is *Dkk3,* which encodes a secreted inhibitor of WNT signaling (Figure 7C) (30). This finding is in line with a previous study that demonstrated that increased Dkk3 in aged muscle tissue leads to muscle atrophy through autocrine signaling(30), while also confirming that the myofiber is the source of this secreted factor. There is also a downregulation in *Ndn* in the old myofibers (Figure 7G). *Ndn* produces Necdin, which is important in muscle regeneration, as it promotes the differentiation of satellite cells, and Necdin knockout mice have a significant impairment in muscle regeneration(31). Necdin was also shown to protect skeletal muscle from tumor-induced muscle wasting, also called cachexia, in part by binding to and inhibiting P53 (32). It would be of interest to see whether Necdin decrease in aging myofibers is also a cause of age-induced muscle atrophy. In addition to genes involved in muscle structure and growth, our data also shows that multiple genes involved in key metabolic processes and maintenance of homeostasis are also deregulated in old compared to young myofibers. For example, we detect a loss in the expression of the long noncoding RNA *H19* (Figure 7E). While the role of H19 during myogenesis has been studied (33), namely in controlling myoblast proliferation, its role in fully differentiated myotubes is still unclear. However, H19 has also been shown to be involved in glucose metabolism and consequently, insulin sensitivity (34, 35). We speculate that H19 reduction in old myofibers may therefore be a contributor to metabolic dysfunction, which is a common feature of aging (35).

In old myofibers, we also see an upregulation of *Atf3*, which is a stress factor that is known to be upregulated in muscle after a disturbance in homeostasis, such as after exercise (36) (Figure 7D). Interestingly, Atf3 was shown to dampen the expression of pro-inflammatory cytokines after exercise (37). We speculate that its upregulation in aging myofibers may occur in response to an age-related disturbance in homeostasis, such as the stress induced by chronic low-grade inflammation(38). Another gene that is upregulated in aging is *Nos1* which produces nitric oxide (NO) and plays a role in skeletal muscle metabolism (Figure 7F). Notably, Nos1 is known to increase glucose uptake and inhibit mitochondrial respiration(39), which may be a contributor to age-related metabolic dysfunctions. In addition to these top deregulated genes, GO Term analysis revealed that genes involved in collagen synthesis, assembly and degradation are all perturbed in aging myofibers. Since collagens are important components of the muscle extracellular matrix, our data suggests that old myofibers may contribute to changes in ECM stiffness (40, 41).

Altogether, we were able to demonstrate the effectiveness of smfRNA-Seq by sequencing the transcriptome of single myofibers from young and old mice. This method allowed us to conclude that aging does have an effect on myofibers at the level of the transcriptome, without the confounding signal from non-myogenic cell types. This comparative analysis proves that smfRNA-Seq can thus be used to study other developmental or muscle-wasting disorders, such as muscular dystrophy. Additionally, smfRNA-Seq has the potential to be adapted to a larger scale in order to perform high-throughput analysis of numerous myofibers, which will be important in the study of myofiber heterogeneity.

## MATERIALS AND METHODS

### Animal care

All procedures that were performed on animals were approved by the McGill University Animal Care Committee (UACC)

### Accession numbers

All gene expression data reported in this study is available through GEO accession number GSE135364 and GSE138591

### Commercial kits

The following commercial kits were used in this experiment.

SMART-Seq HT Kit (Takara Cat# 634437)

Nextera XT DNA Library Preparation Kit (Illumina Cat# FC-131-1024)

Nextera XT Index Kit (Illumina Cat# FC-131-1001)

### Buffers

Myofiber digestion buffer was prepared using 1000 U/mL of Collagenase from Clostridium histolyticum (Sigma Cat# C0130) in unsupplemented DMEM (Invitrogen Cat# 11995073)

Myofiber immunofluorescence blocking buffer is composed of 5% horse serum (Wisent Cat# 065-250), 2% Bovine Serum Albumin (Sigma Cat# A8022), 1% Triton-X_100_ (Sigma Cat# T9284), in PBS (Wisent Cat# 311-425-CL)

RNA extraction buffer is made using 19 μL of the 10X lysis buffer plus 1 μL of RNAse inhibitor from the SMART-Seq HT Kit. 1μL of the previously composed 10X lysis buffer is added to 9 μL of RNAse free water to make 1X lysis buffer

### Dissection of the Extensor Digitorum Longus (EDL)

The EDL was dissected from the hindlimb in the following manner: the skin of the hindlimb was removed by cutting around the ankle with a pair of scissors, and an incision was made along the ventral side of the leg. The epimysium around the Tibialis Anterior (TA) was removed and the tendon of the TA was cut at the ankle, while making sure to only cut the top tendon as the bottom tendon belongs to the EDL. Using a pair of forceps, the TA was gently peeled off the leg up to the knee and was then cut out as close to the knee as possible with a pair of scissors. To expose the EDL tendon at the knee, the biceps femoris was first removed with a pair of forceps. The EDL was removed by cutting from tendon to tendon with a pair of scissors.

### Myofiber isolation

The dissected EDL was placed in a 1.5 mL Eppendorf tube with 800 μL of myofiber digestion buffer. Trypsin was added to a final concentration of 0.25% to remove the associated satellite cells. The EDL was incubated at 37°C and 5% CO_2_ for at least one hour.

To disassociate the myofibers, we transferred the EDL to a 6-well plate with 2 mL of unsupplemented DMEM, that had previously been coated with 10% horse serum (HS) in DMEM for at least 30 minutes. The EDL was gently pipetted up and down with a large bore pipette coated in HS until no more myofibers could be retrieved.

### Immunofluorescence of myofibers

Briefly, freshly isolated myofibers were fixed at T_0_ using 4% paraformaldehyde in PBS for 5 minutes. They were washed 3 times with 0.1% Triton-X_100_ in PBS, then permeabilized with 0.1% Triton-X_100_ and 0.1M glycine in PBS for 15 minutes, and again washed 3 times with 0.1% Triton-X_100_ in PBS. Myofibers were blocked for 1 hour with blocking buffer, followed by incubation with a Pax7 hybridoma primary antibody (DSHB #AB_528428) at 1:100 dilution in blocking buffer overnight at 4°C. The next day, fibers were washed 3 times with 0.1% Triton-X_100_ in PBS, before incubating the Alexa Fluor 488 goat anti-mouse IgG1 secondary antibody (Invitrogen Cat# A21121) at a 1:400 dilution in blocking buffer for 1 hour. Fibers were washed 3 times with 0.1% Triton-X_100_ in PBS and mounted on a microscope slide with Prolong Gold Antifade Reagent with DAPI (Invitrogen Cat# P3695)

### Single Myofiber RNA extraction

Using a small bore glass pipette, coated with HS, myofibers were transferred to a 6-well plate with 2 mL of PBS to wash the fibers. A single myofiber was transferred, by using a coated small bore pipette, to a 0.2 mL PCR tube and the excess PBS was removed using a pipette. Next, we added 10 μL of lysis buffer, and gently pipetted the myofiber up and down for 3 minutes and then incubated the fiber on ice for 5 minutes while periodically vortexing and spinning down the sample.

The residual fiber pieces were removed by spinning down the sample and transferring the supernatant to a fresh PCR tube.

### Whole muscle RNA extraction

The whole muscle from the hindlimb of a mouse was dissected, frozen in liquid nitrogen and ground into a powder using a mortar and pestle. RNA from whole muscle was extracted using TRIzol reagents (Ambion #15596018).

### cDNA library preparation

cDNA was constructed using the DNA SMART-Seq HT kit and following the manufacturers recommendations. For a single fiber, we used 12 cycles of PCR amplification on the cDNA. The cDNA was then purified using AMPure XP beads (Beckman Coulter #A63880) at a 1:1 ratio and quantified using the Quant-iT PicoGreen dsDNA Assay kit (ThermoFisher #P11496)

For the addition of Illumina sequencing adapters, we used 250 pg of cDNA in 1.25 μL of water as a starting material and followed the directions provided with the Nextera XT DNA Library Preparation Kit, but reduced all quantities by 4x volume. The libraries were size selected using AMPure XP beads at 0.85X of the sample volume, to remove all fragments below 200 bp.

### Sequencing and gene expression analysis from a single fiber and whole muscle

The Illumina NextSeq 500 High Output Flow Cell was used for sequencing. The sequenced reads were then mapped to the mouse mm10 genome by using HISAT2(42), using an index downloaded from the HISAT2 website that jointly indexes the mm10 genome and the ENSEMBL transcriptome definition. (43).

### Differential expression analysis between old and young single myofiber from mouse

RNA-seq raw reads from young and old single myofiber were included in the analysis. Reads were mapped to the mouse genome assembly (mm10) using HISAT2 (42). The number of aligned reads per gene was obtained with HTSeq (44), using gene annotations from GENCODE M23 (45). Genes with an average read counts smaller than 10 were filtered out. Differentially expressed genes between young and old samples were identified using the R package DESeq2 (46). Log fold changes were calculated, and their associated P-values were corrected by independent hypothesis weighting (47).

### Gene Set Enrichment Analysis

To obtain the pathways affected between old and young mouse fiber, we used the implementation of Gene Set Enrichment Analysis of the fgsea R package (45). We used the Log fold changes calculated by DESeq2 to create a pre-ranked gene-list. The Reactome database was used as a reference for pathways (48).

## ACKNOWLEDGMENTS

This work was supported by grants from the Canadian Institute of Health Research (CIHR) [PJT-15087]; and a Natural Sciences and Engineering Council (NSERC) discovery grant to VDS.

## AUTHORS CONTRIBUTIONS

Conceived the idea VDS, performed experiments DMB and FL, analyzed data VDS, AHC, HN, TJP, DMB and FL; wrote the manuscript DMB, FL, VDS

---

**Table S1A:**
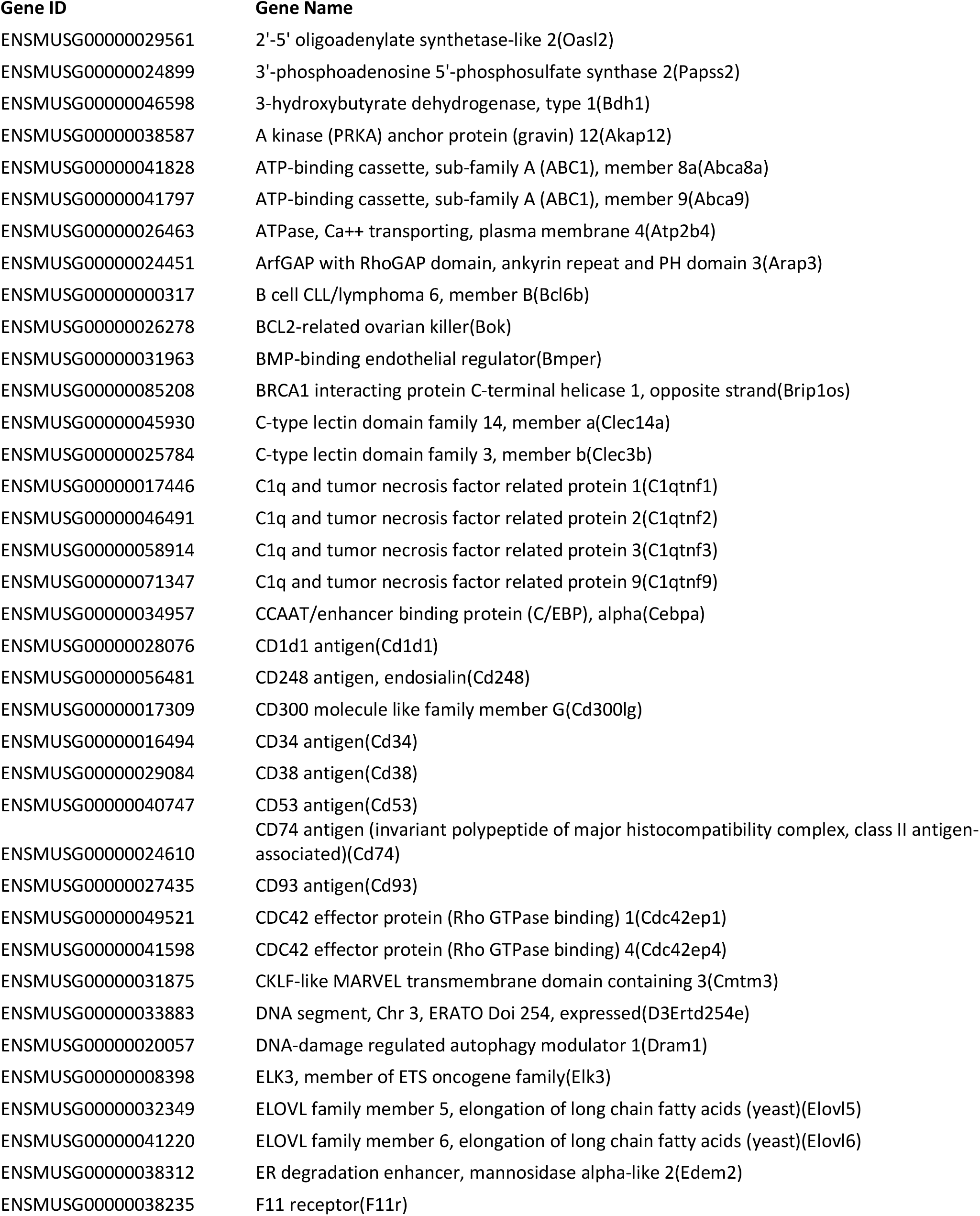
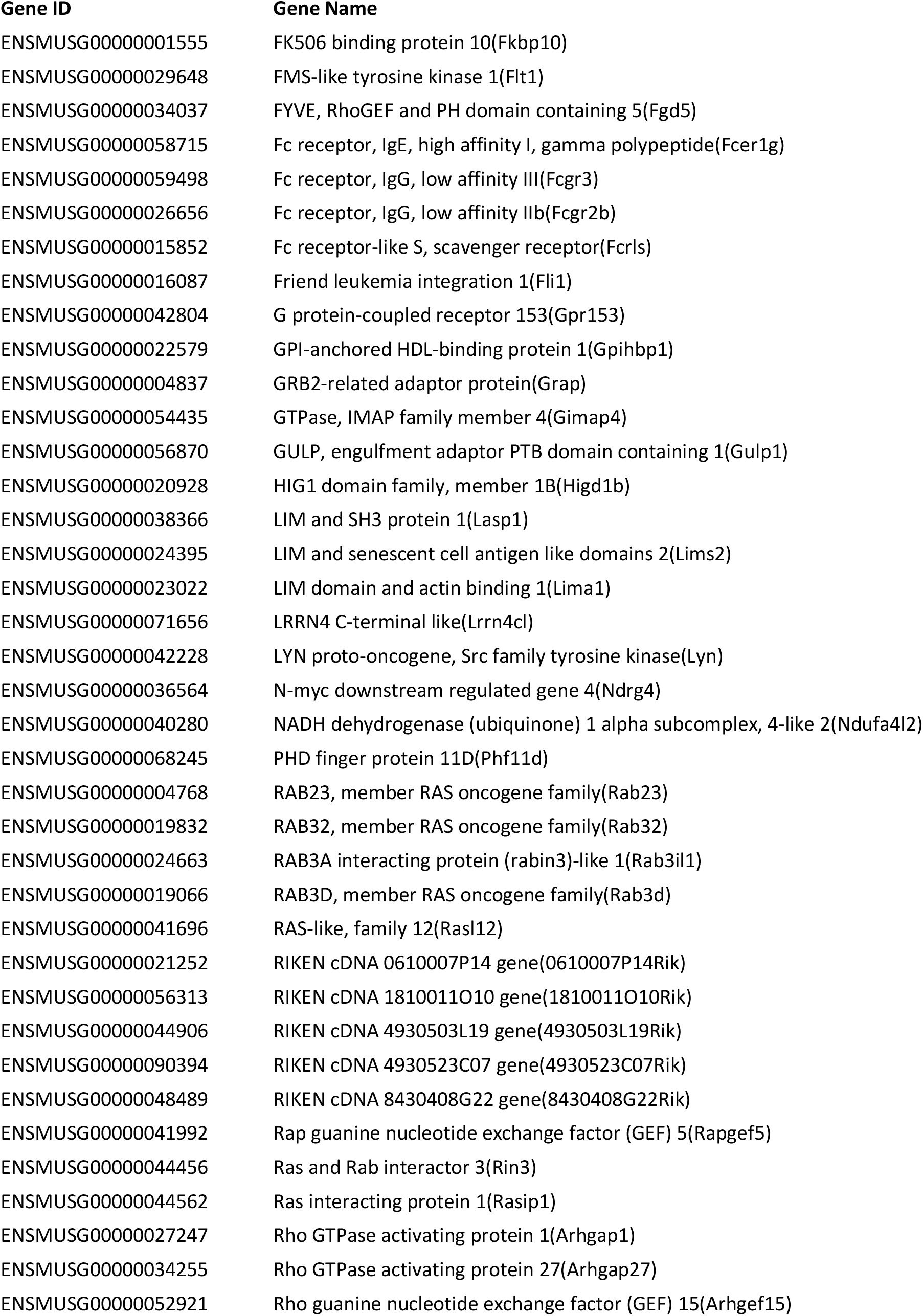
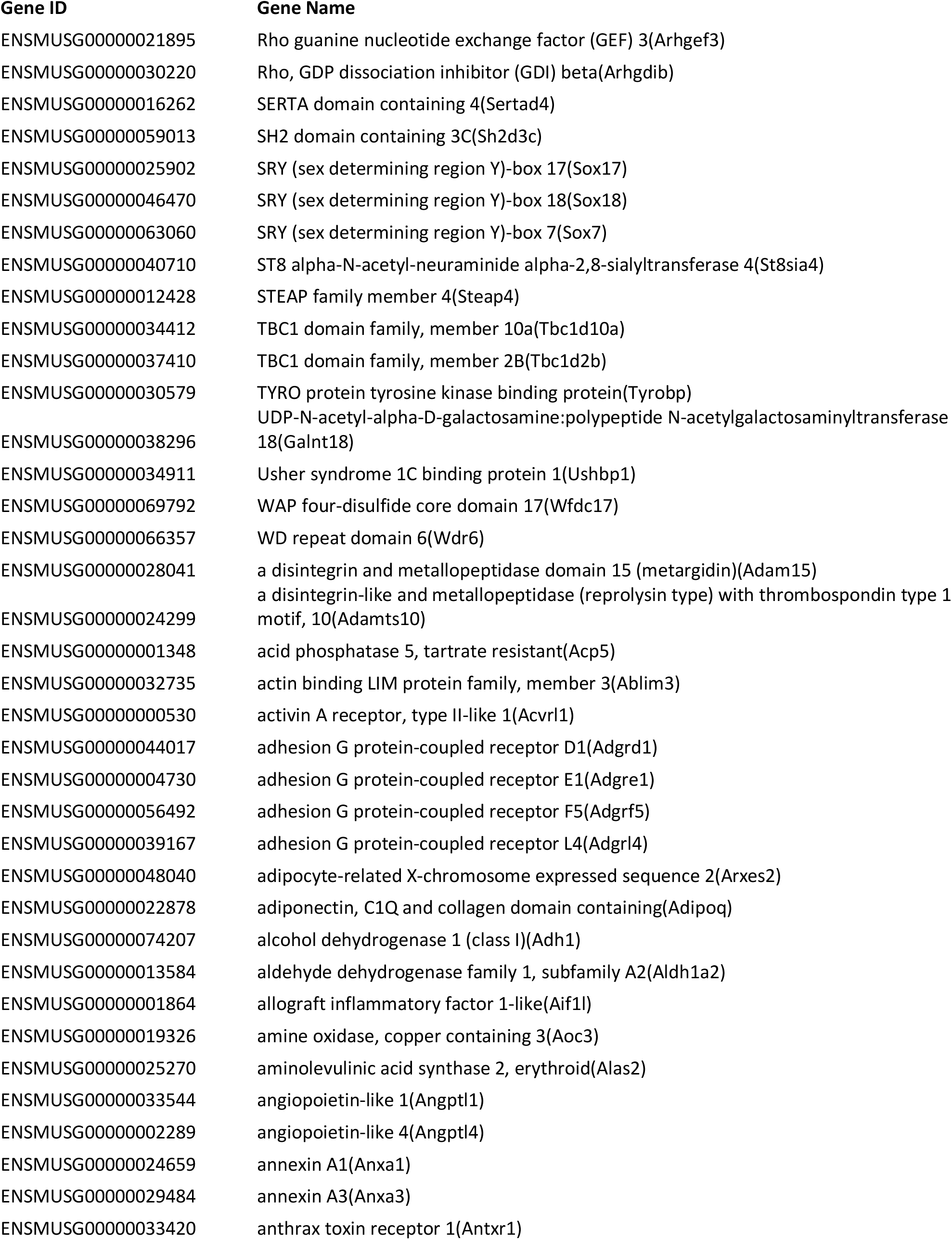
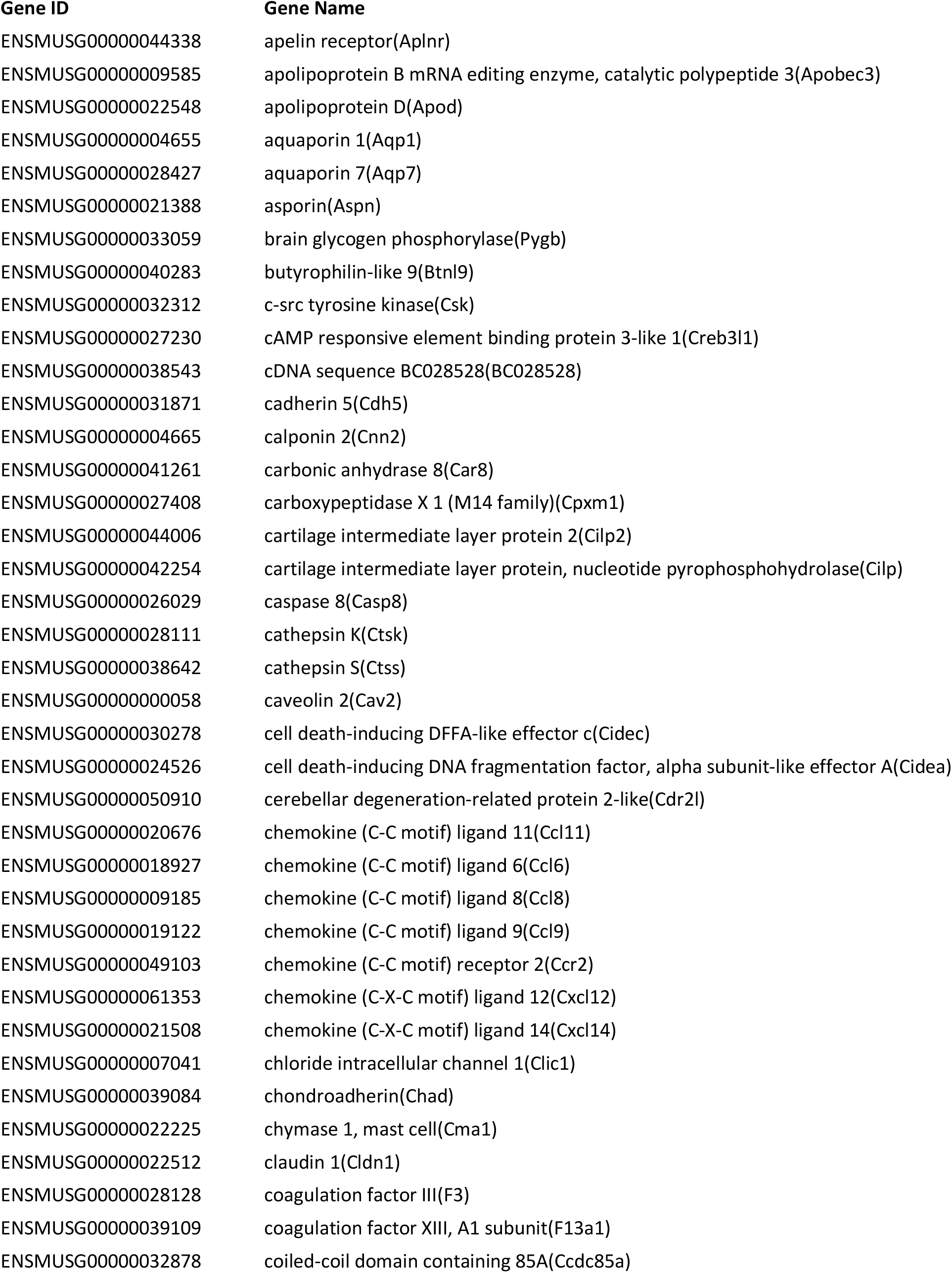
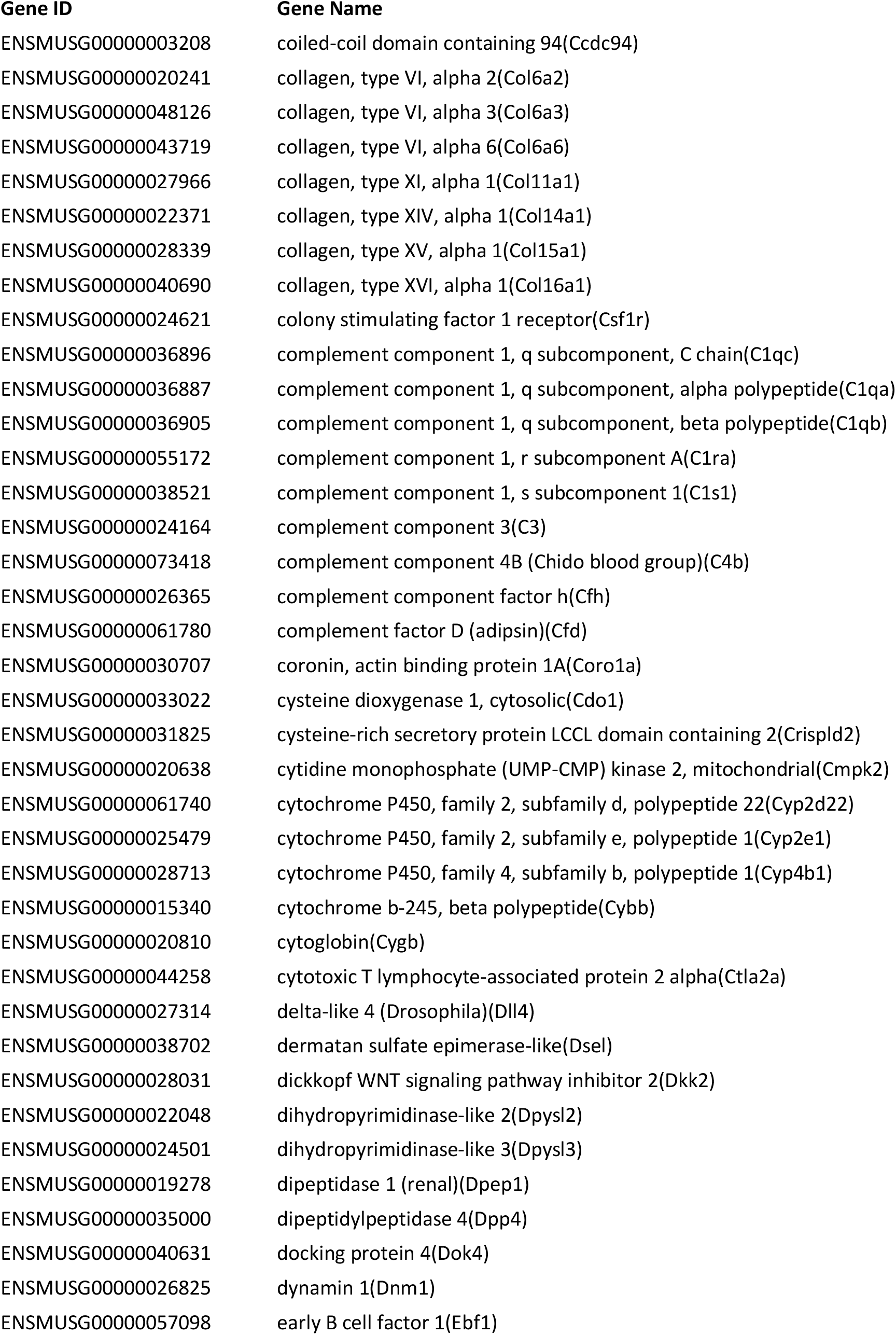
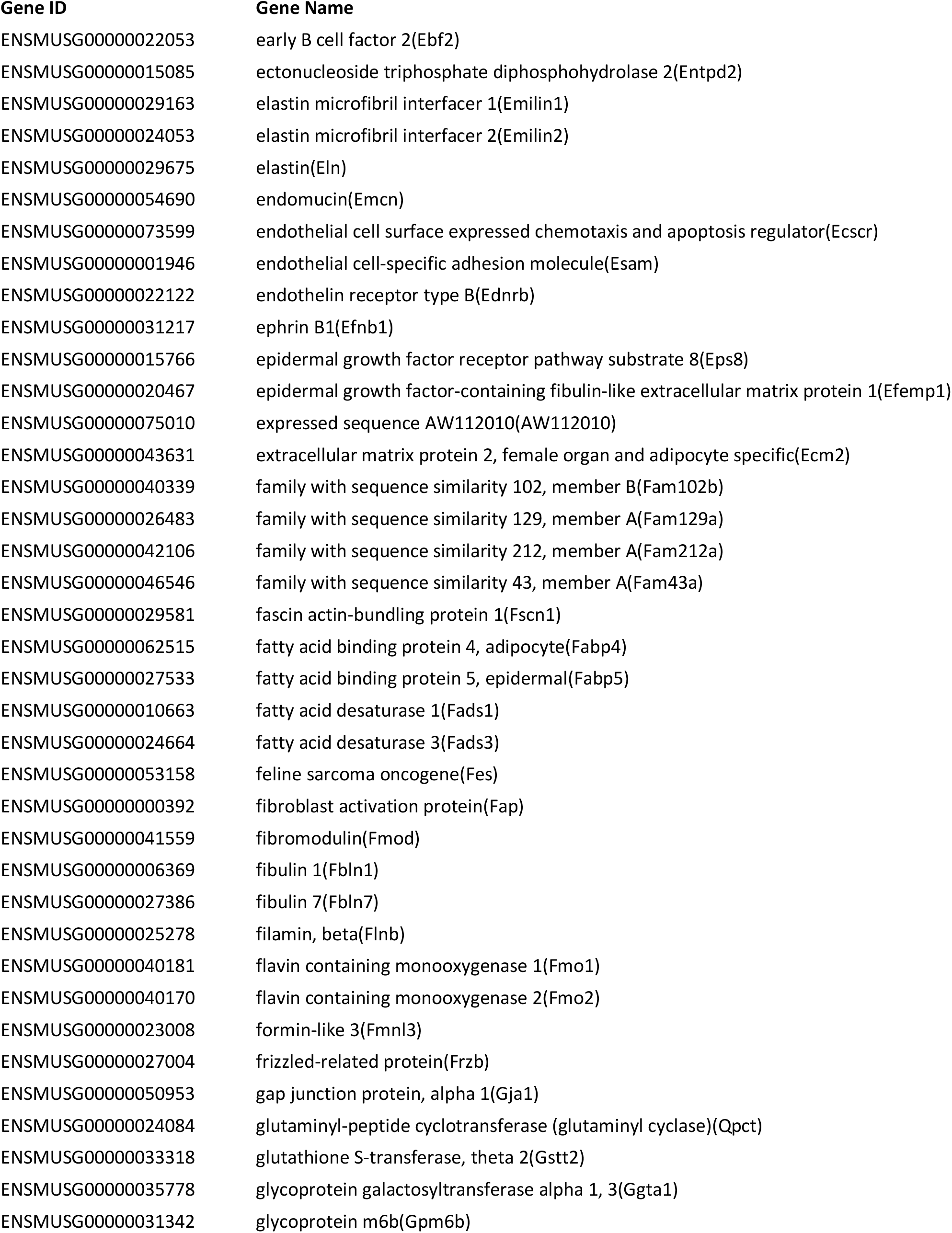
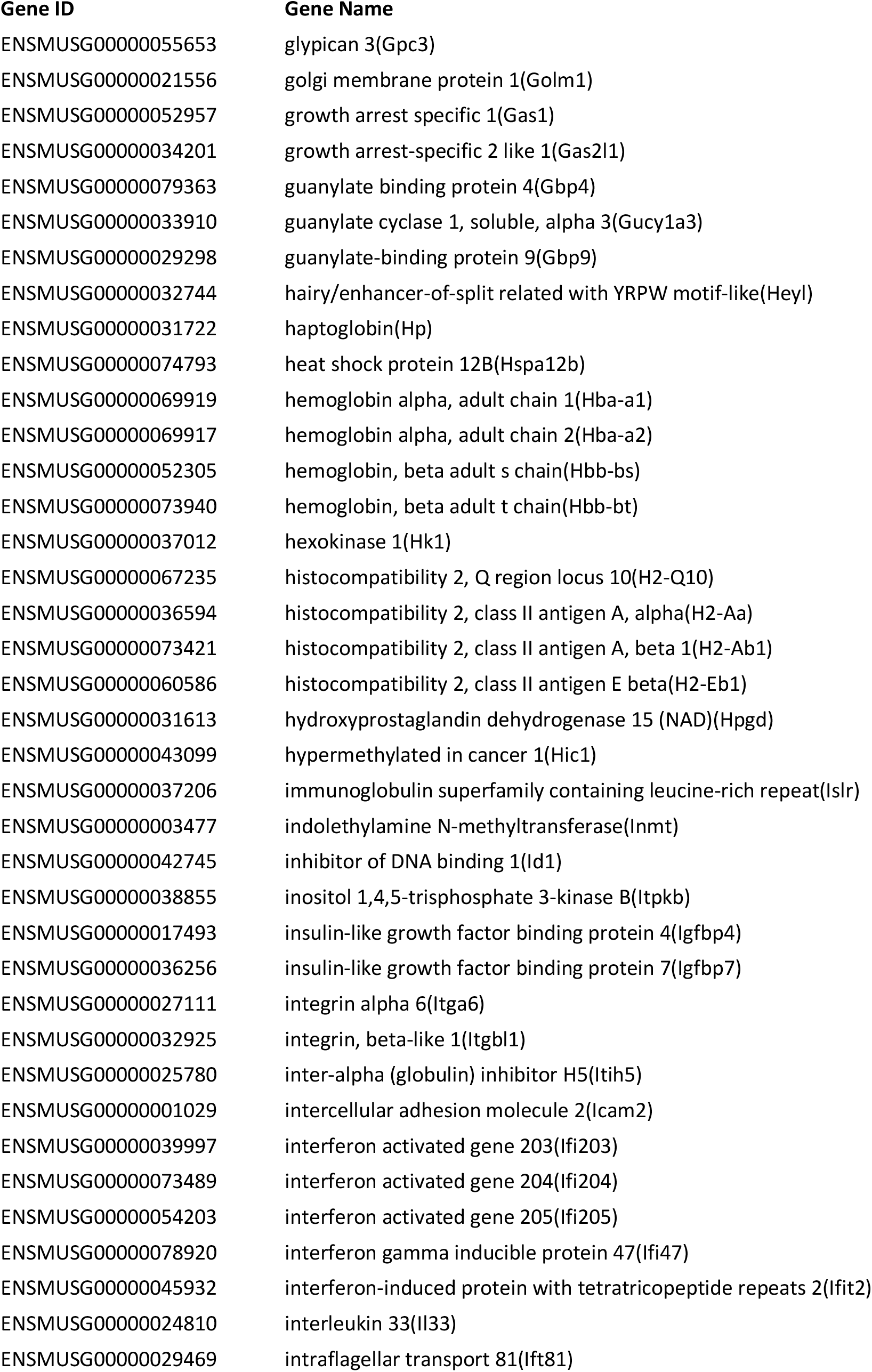
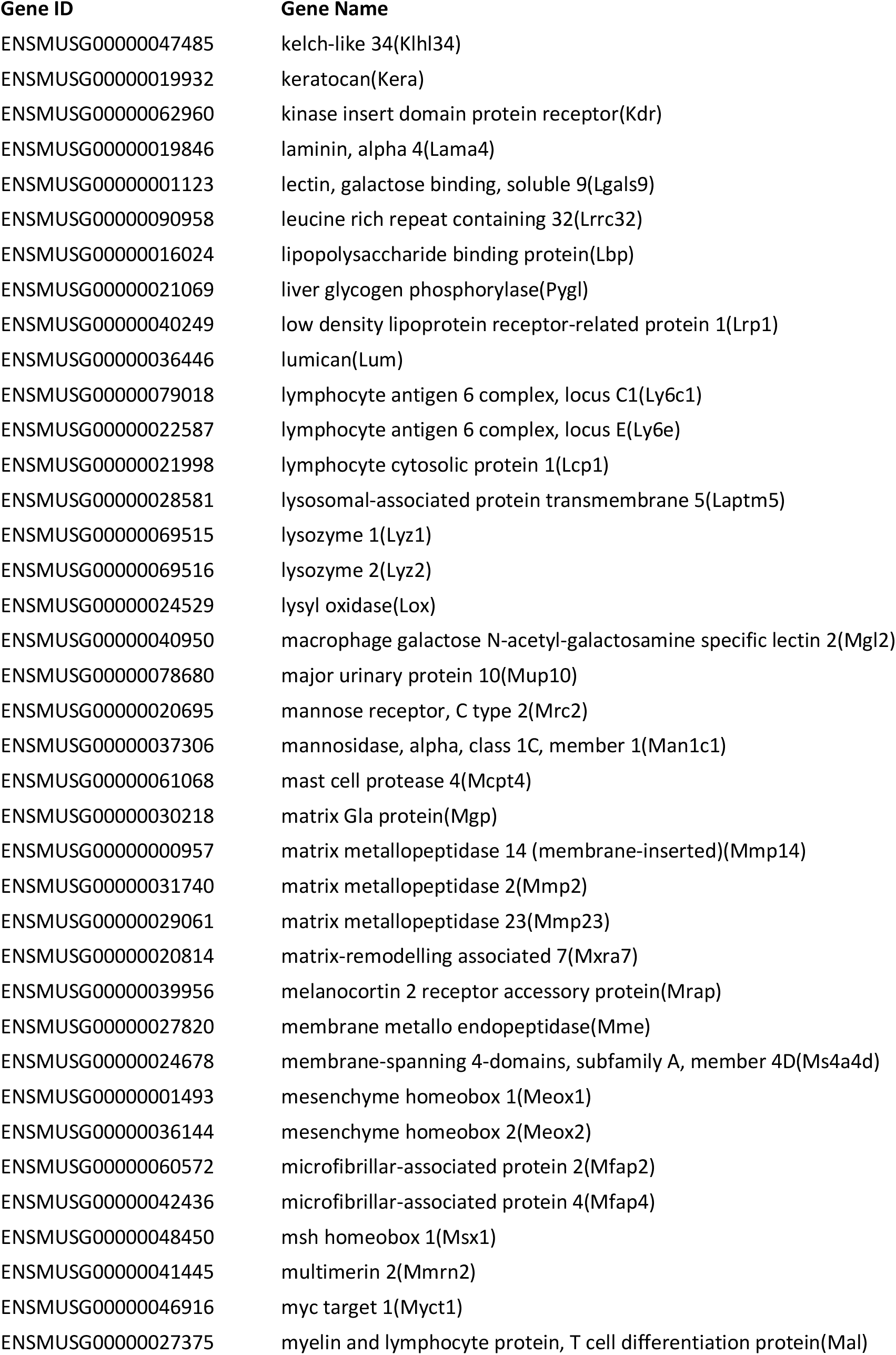
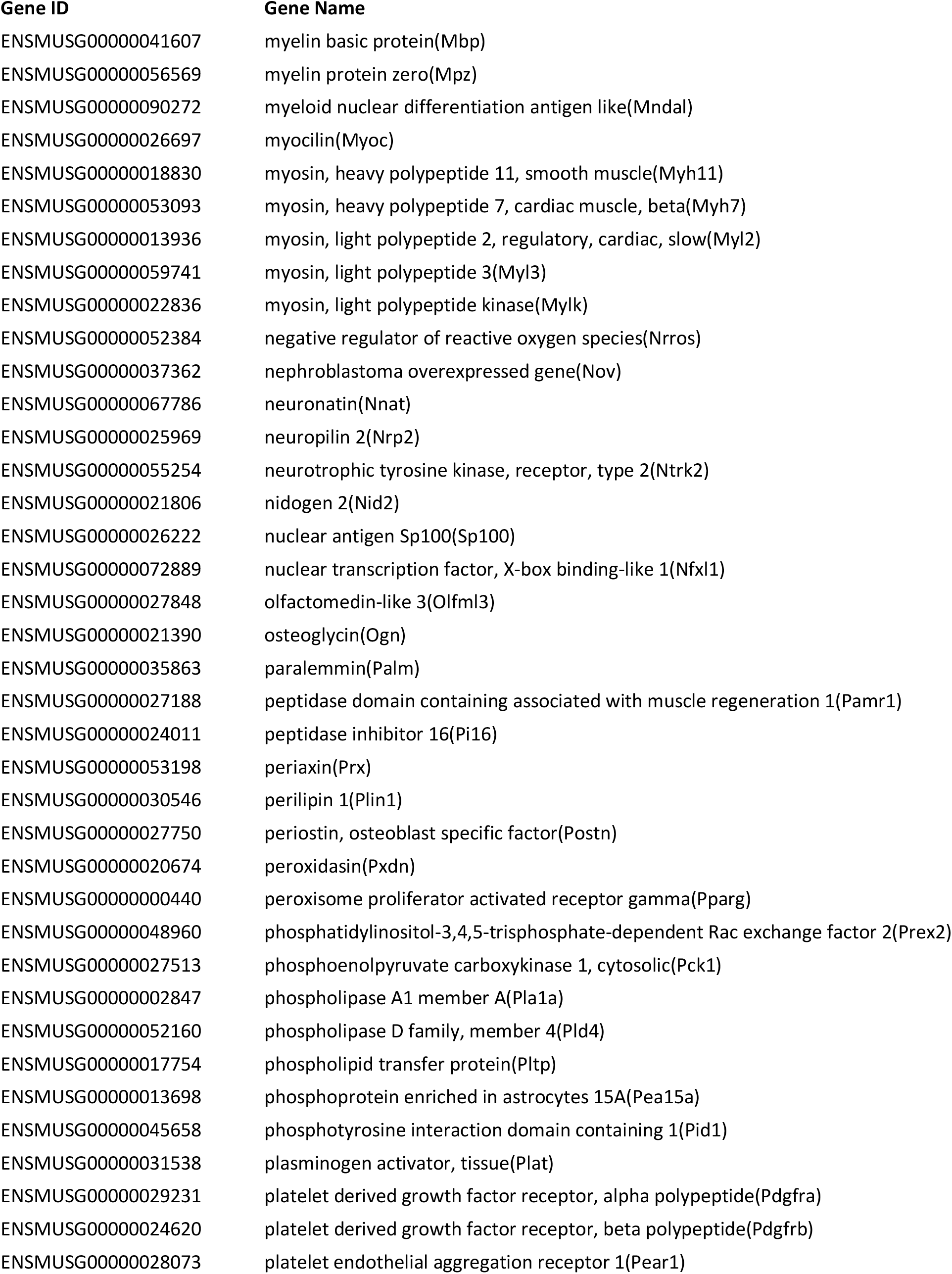
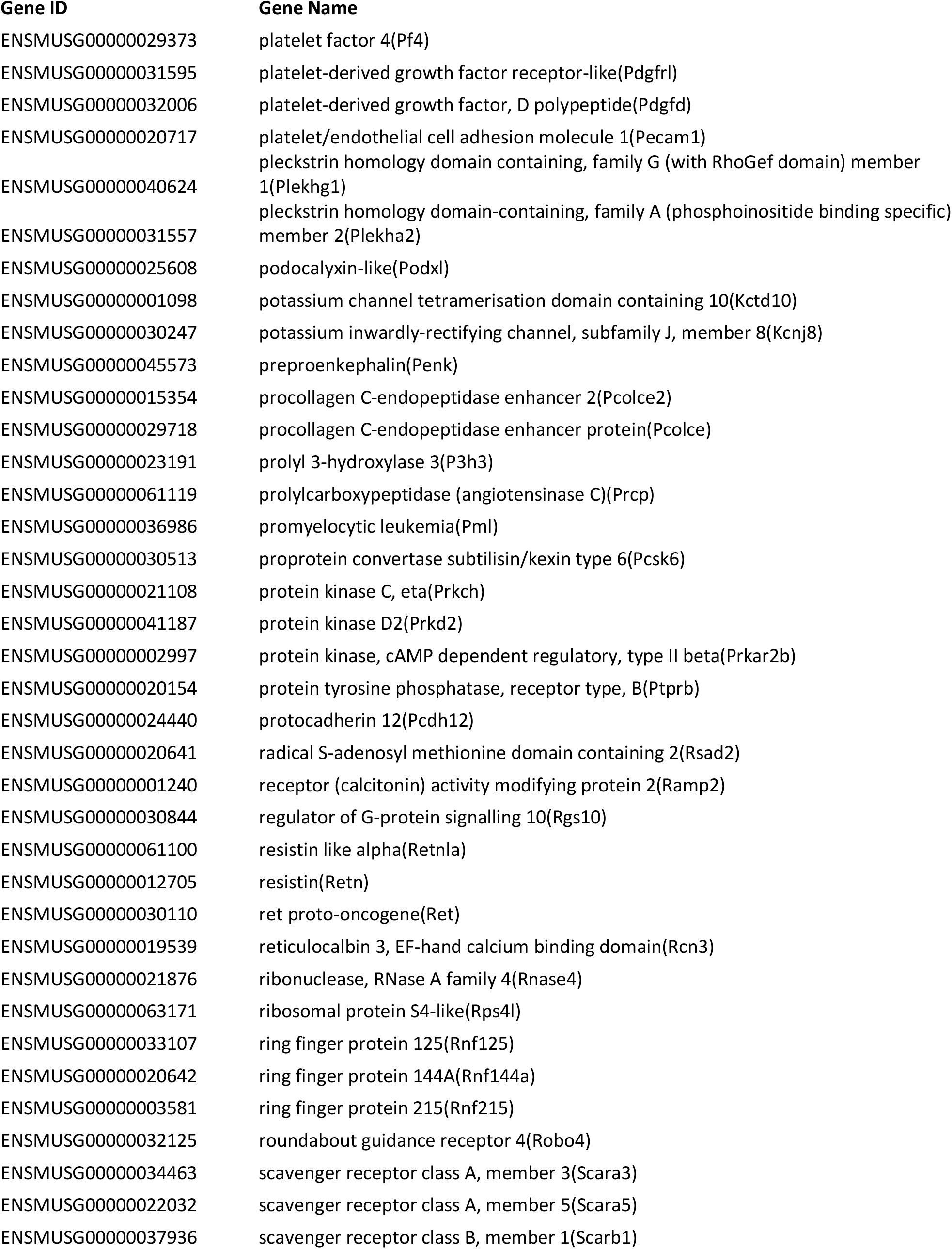
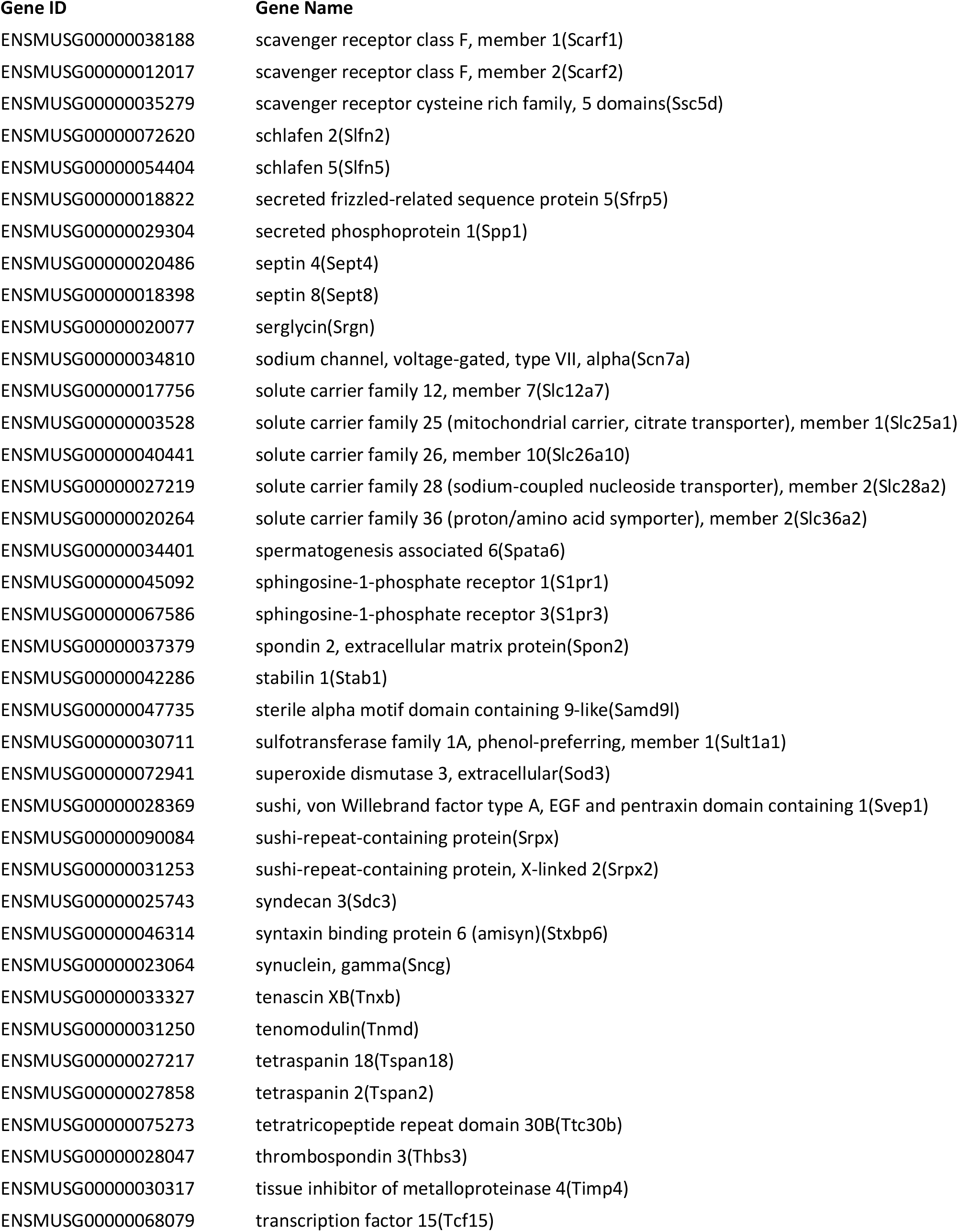
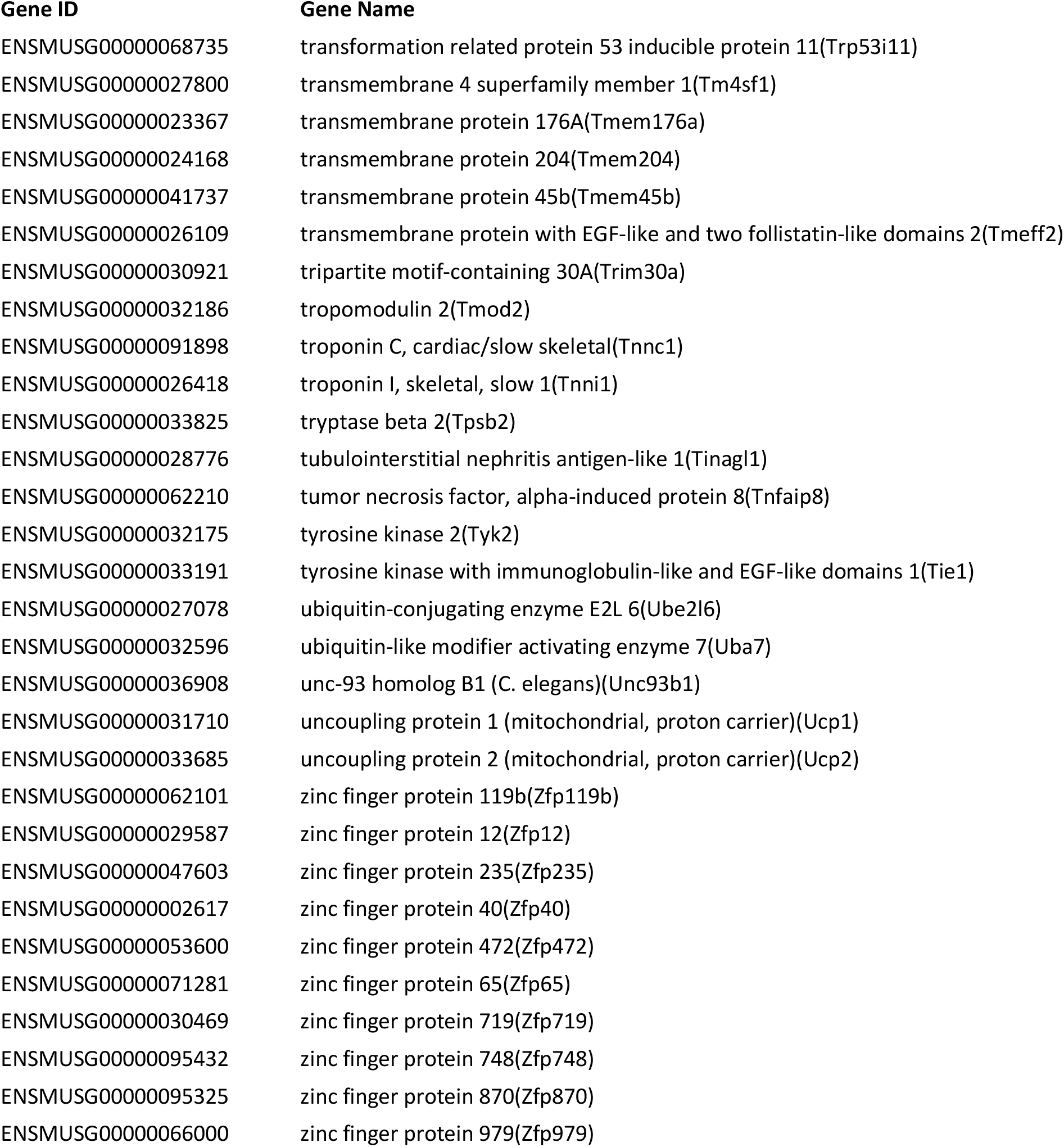
List of genes found in the whole muscle but not expressed in myofibers, defined as having an RPM value of at least 10 in the whole muscle and 0 in the single fiber.

**Table S1B:**
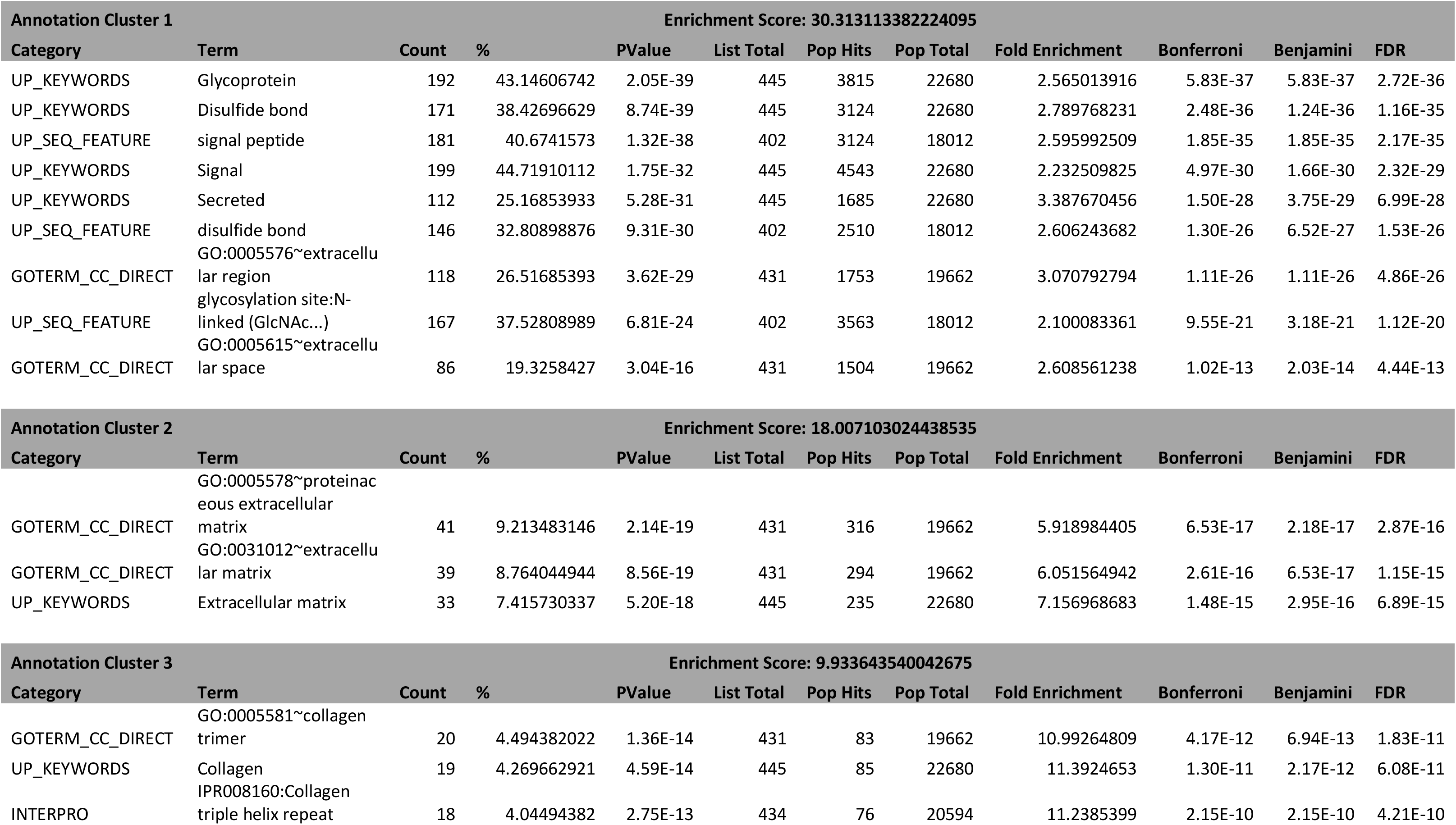
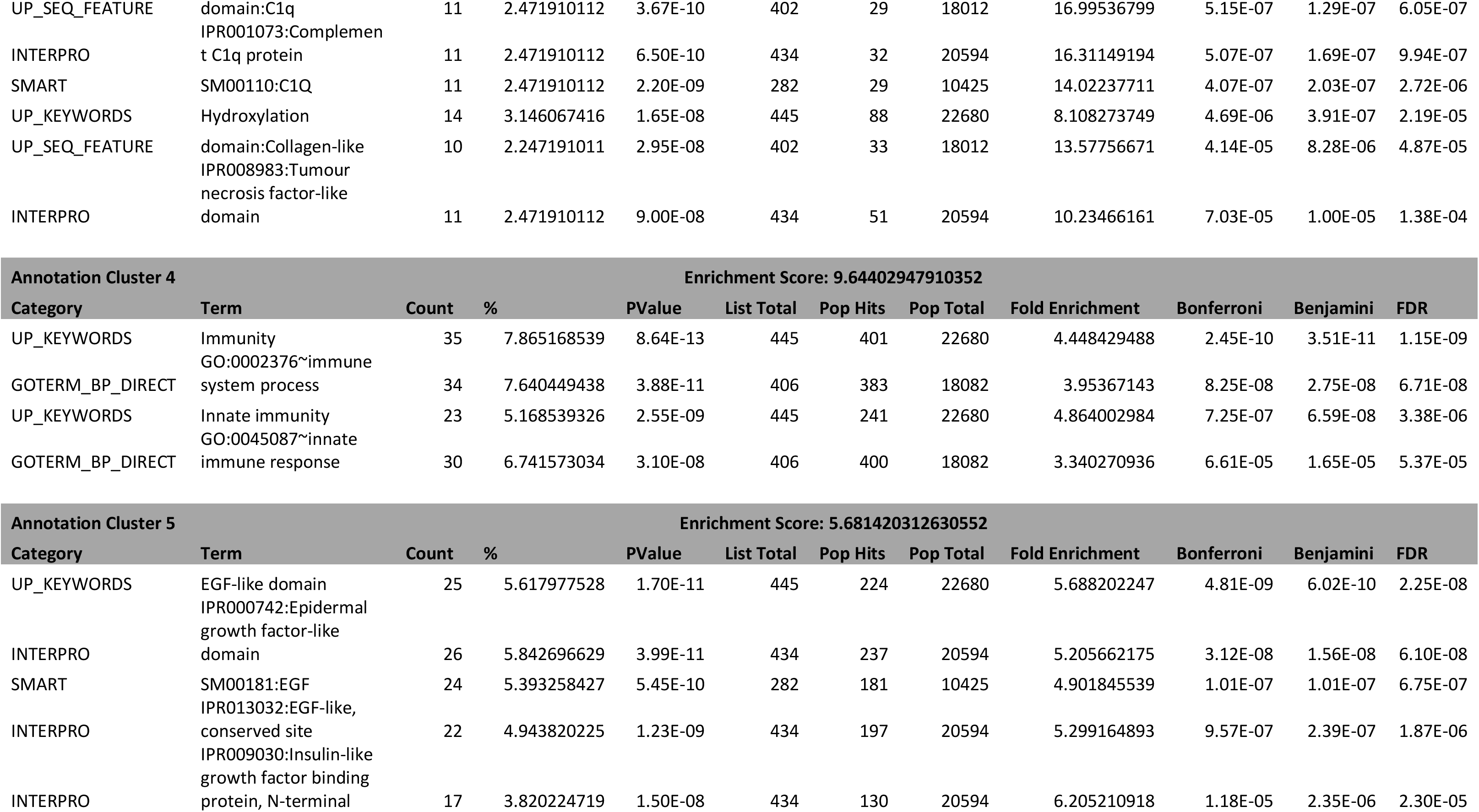
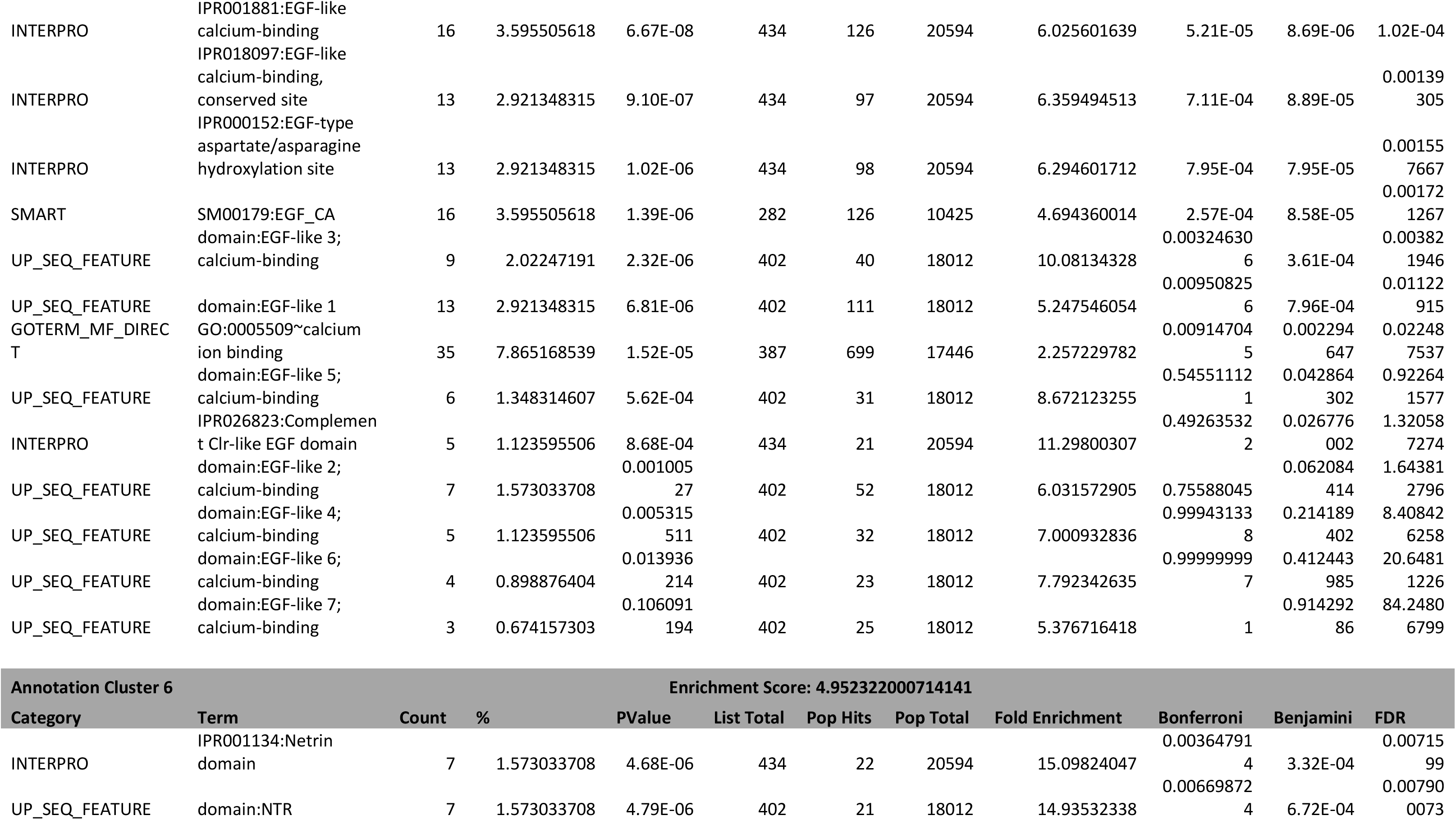
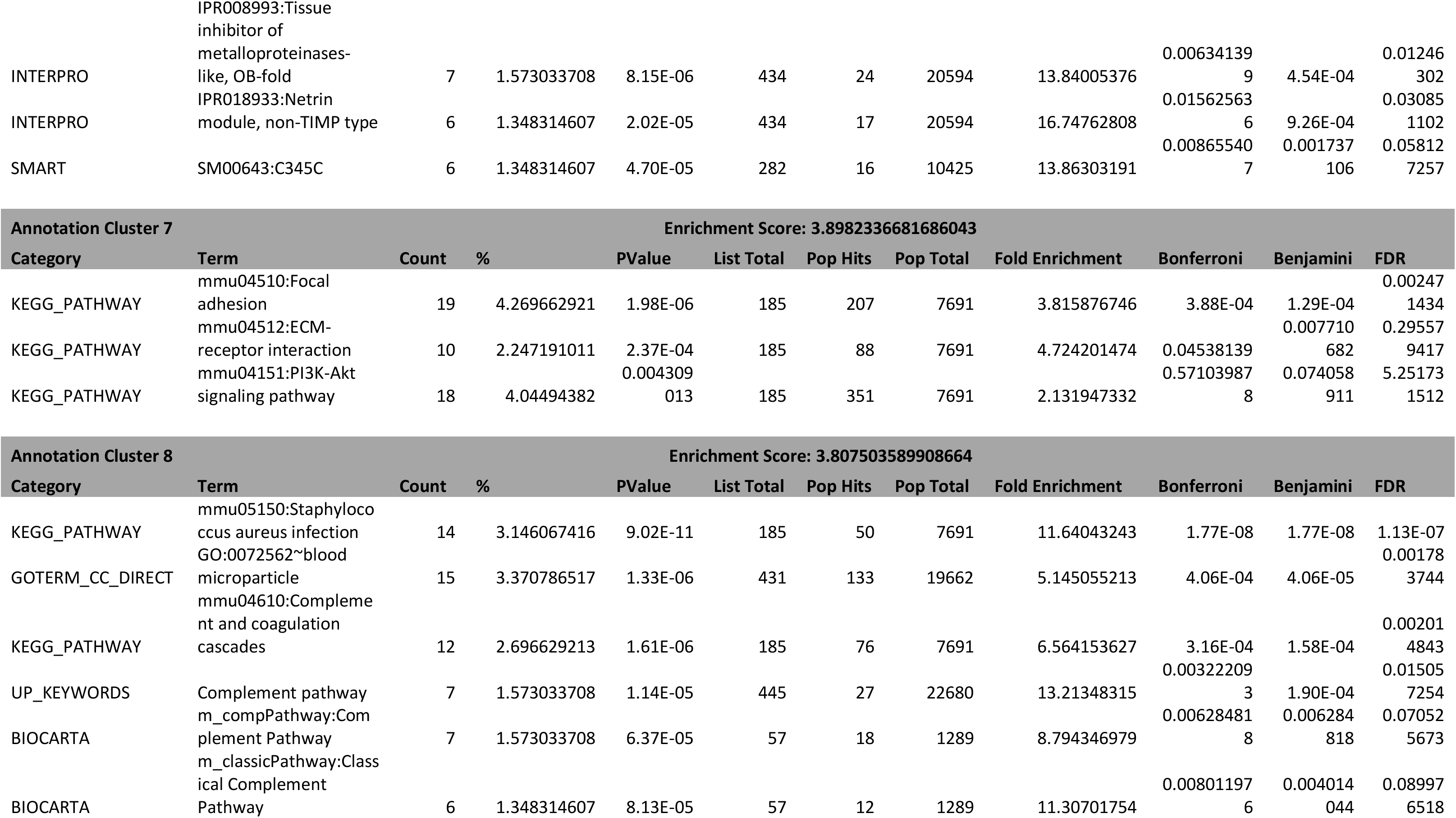
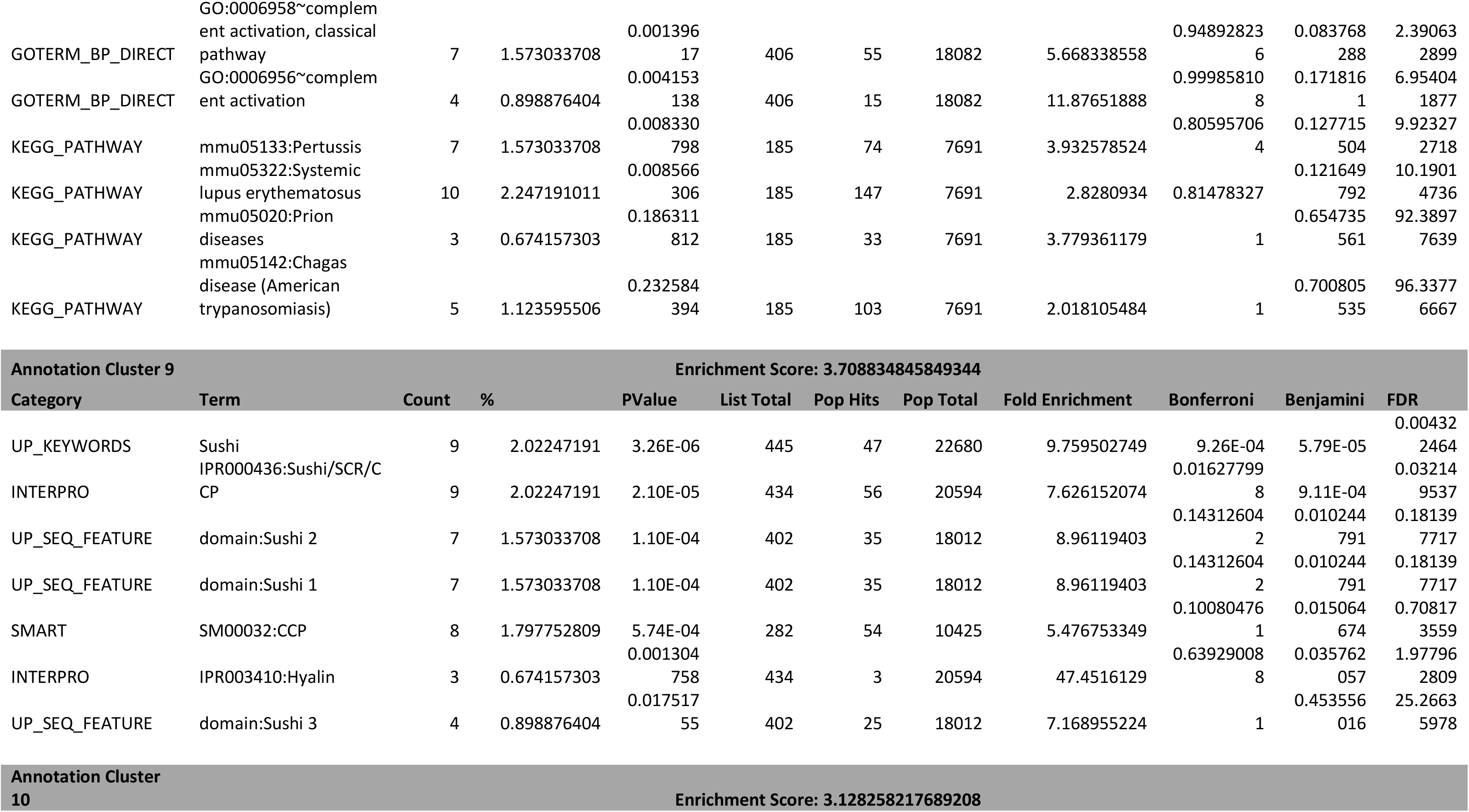
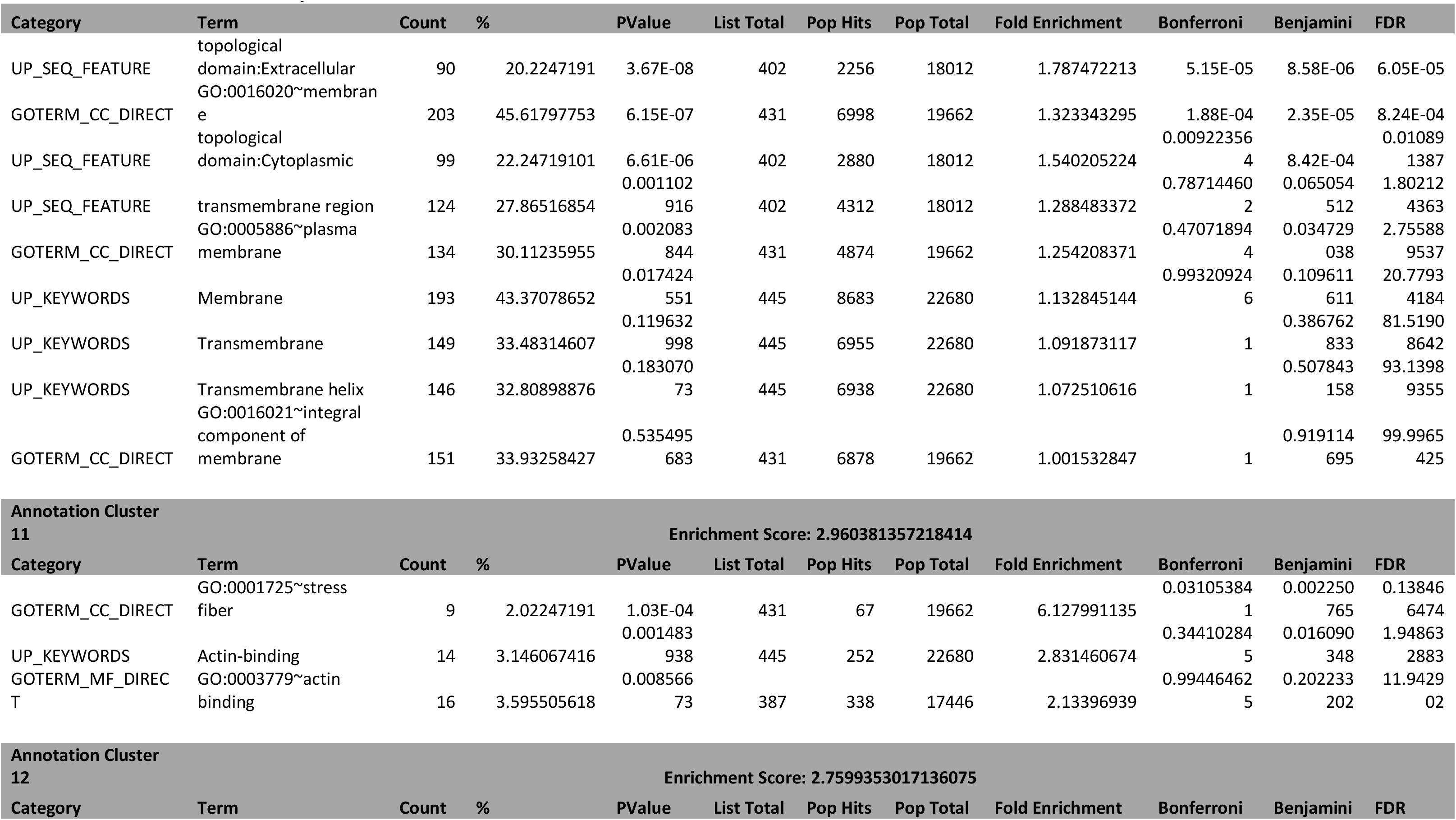
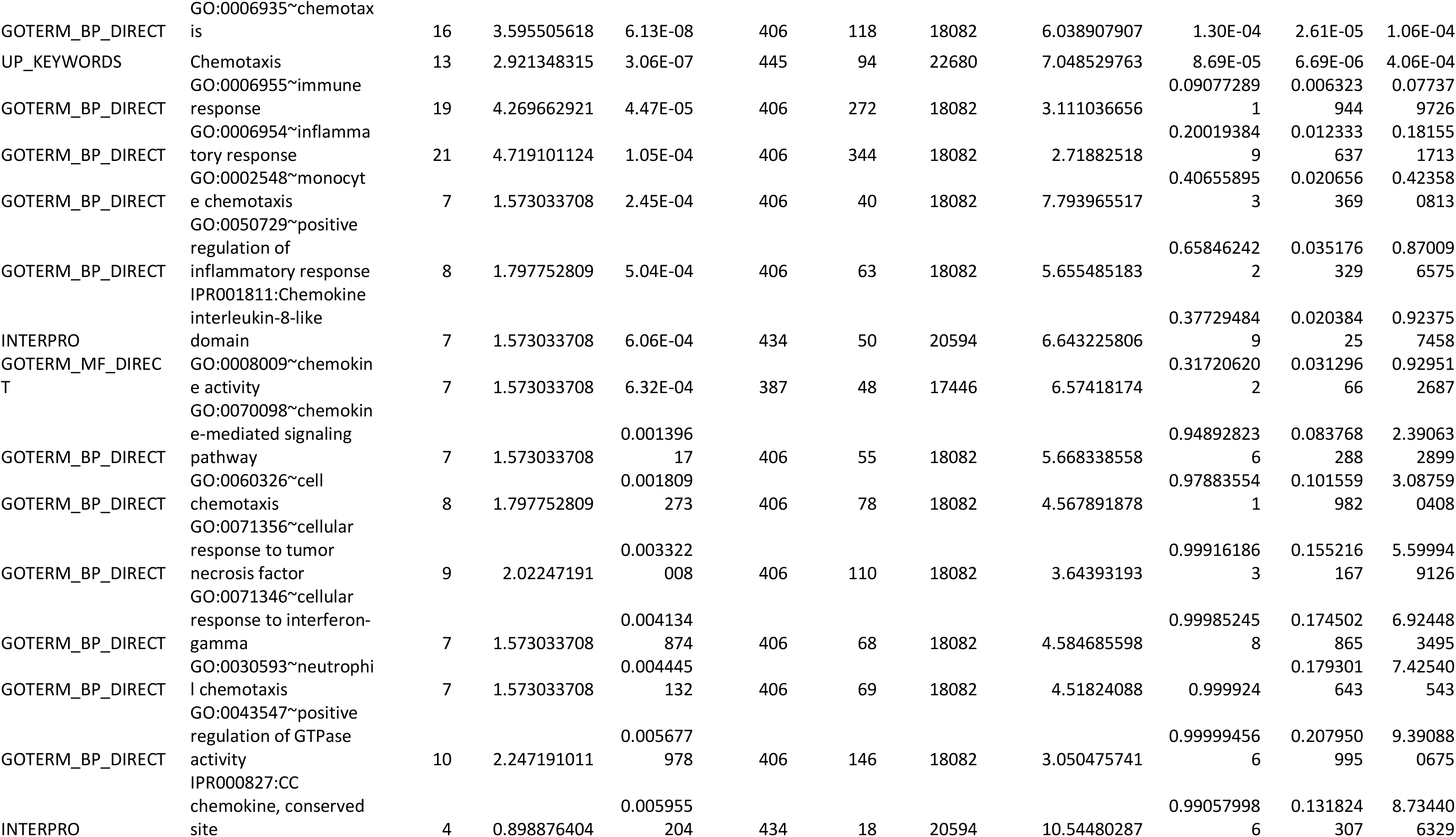
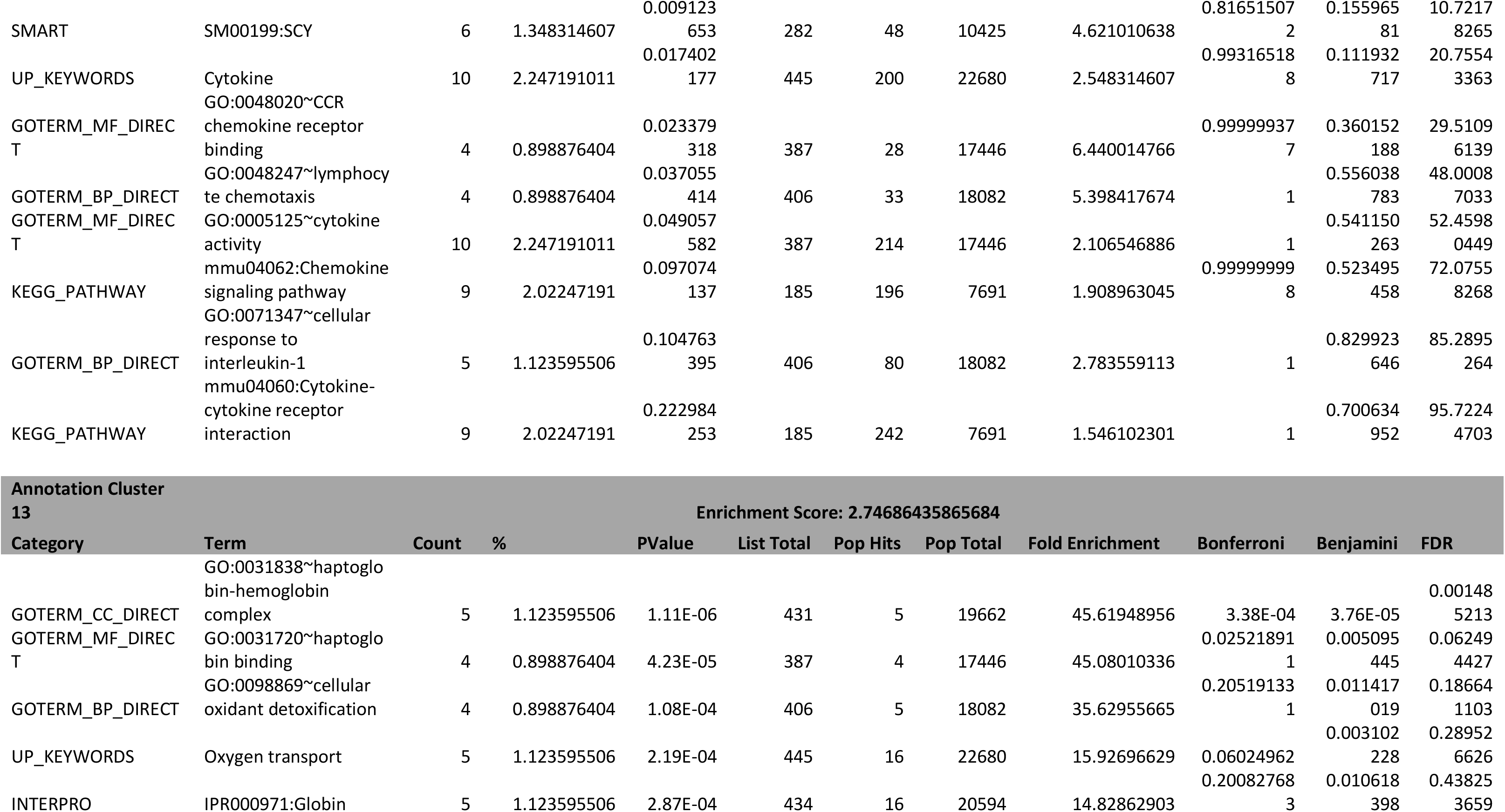
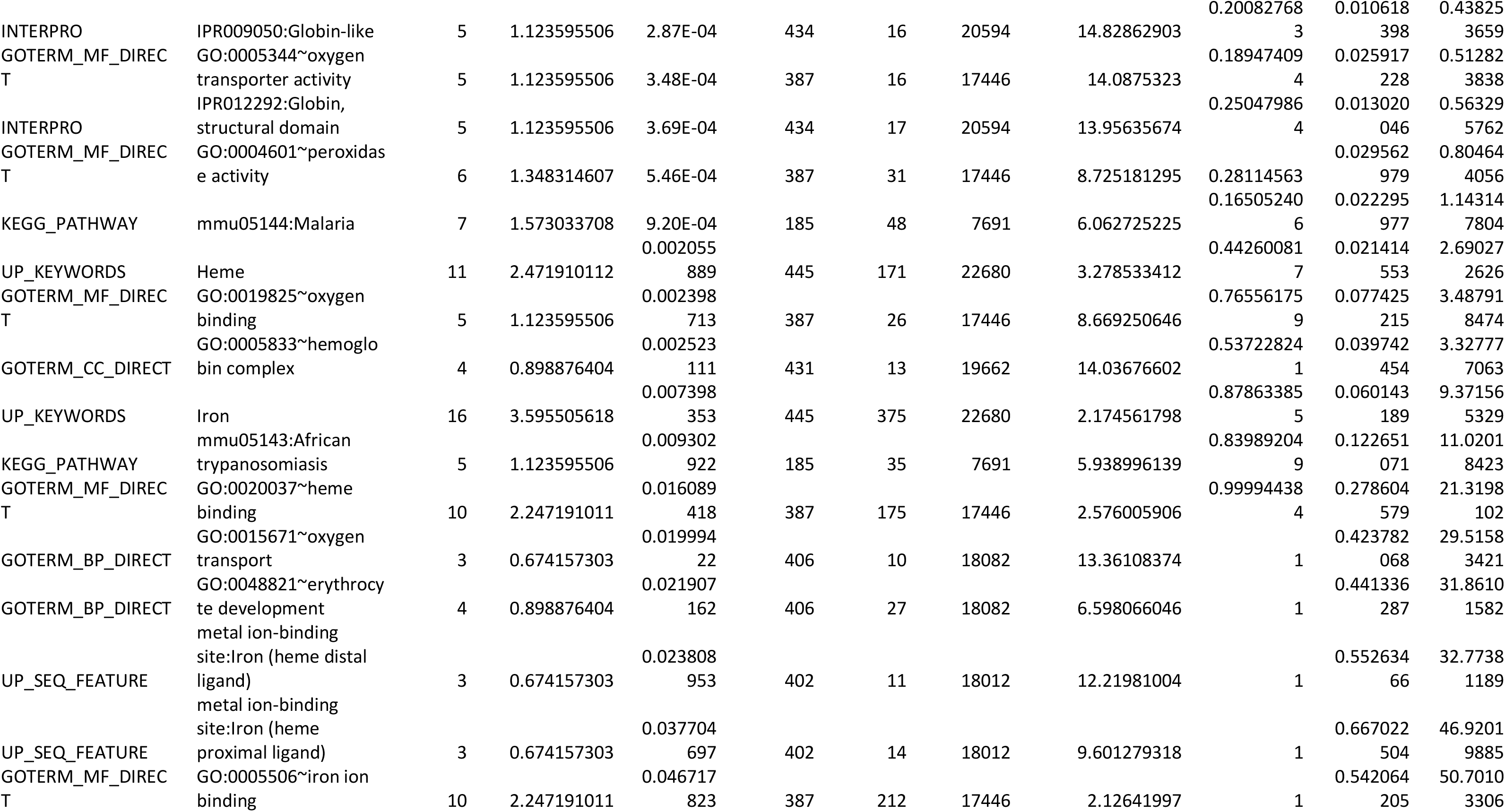
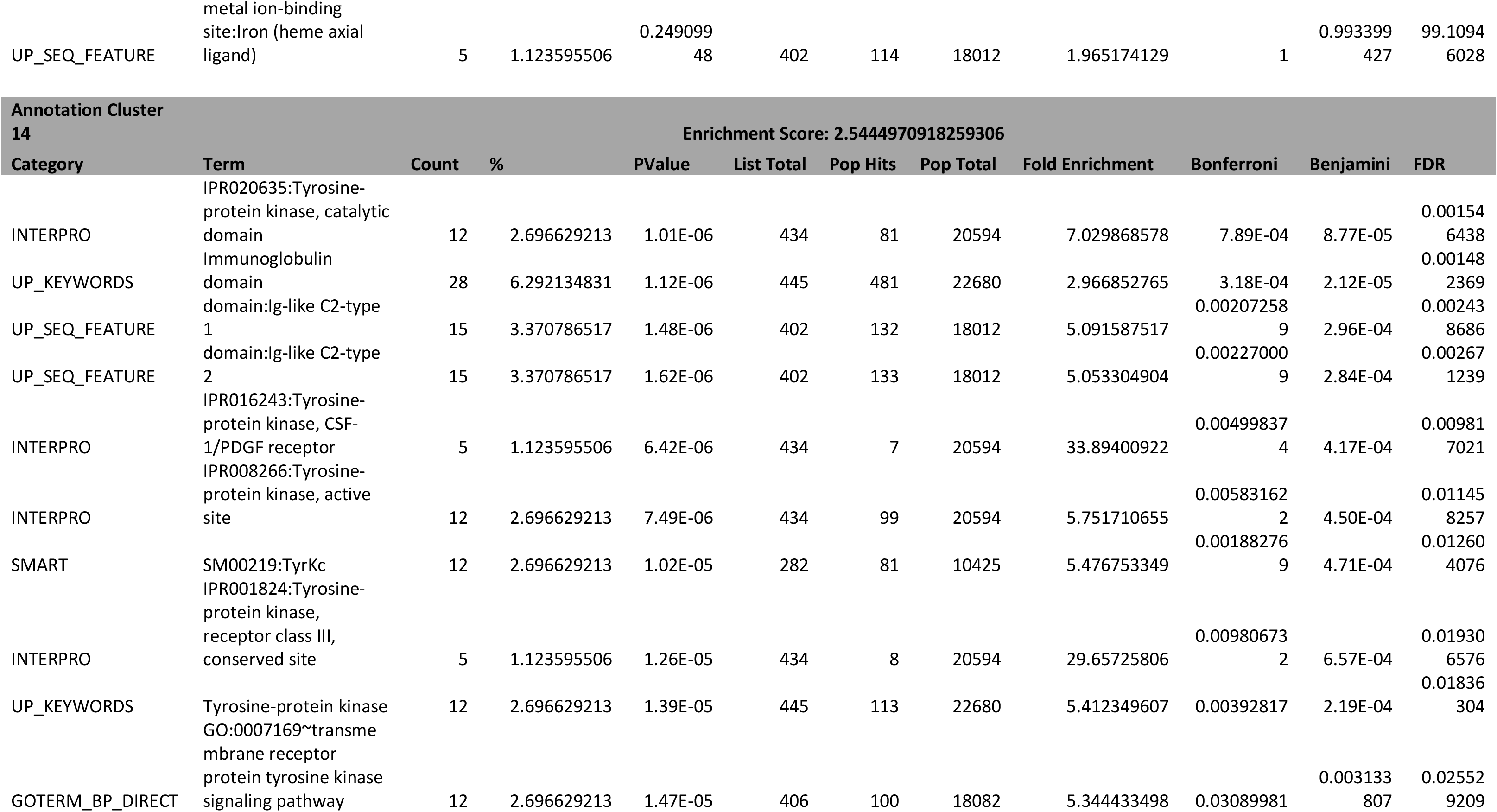
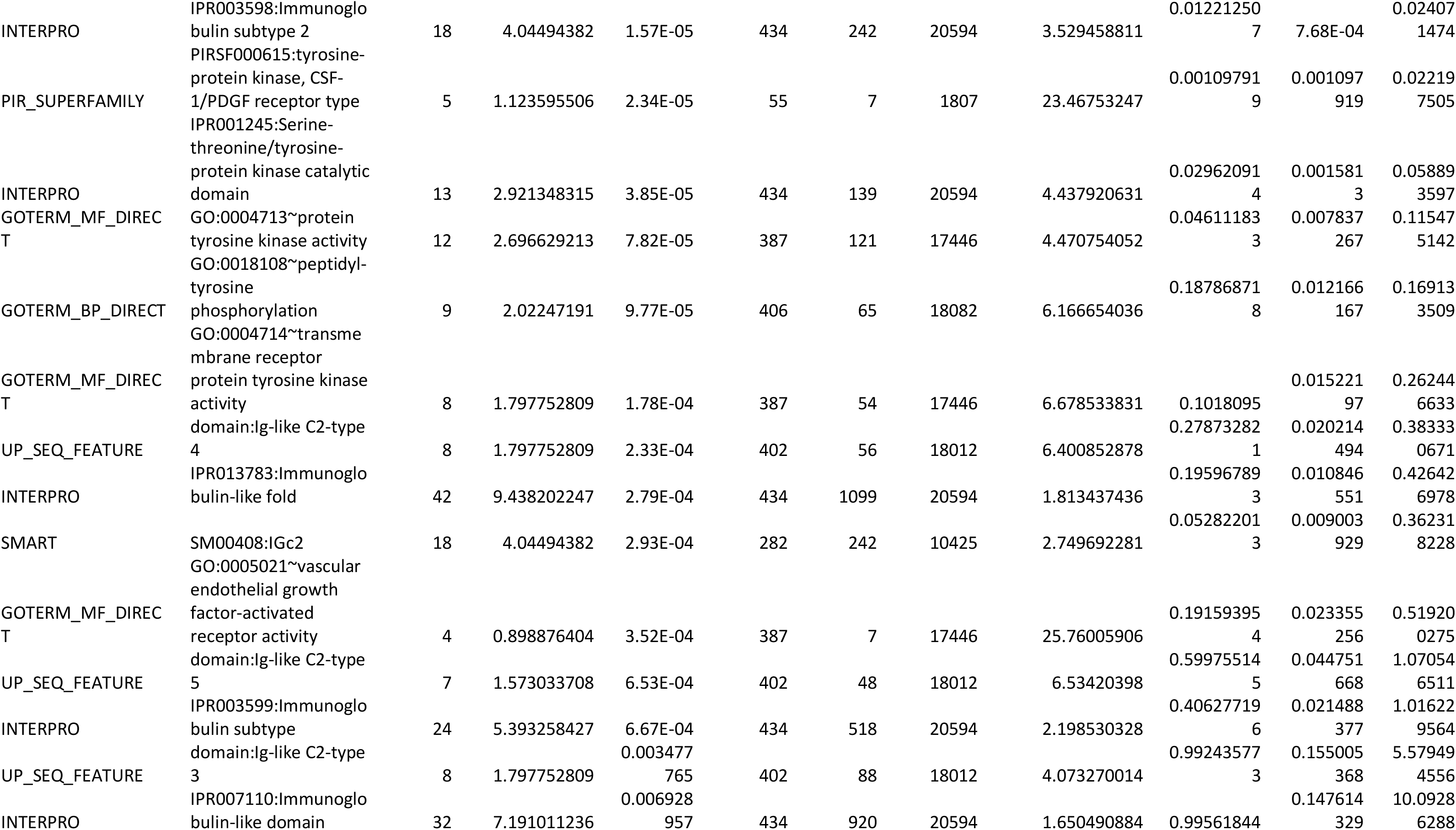
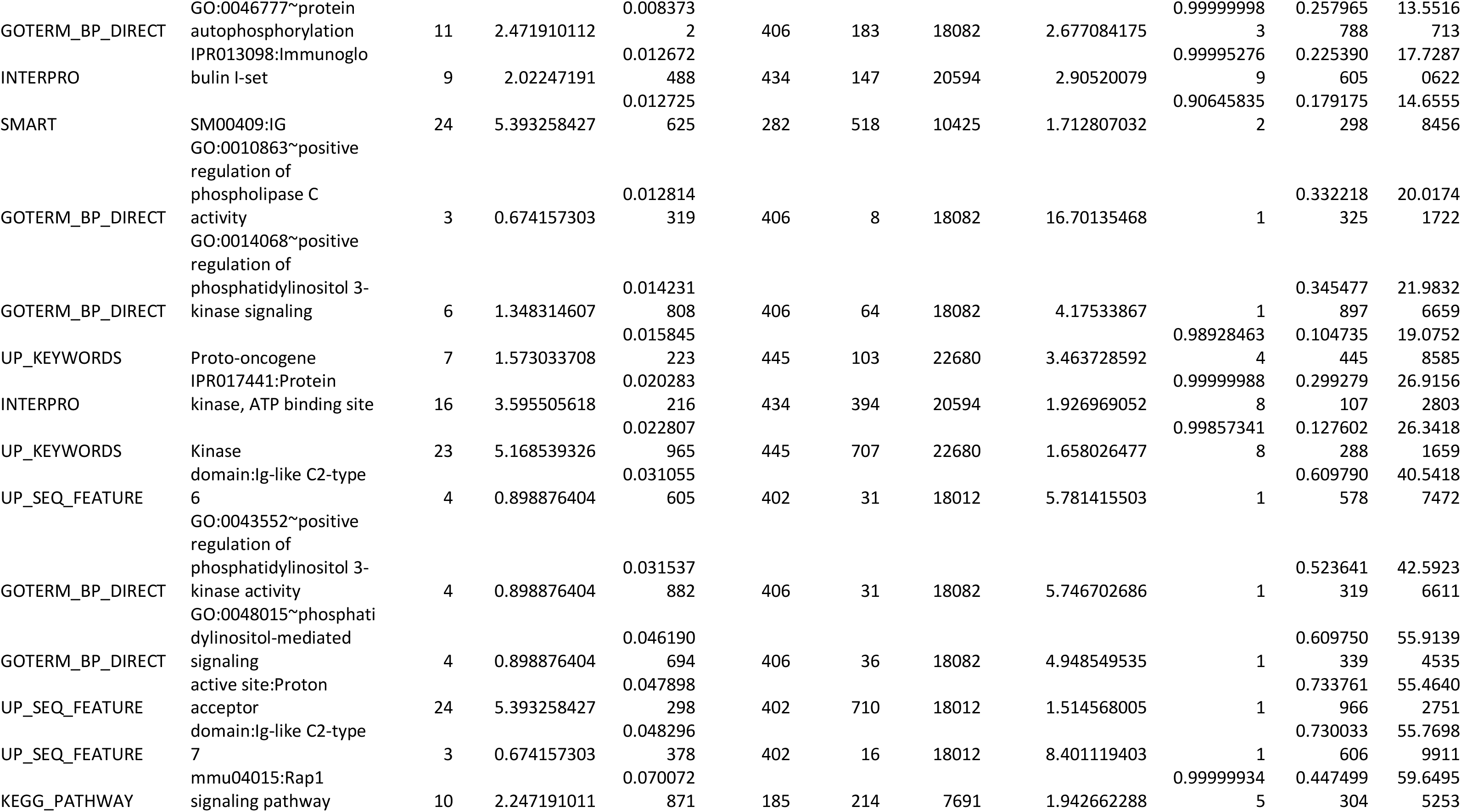
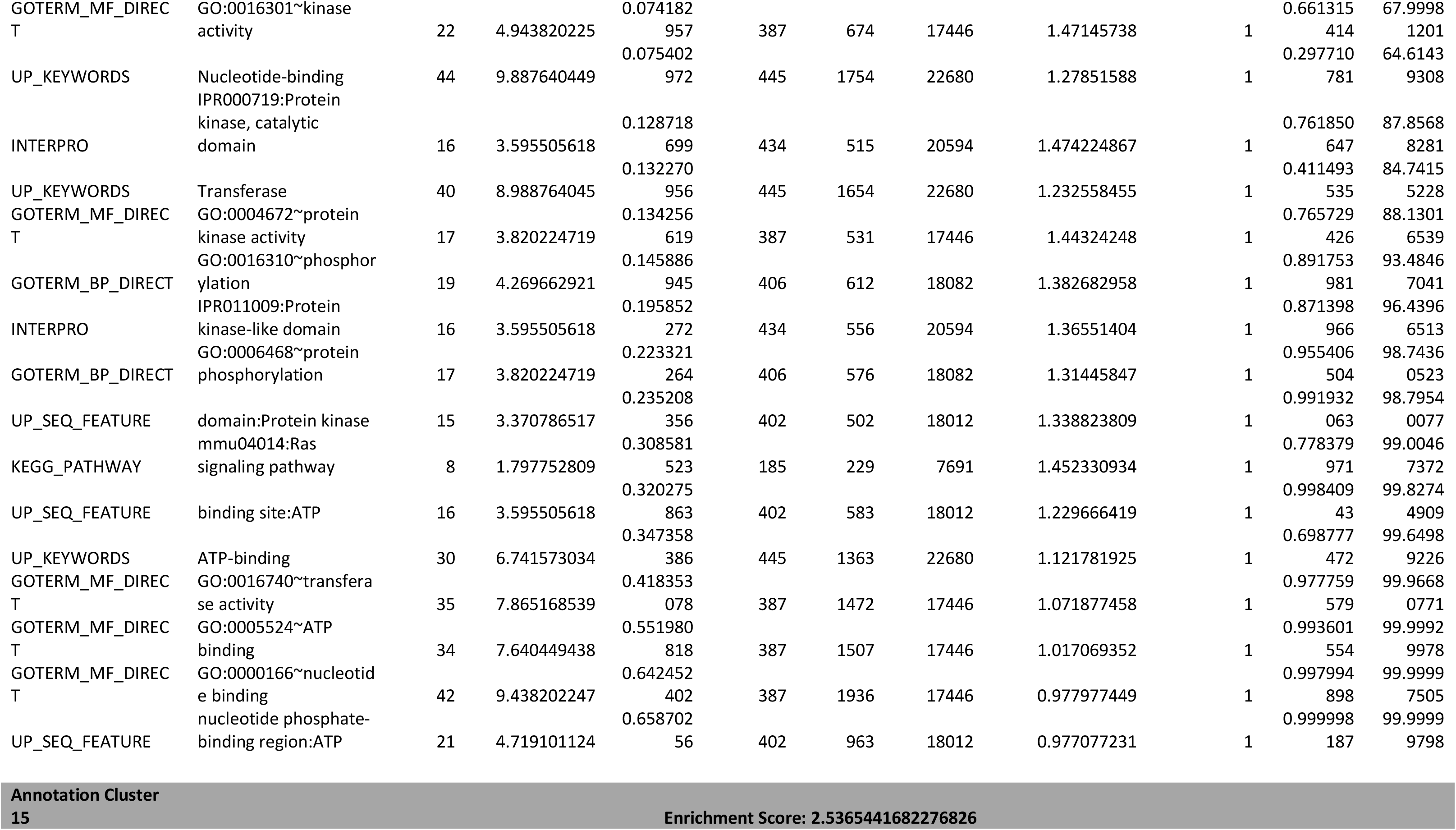
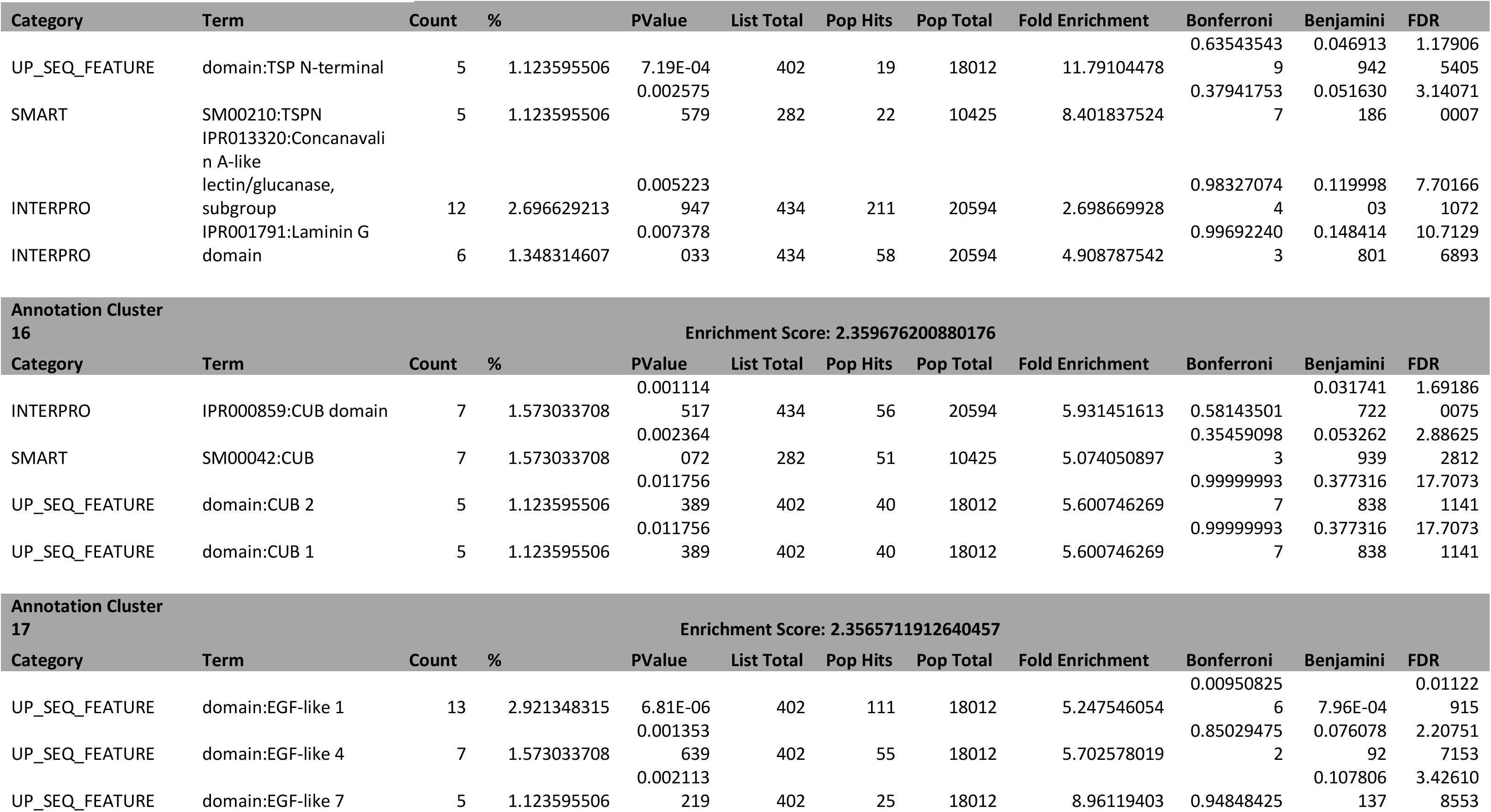
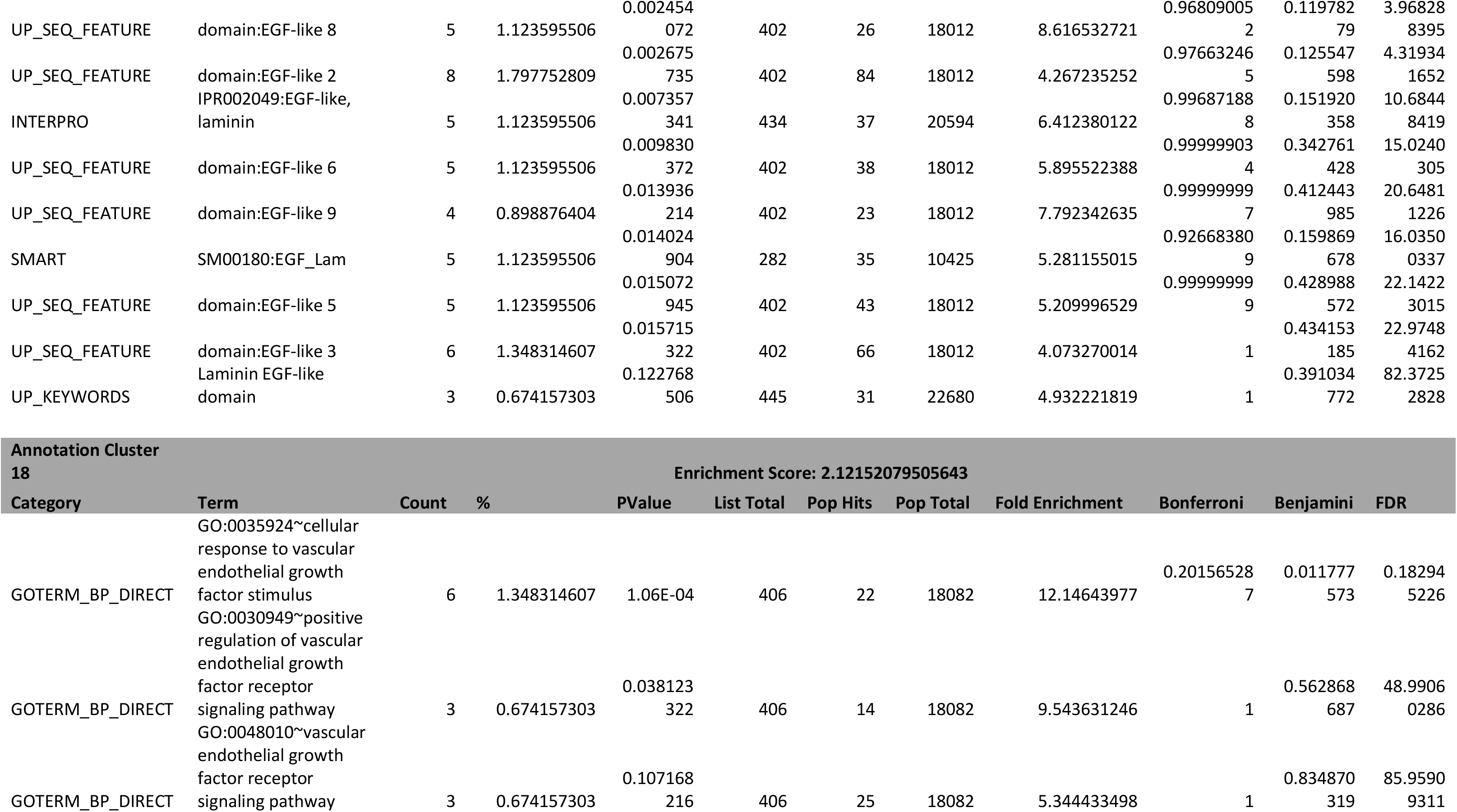
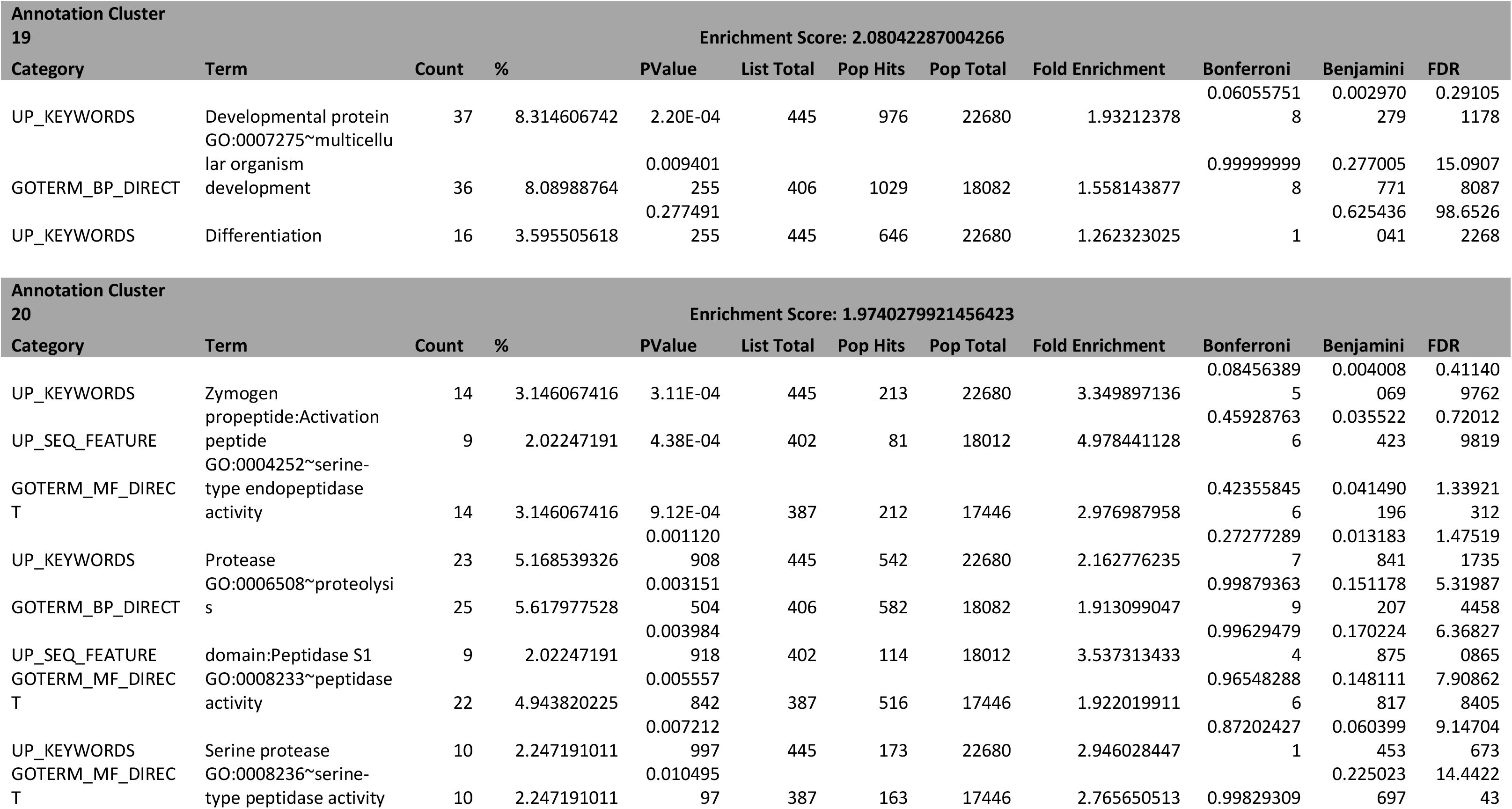
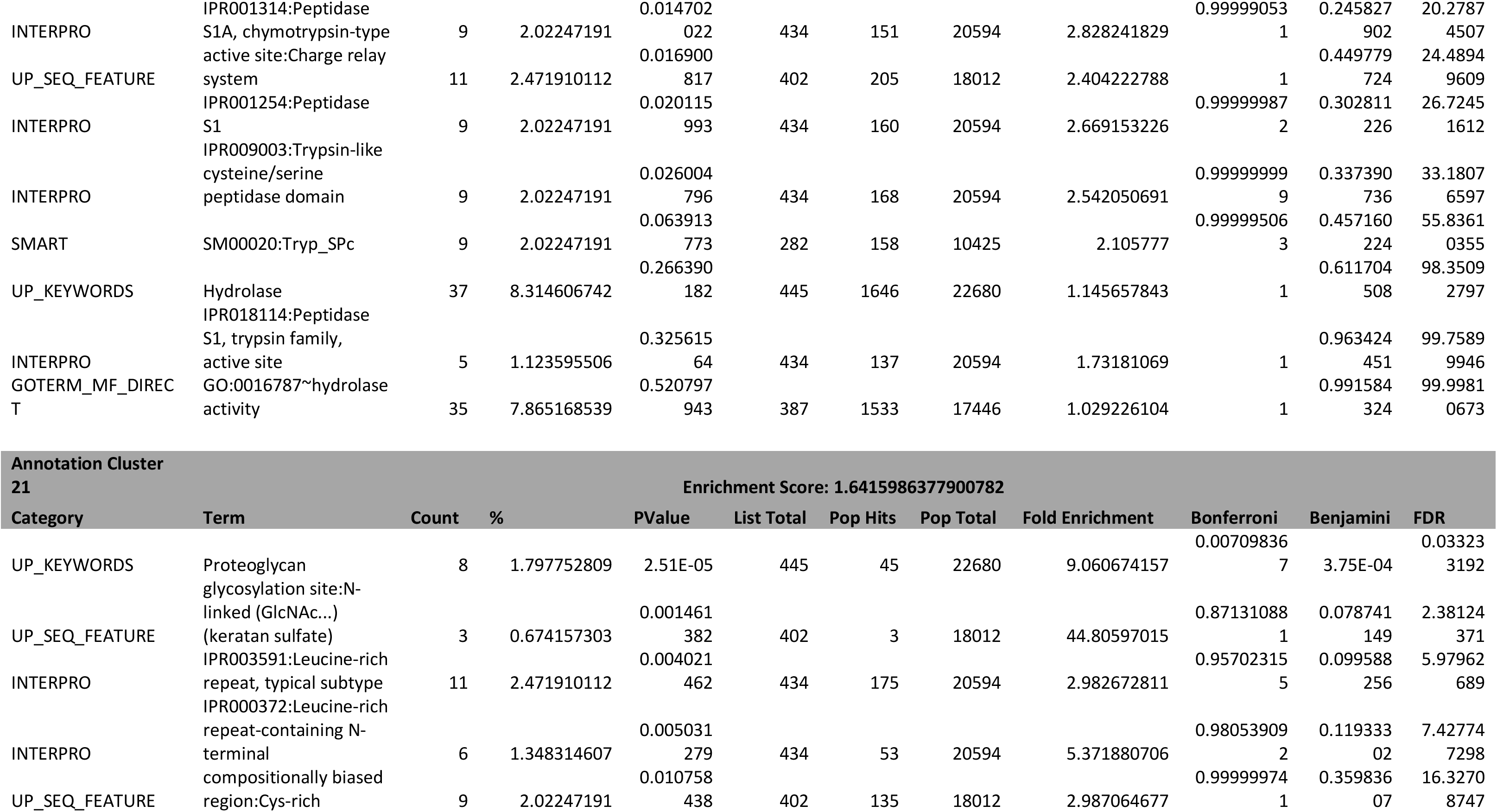
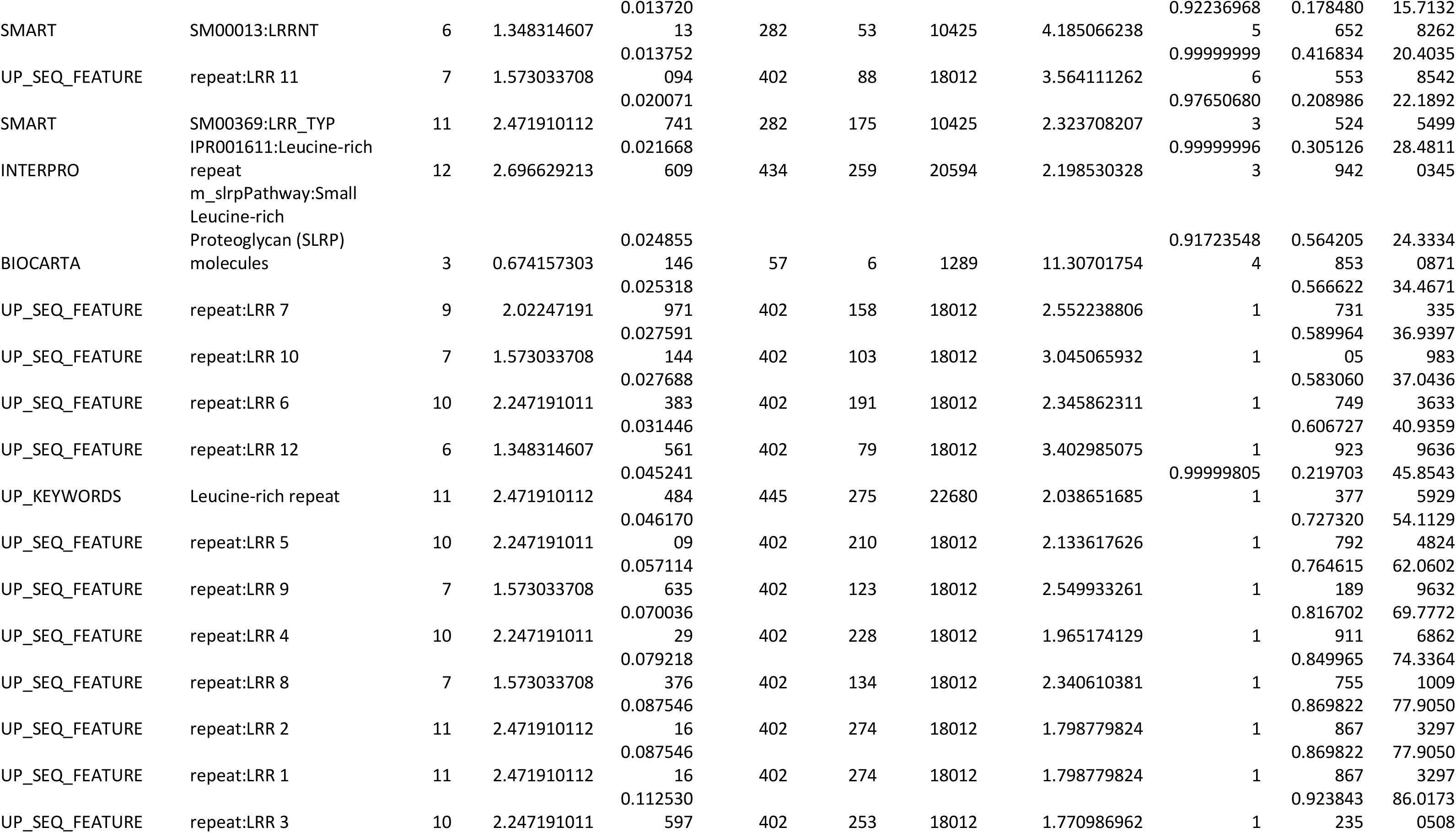
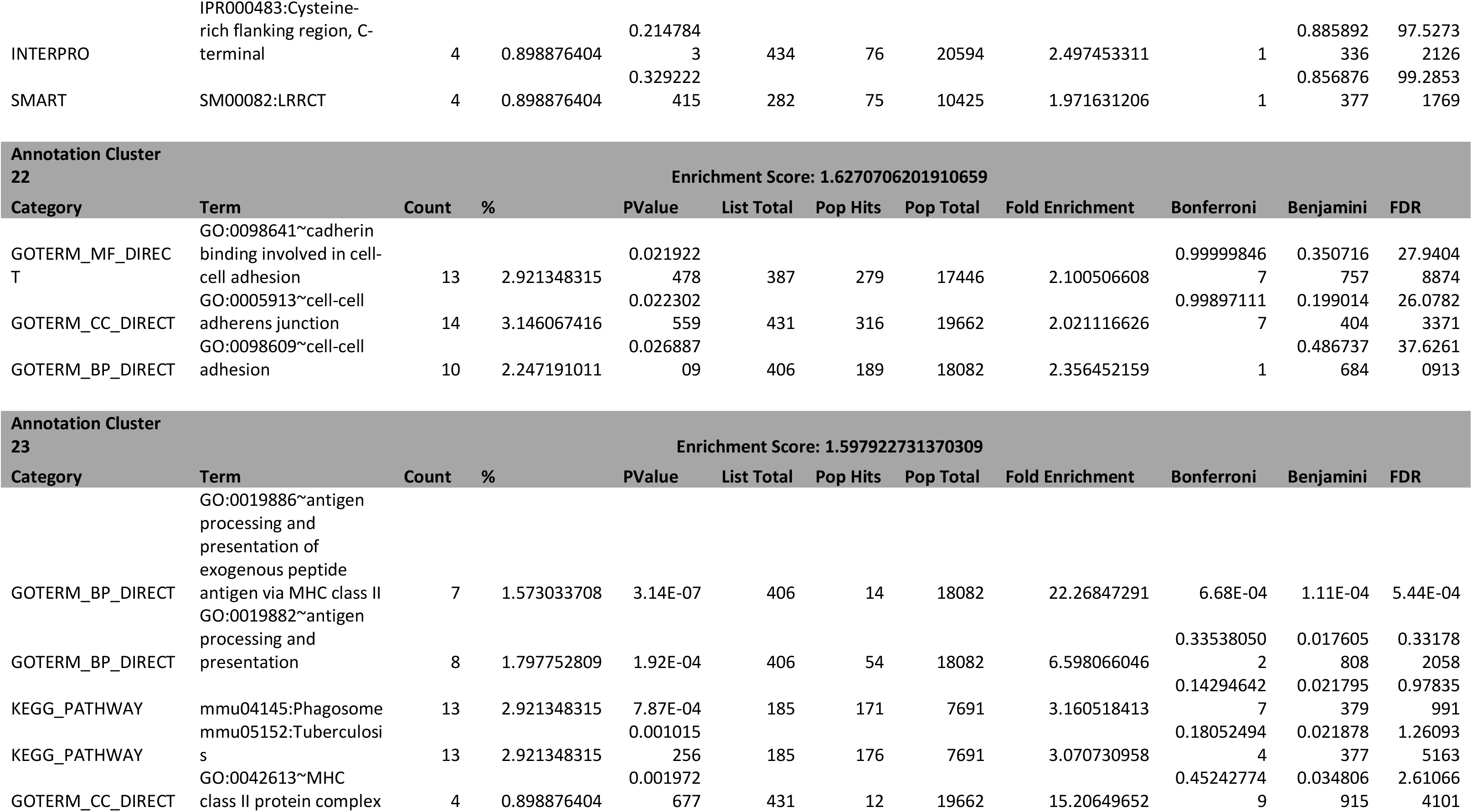
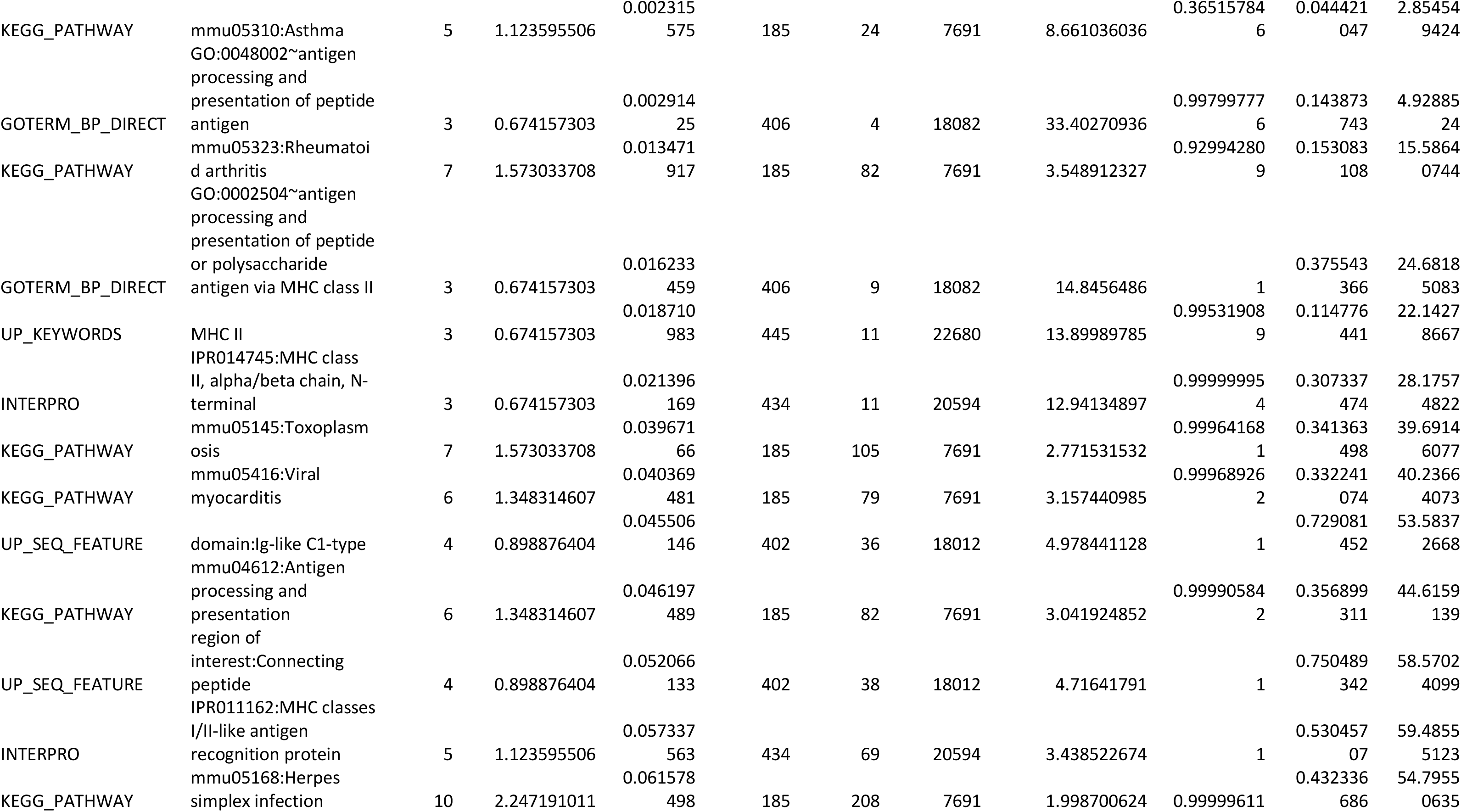
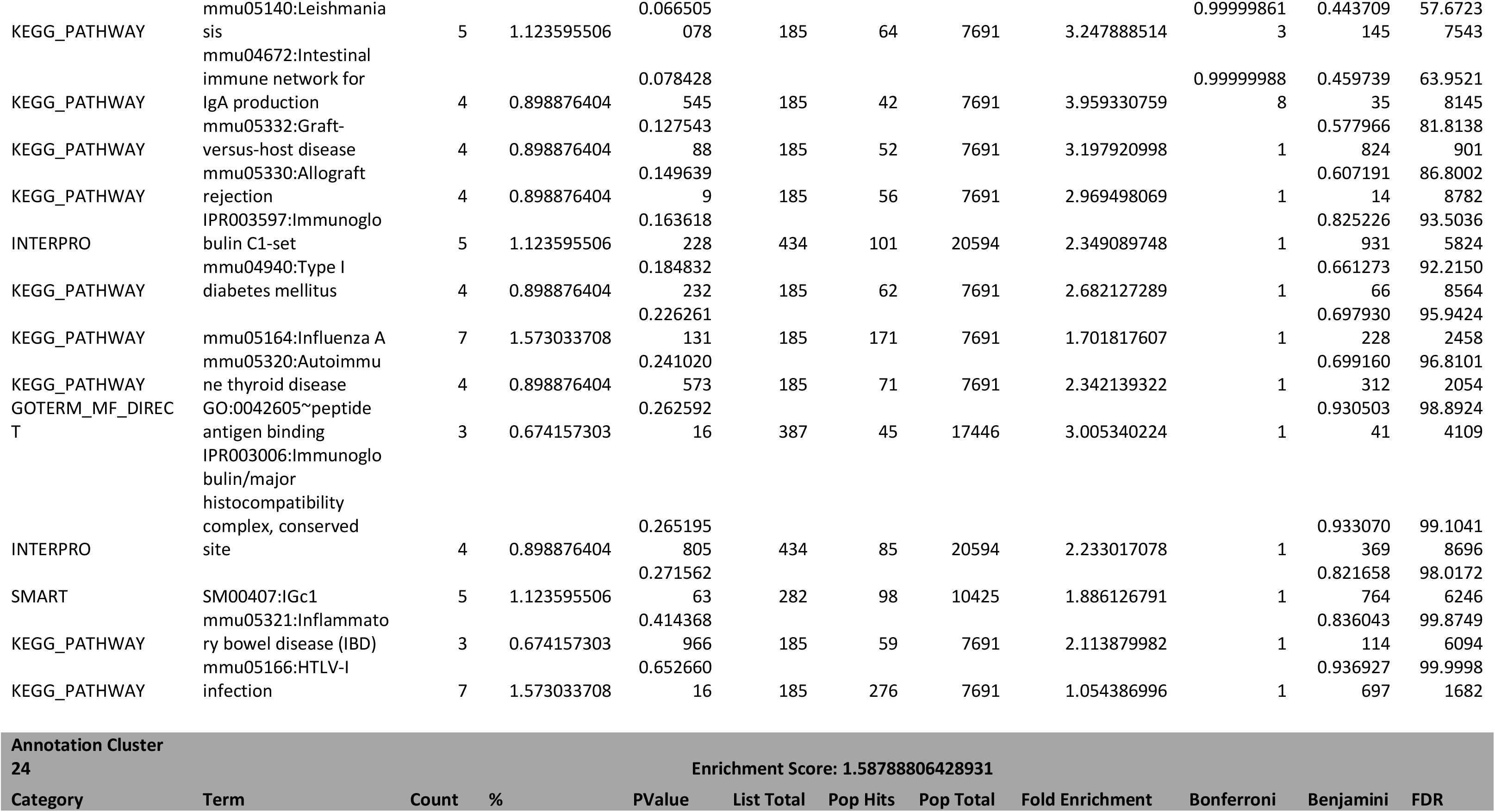
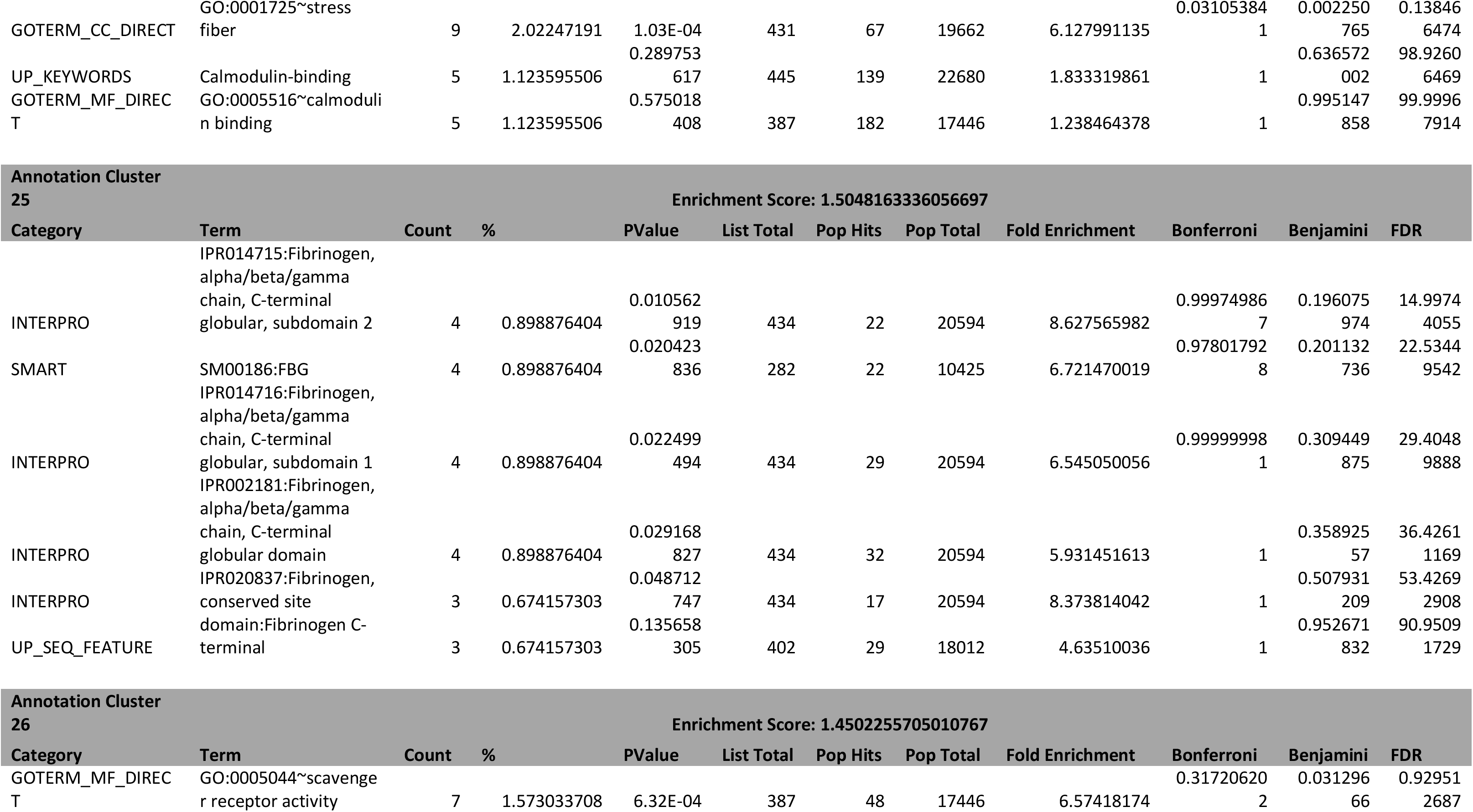
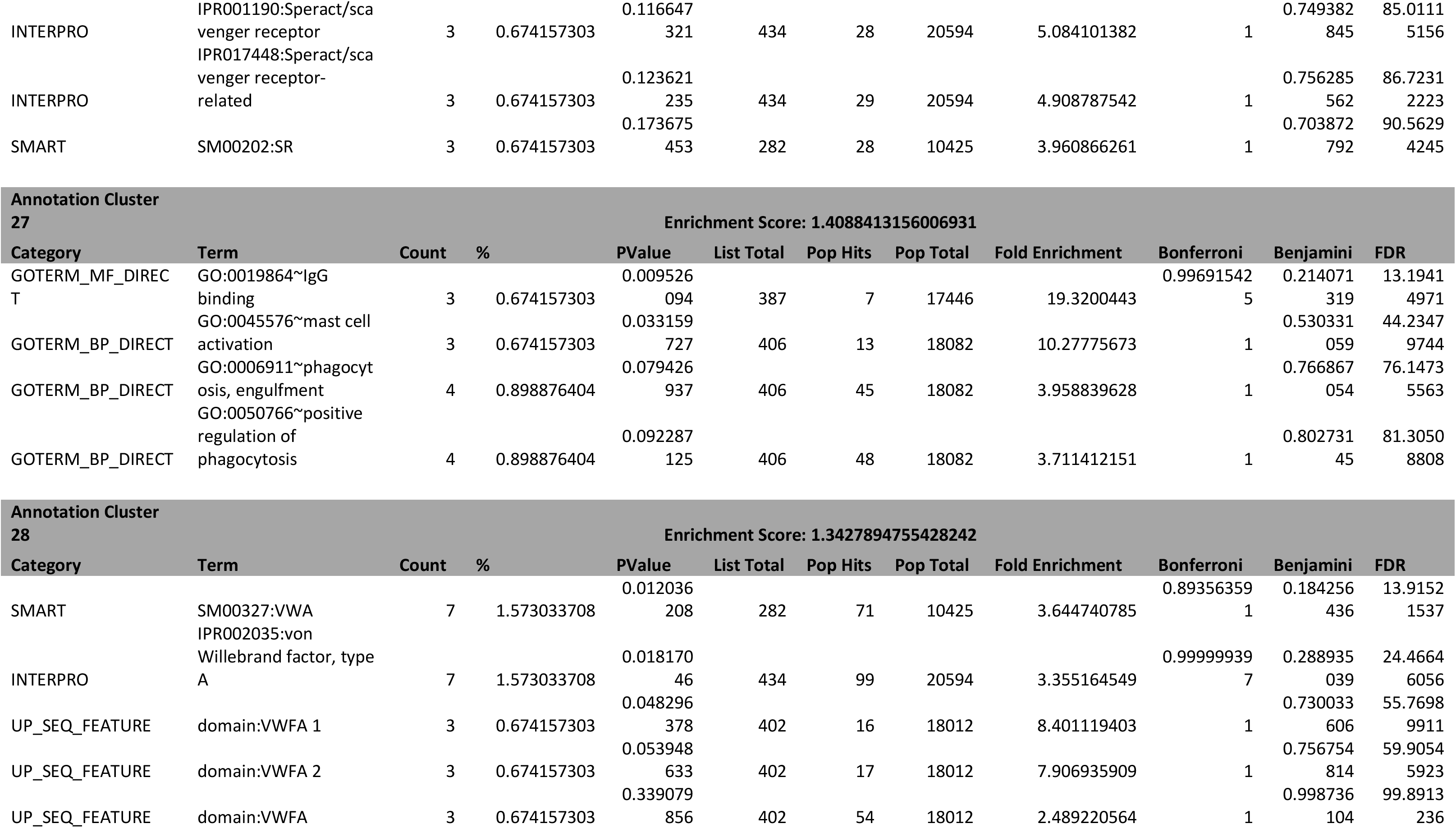
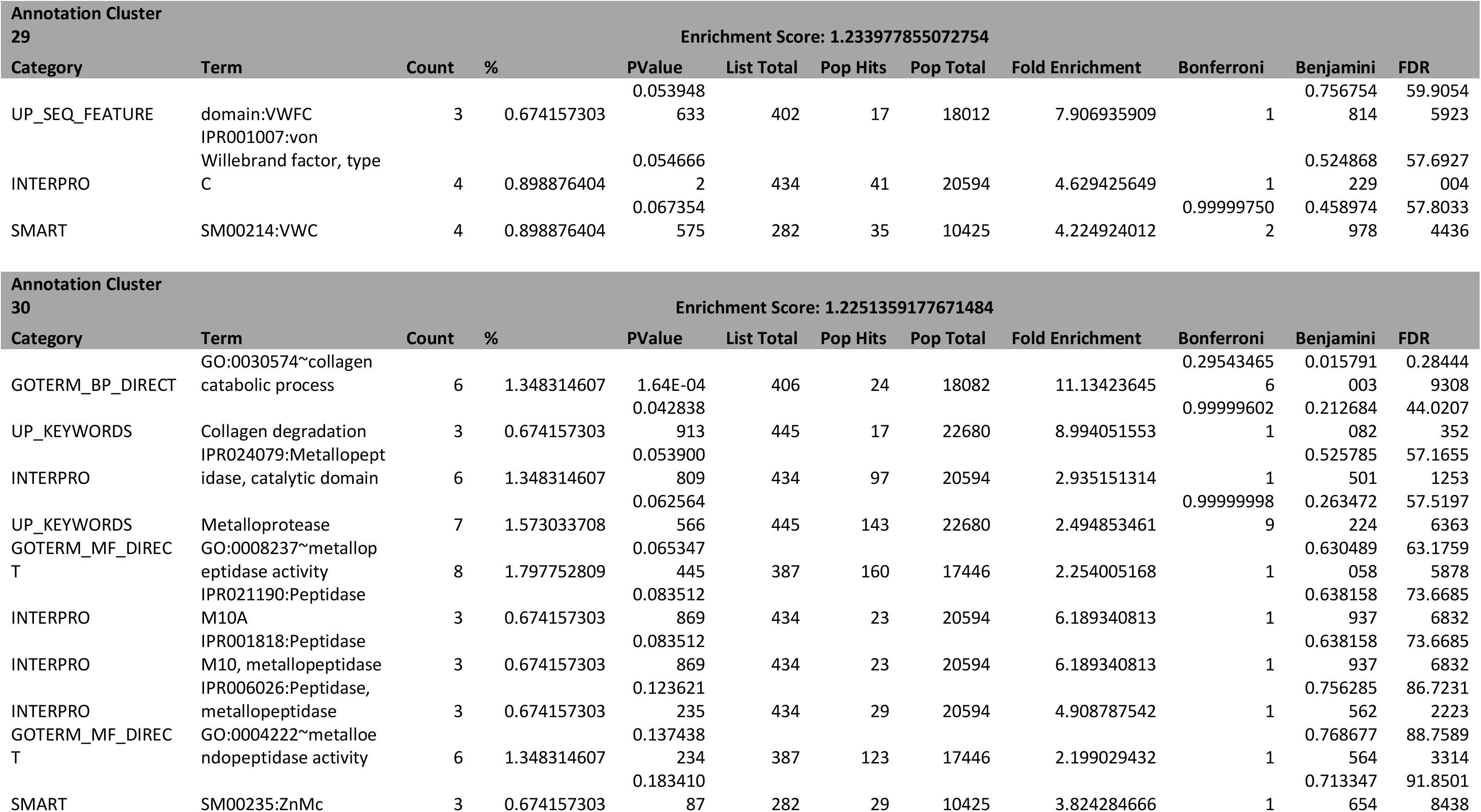
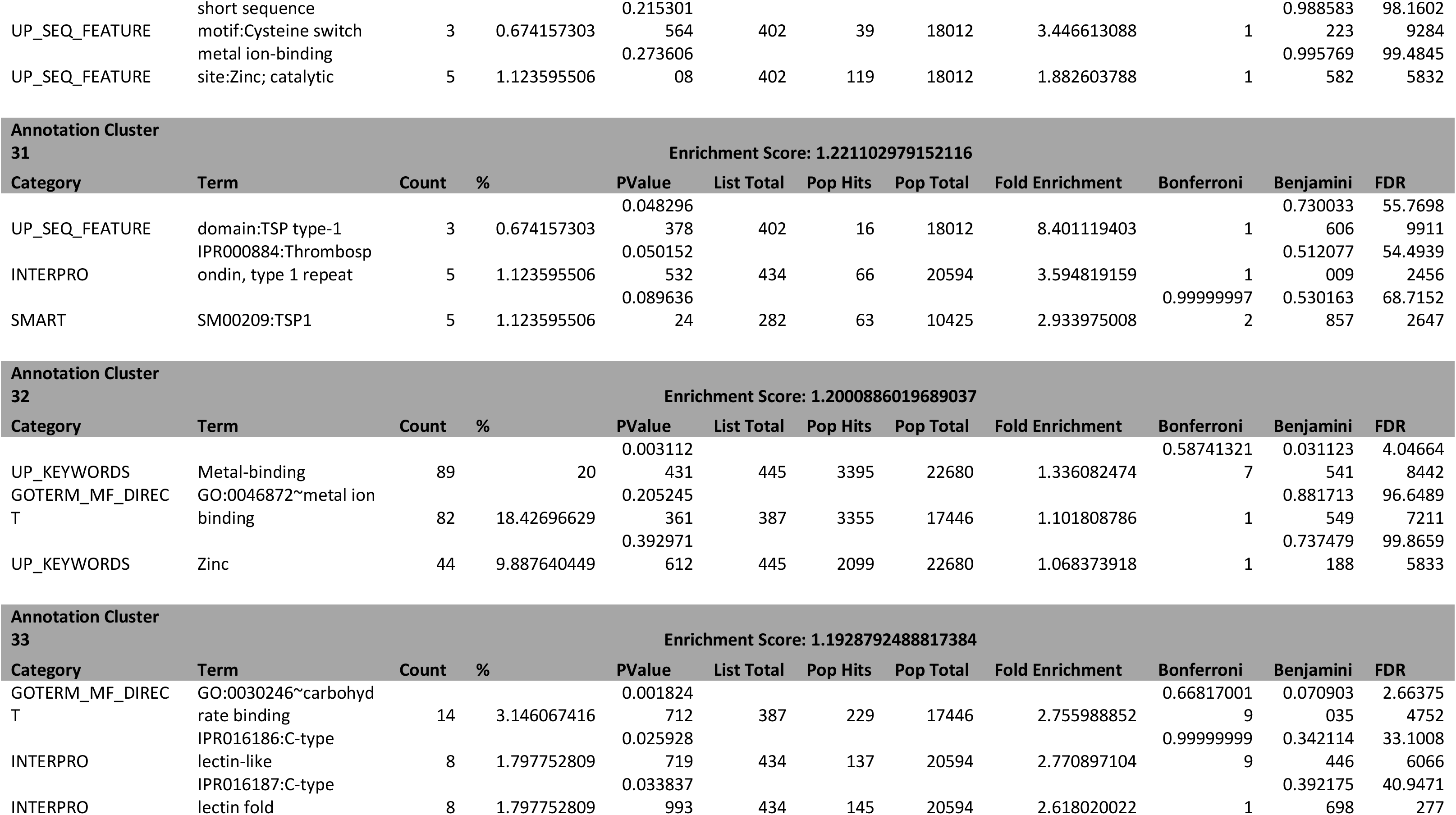
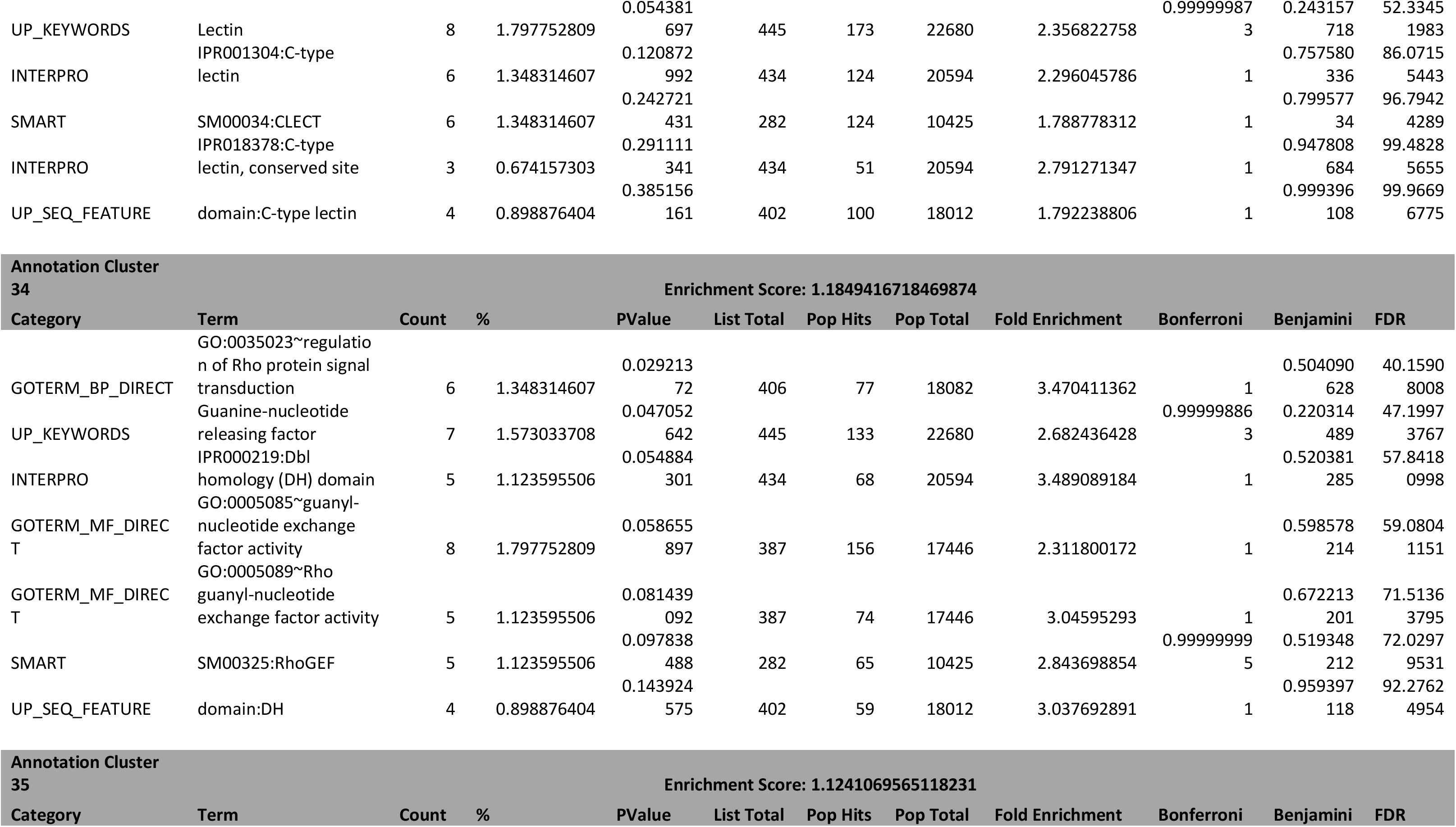
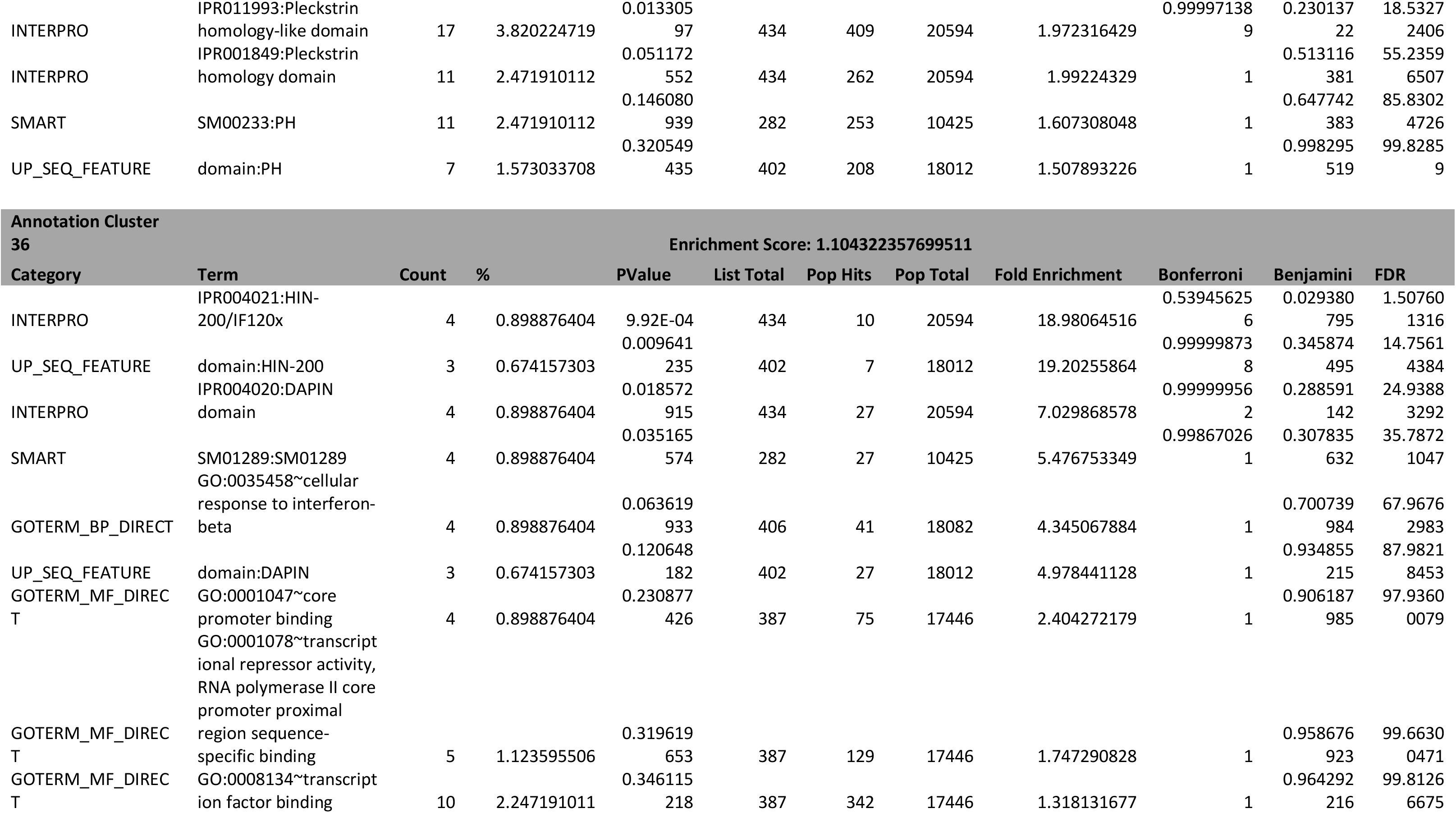
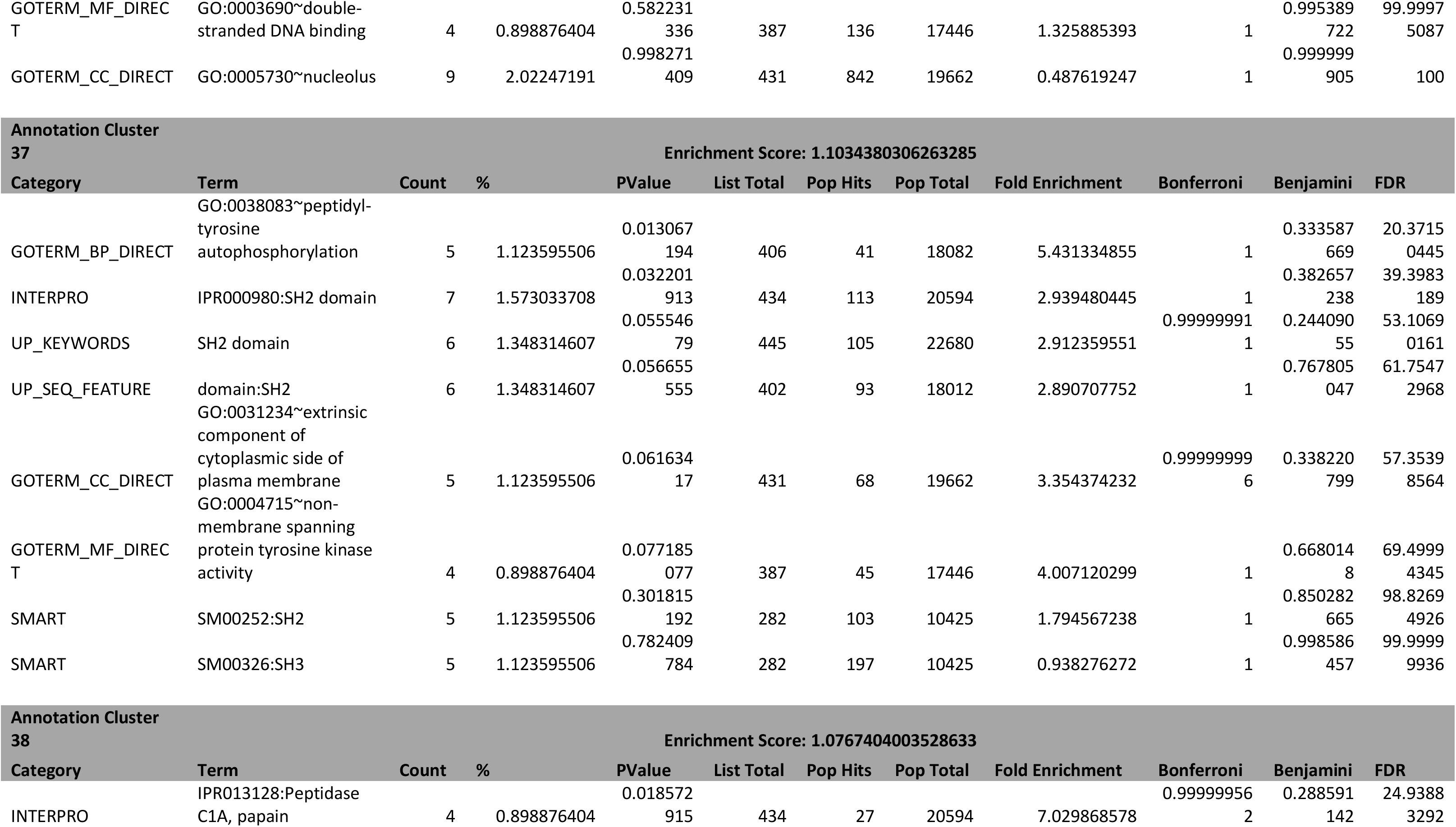
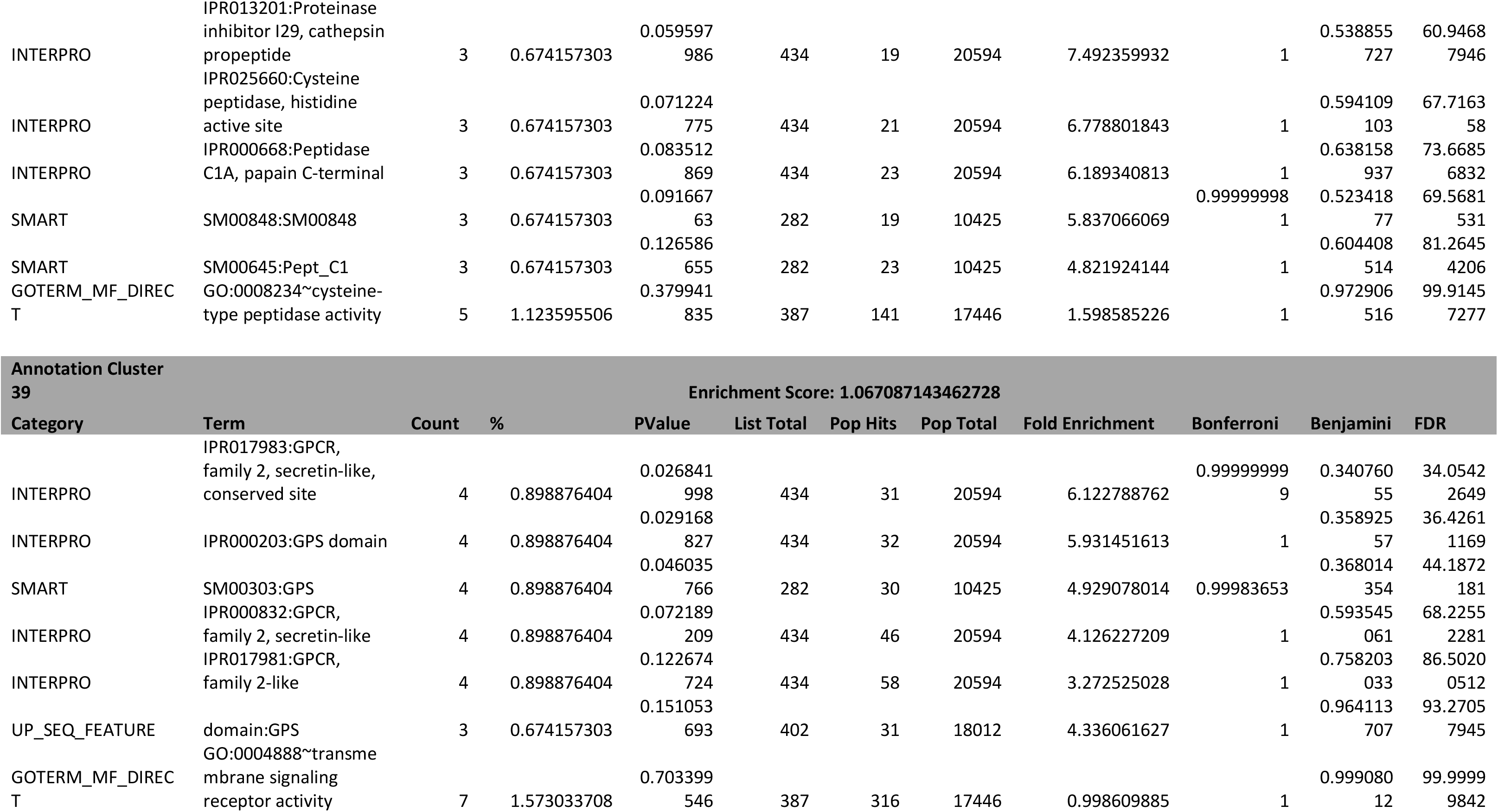
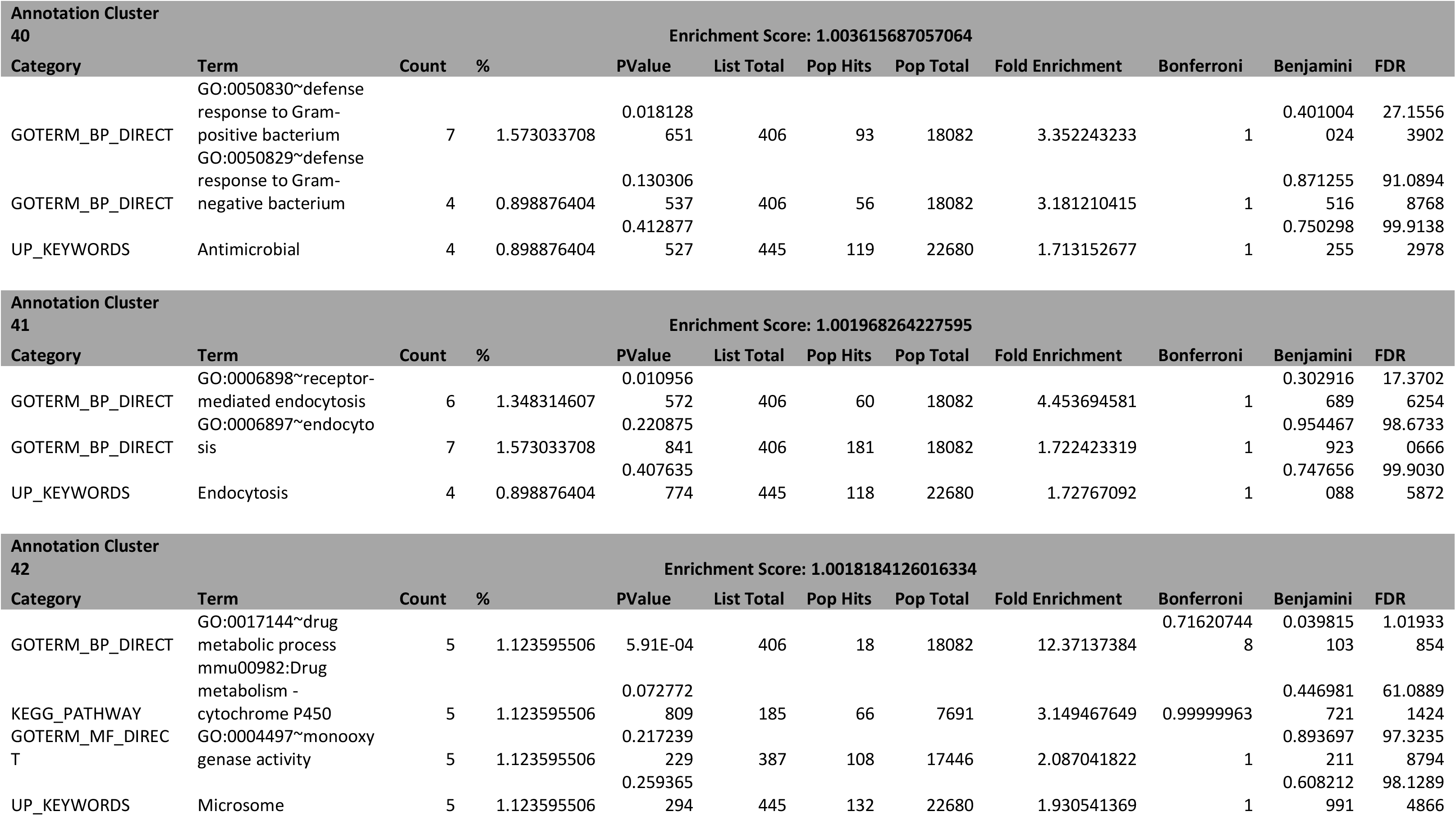
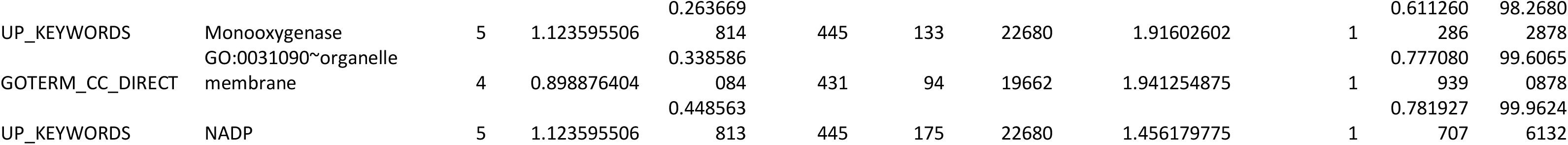
GO term analysis of genes not expressed in the myofiber but found in the whole muscle, defined as having an RPM value of at least 10 in the whole muscle and 0 in the single fiber. Used DAVID 6.8 online functional annotation tool with the ENSEMBL_GENE_ID identifier and with a classification stringency of medium.

**Table S2A:**
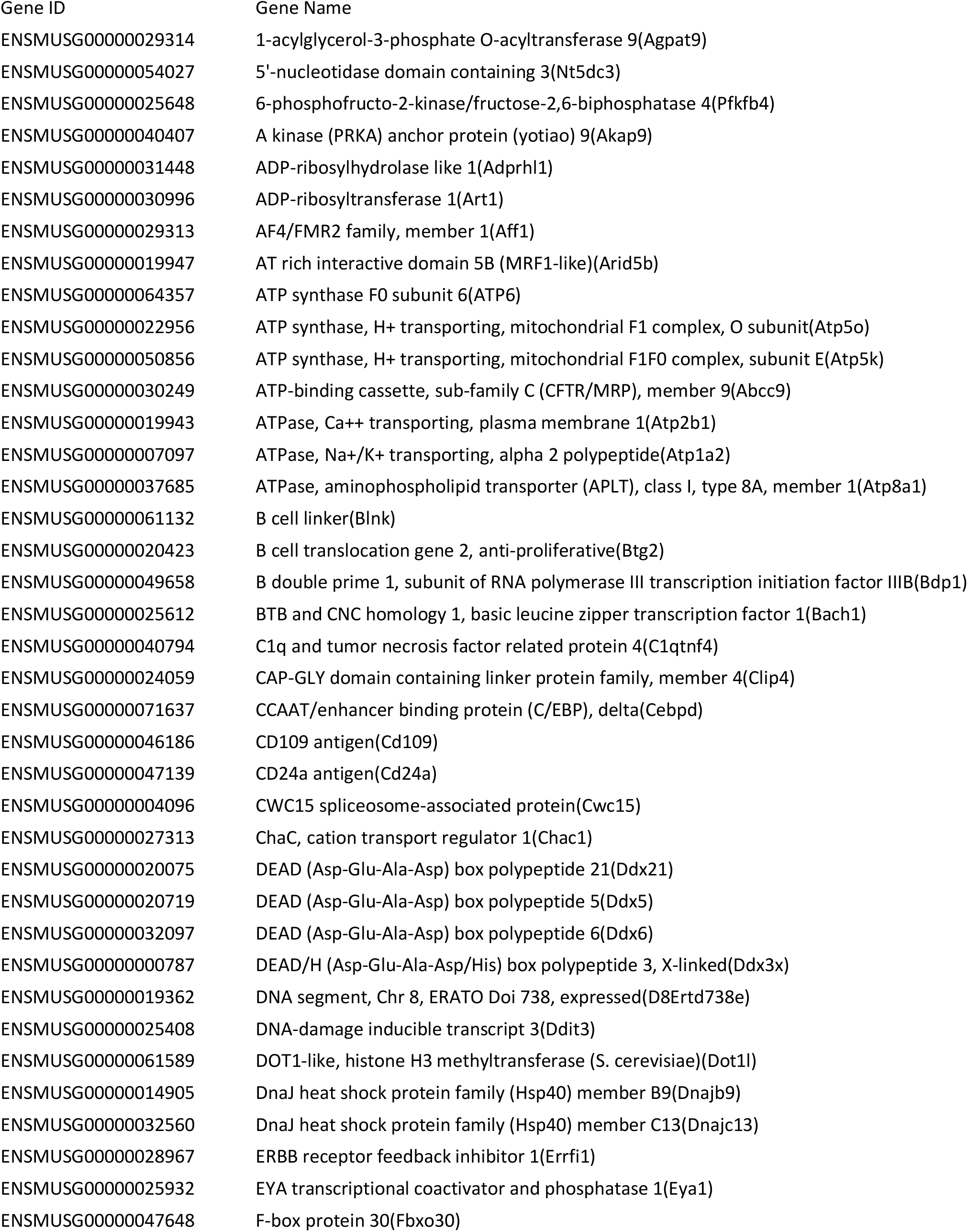
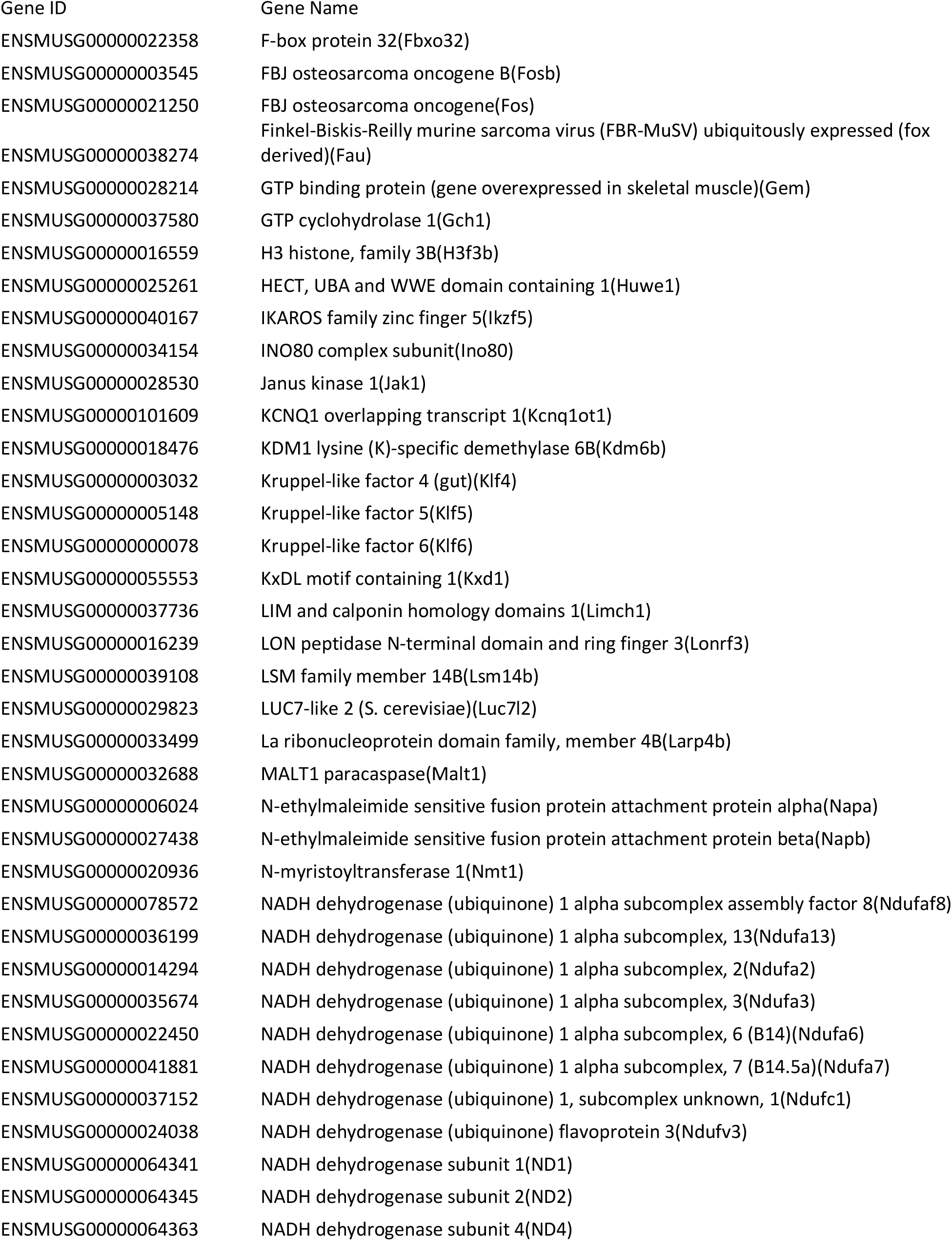
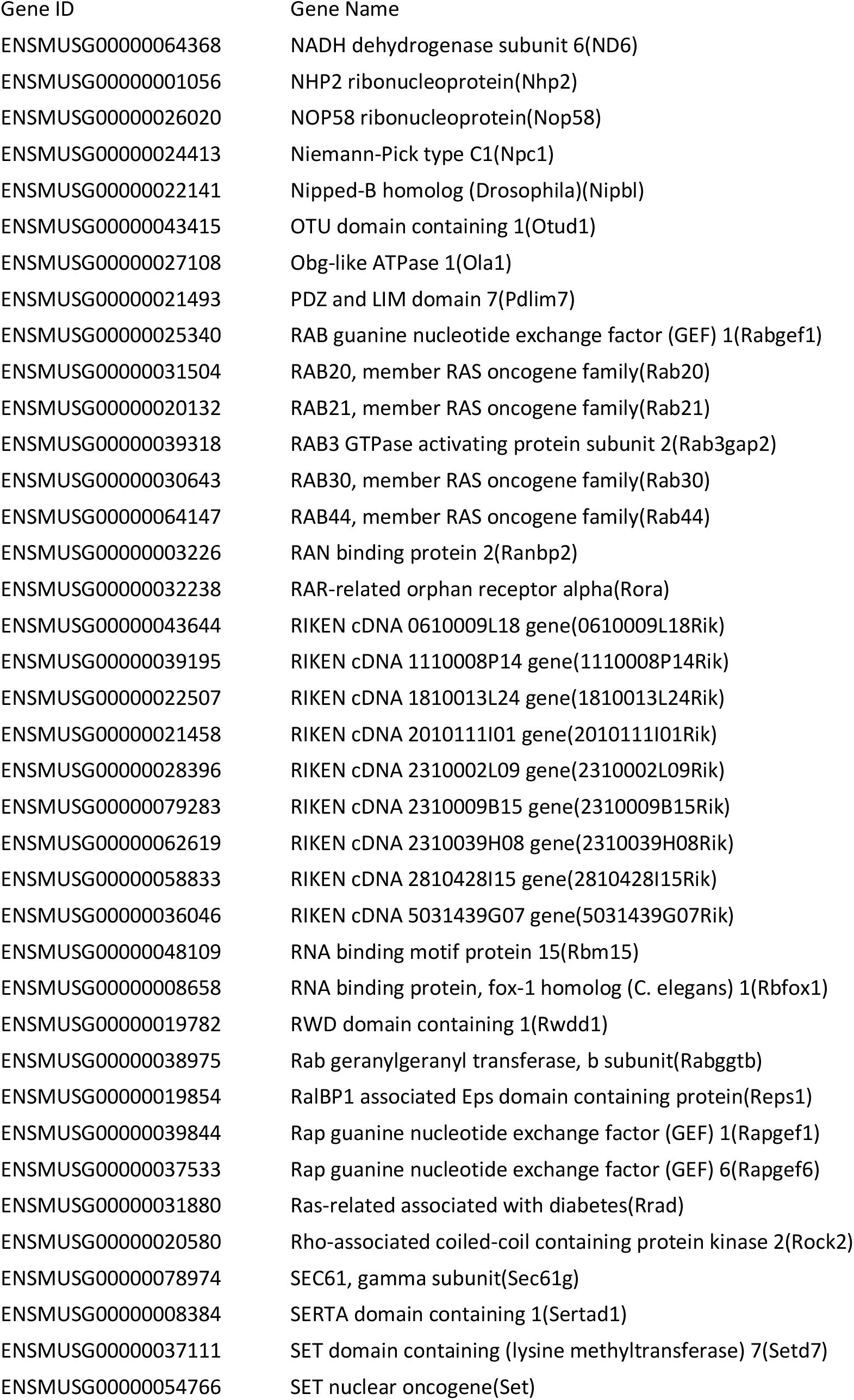
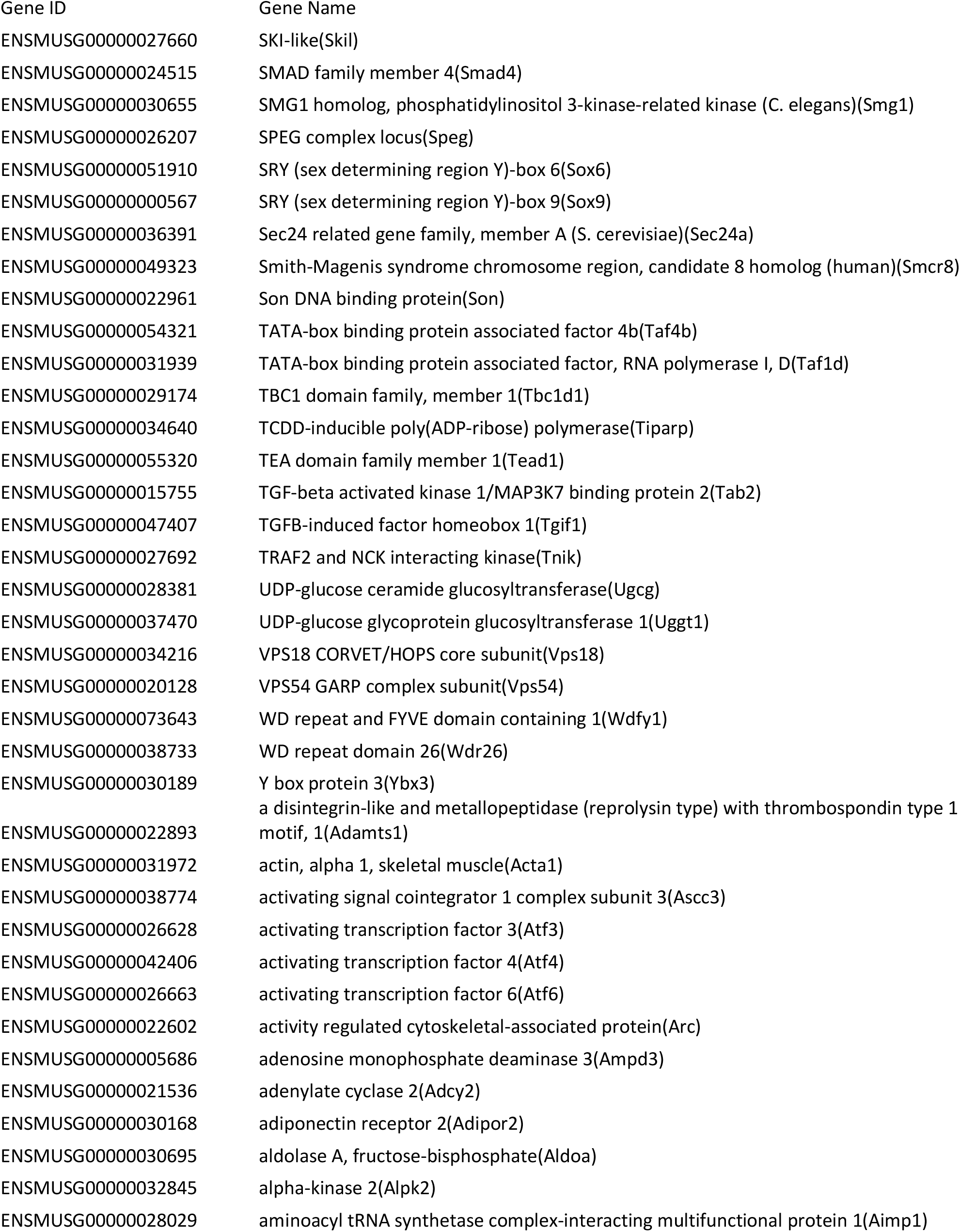
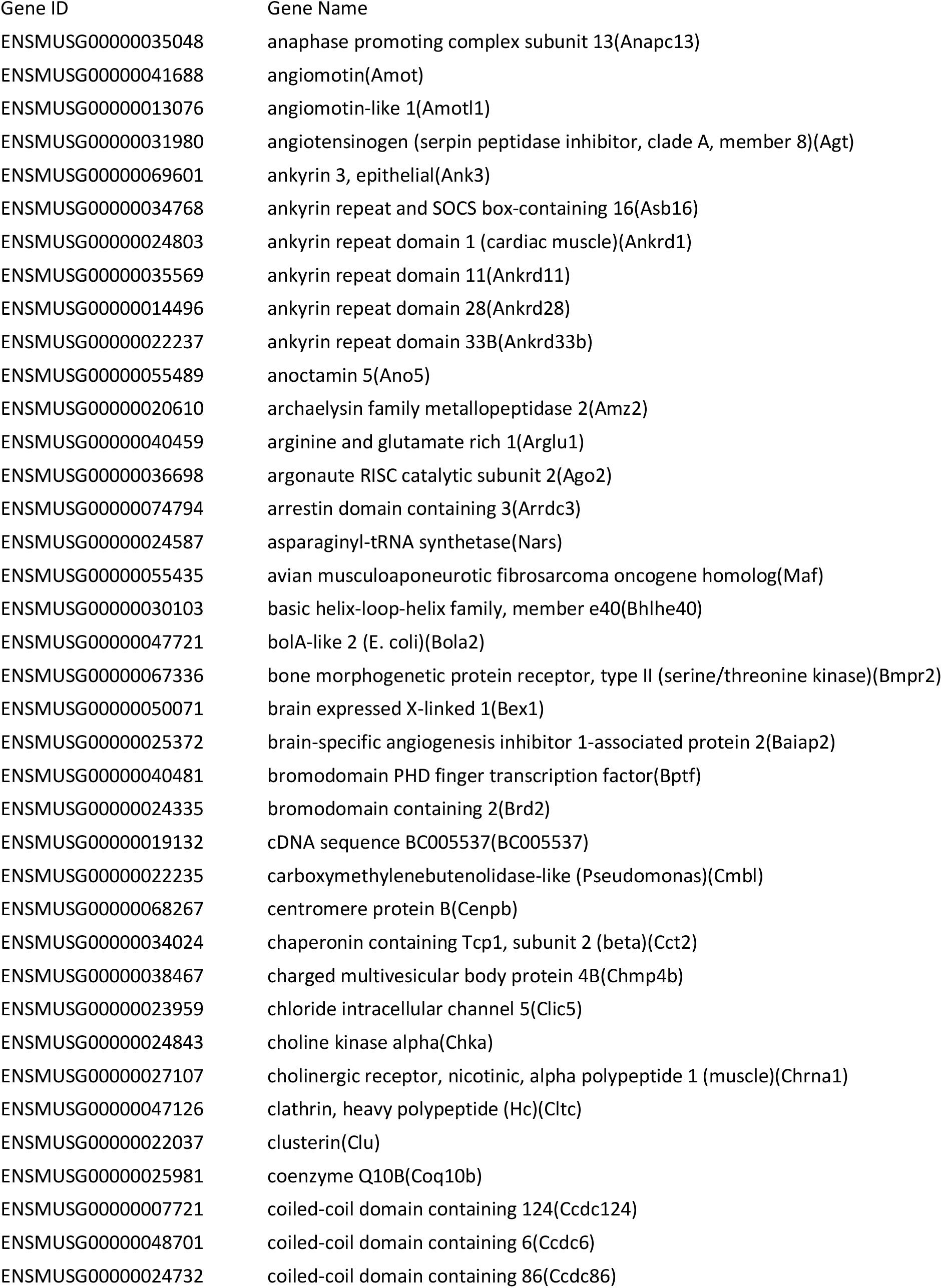
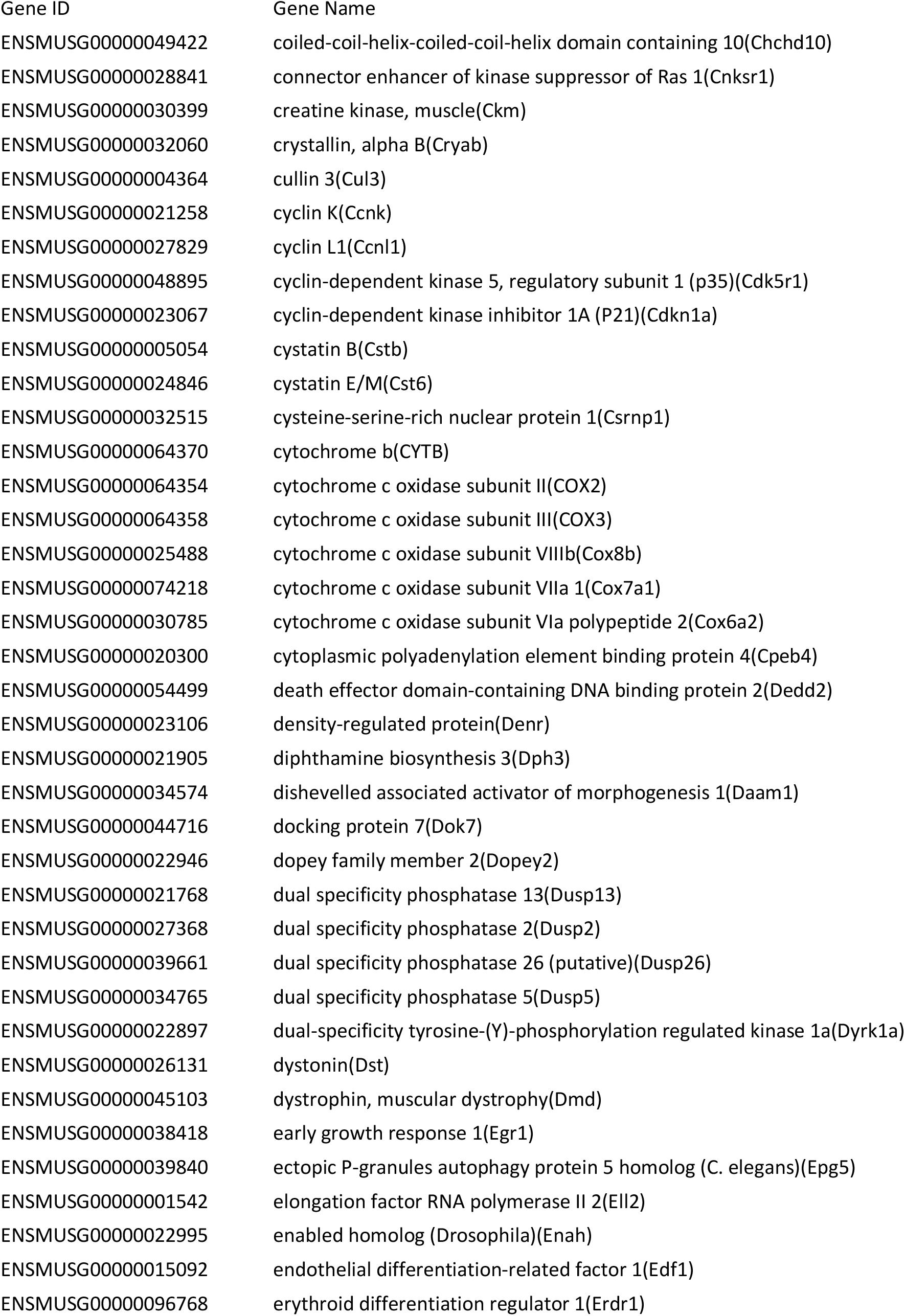
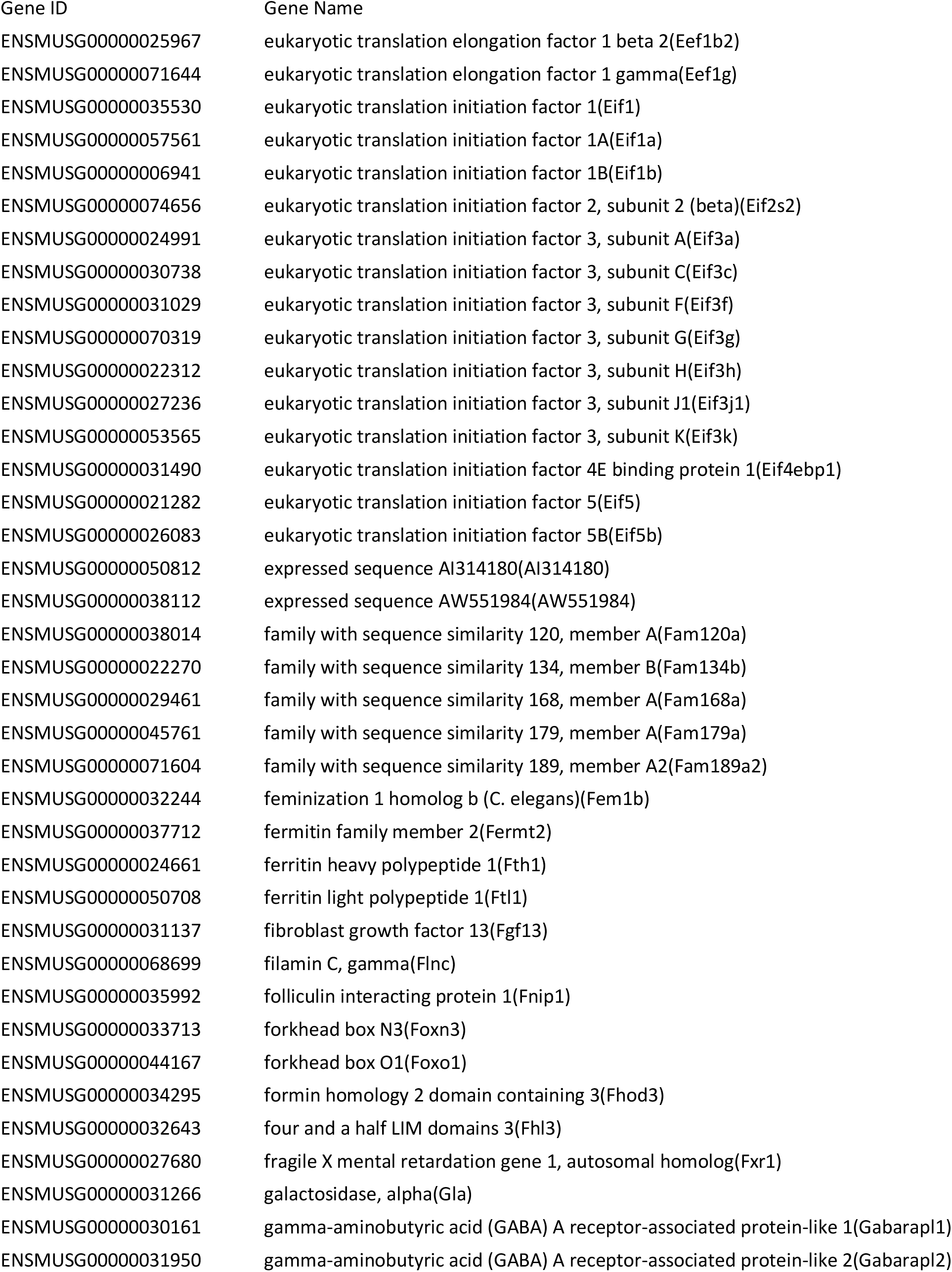
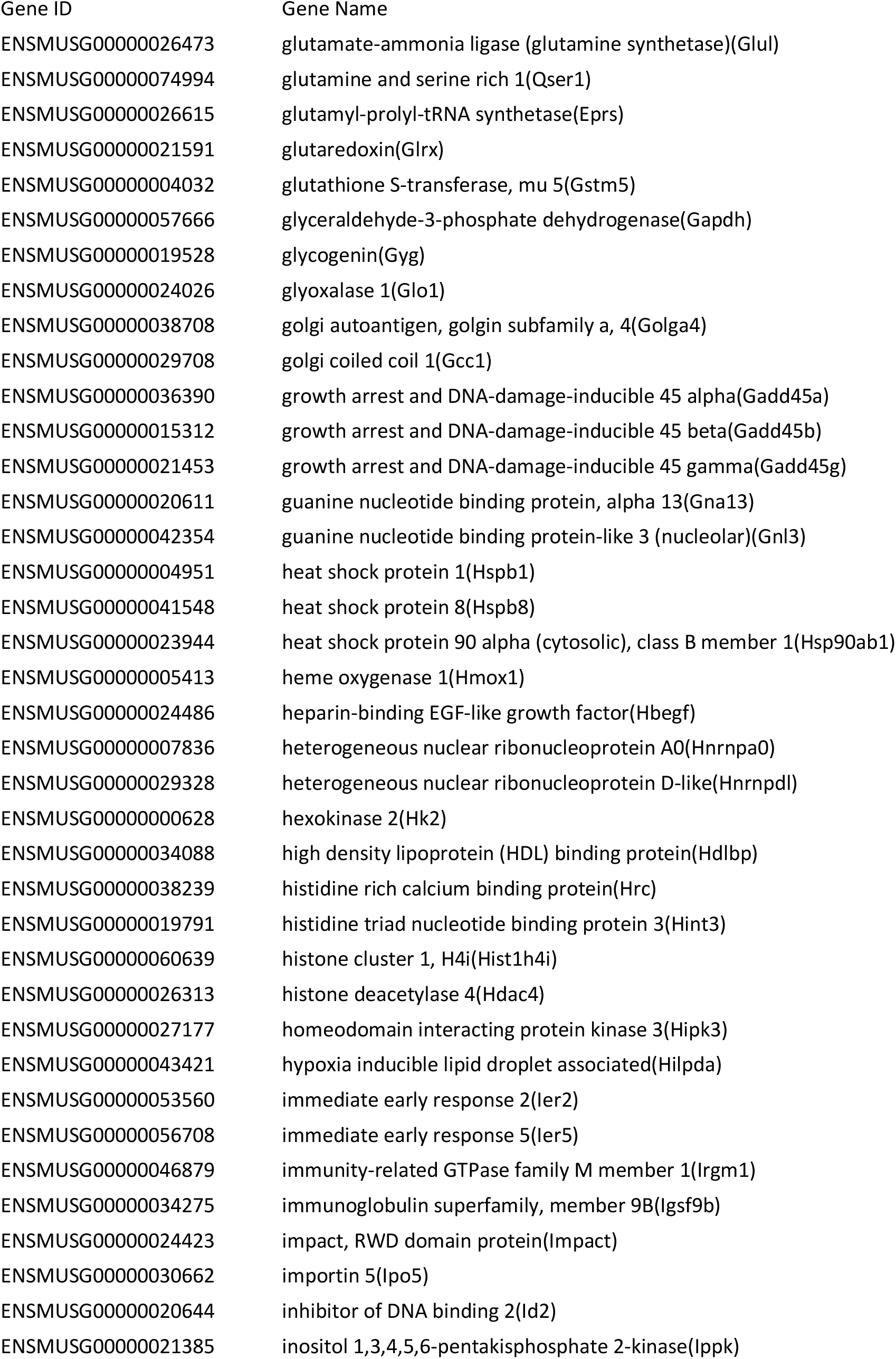
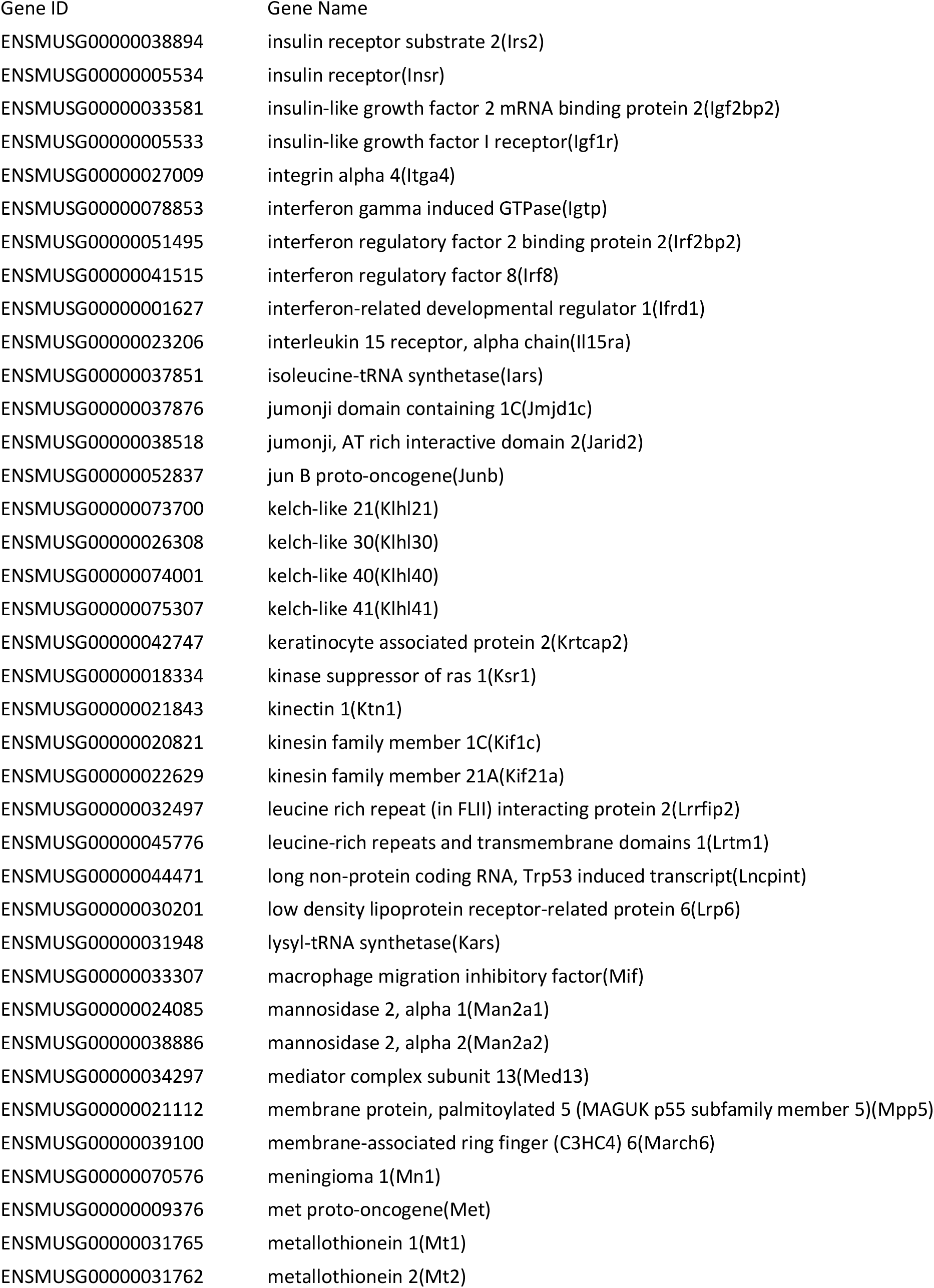
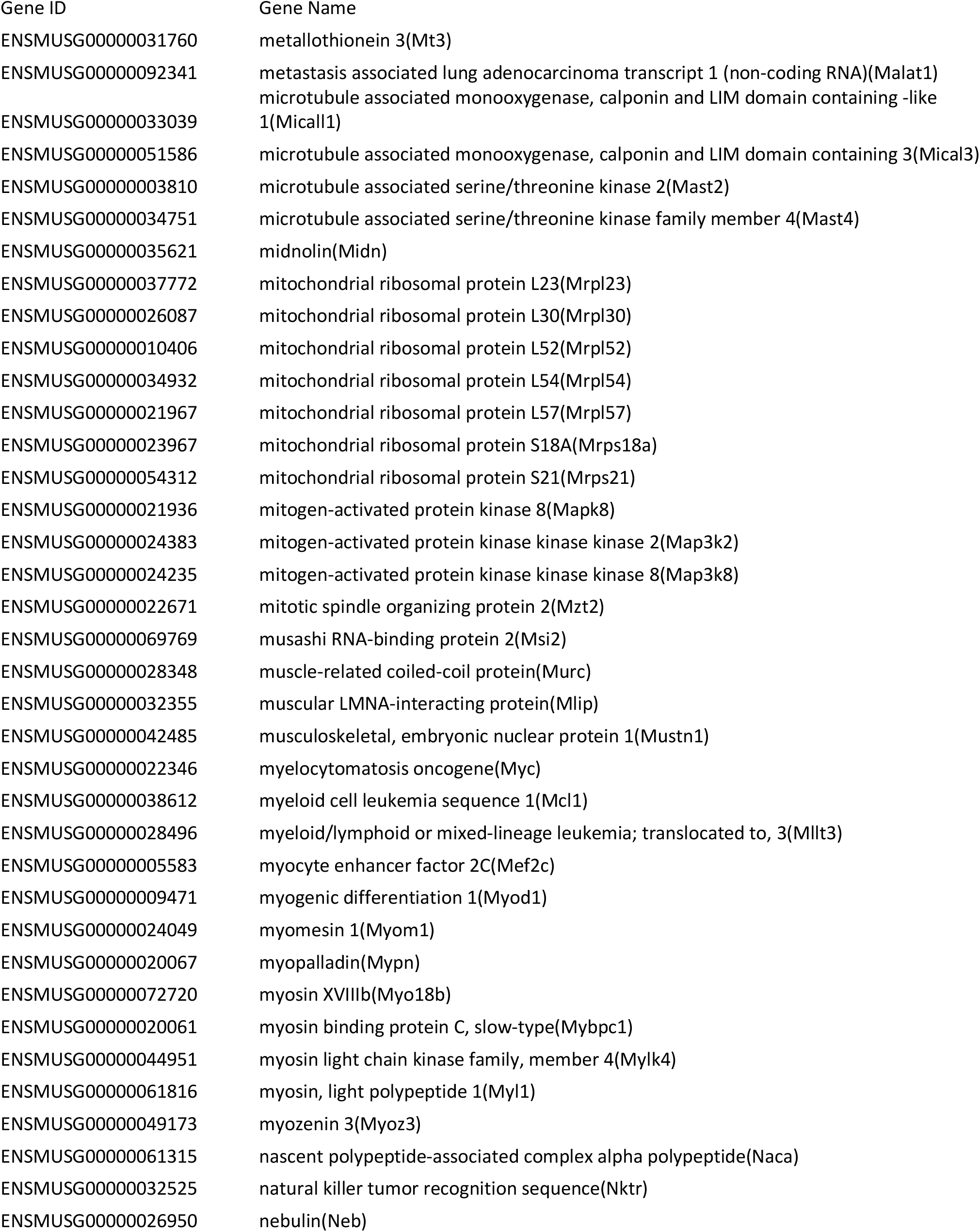
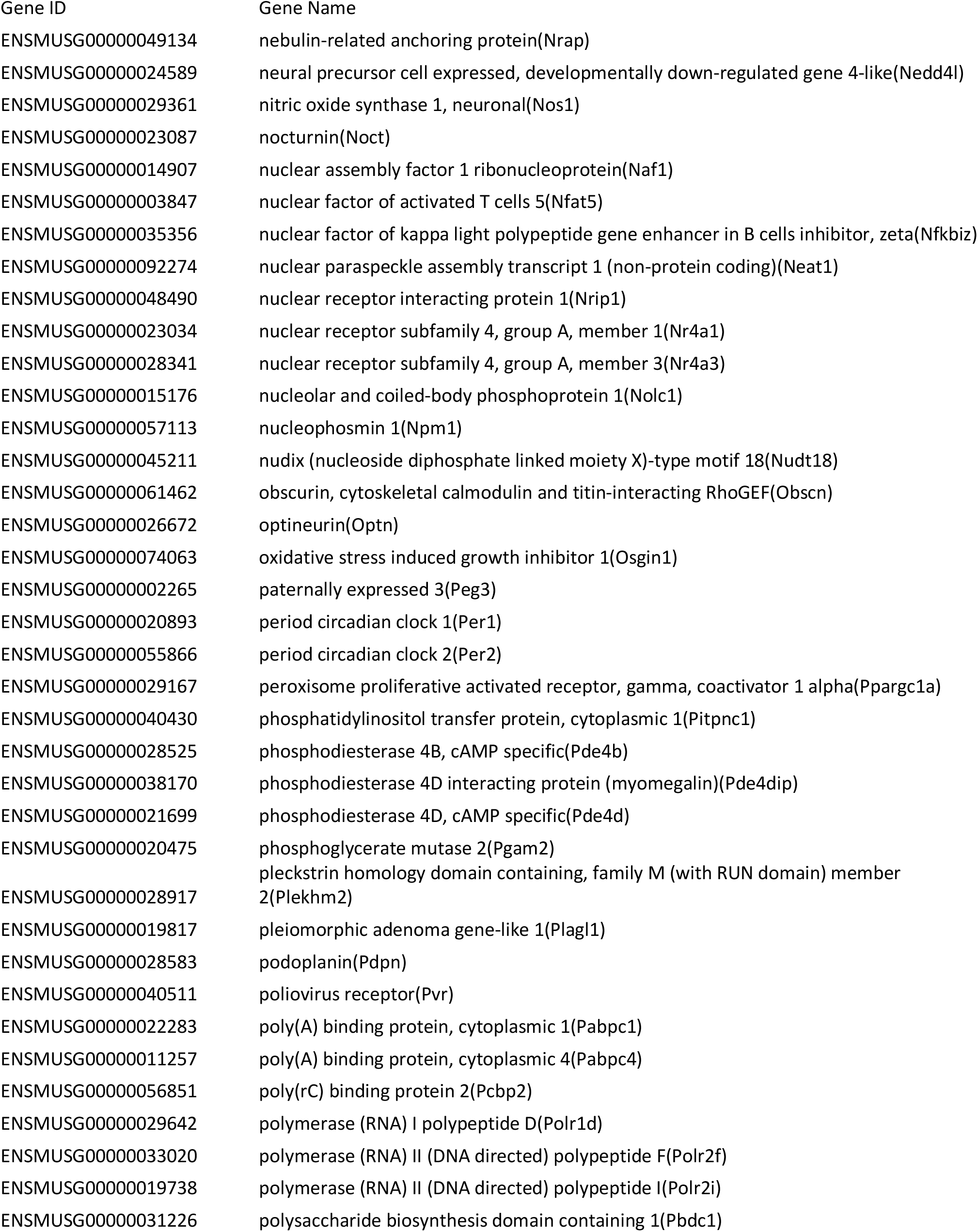
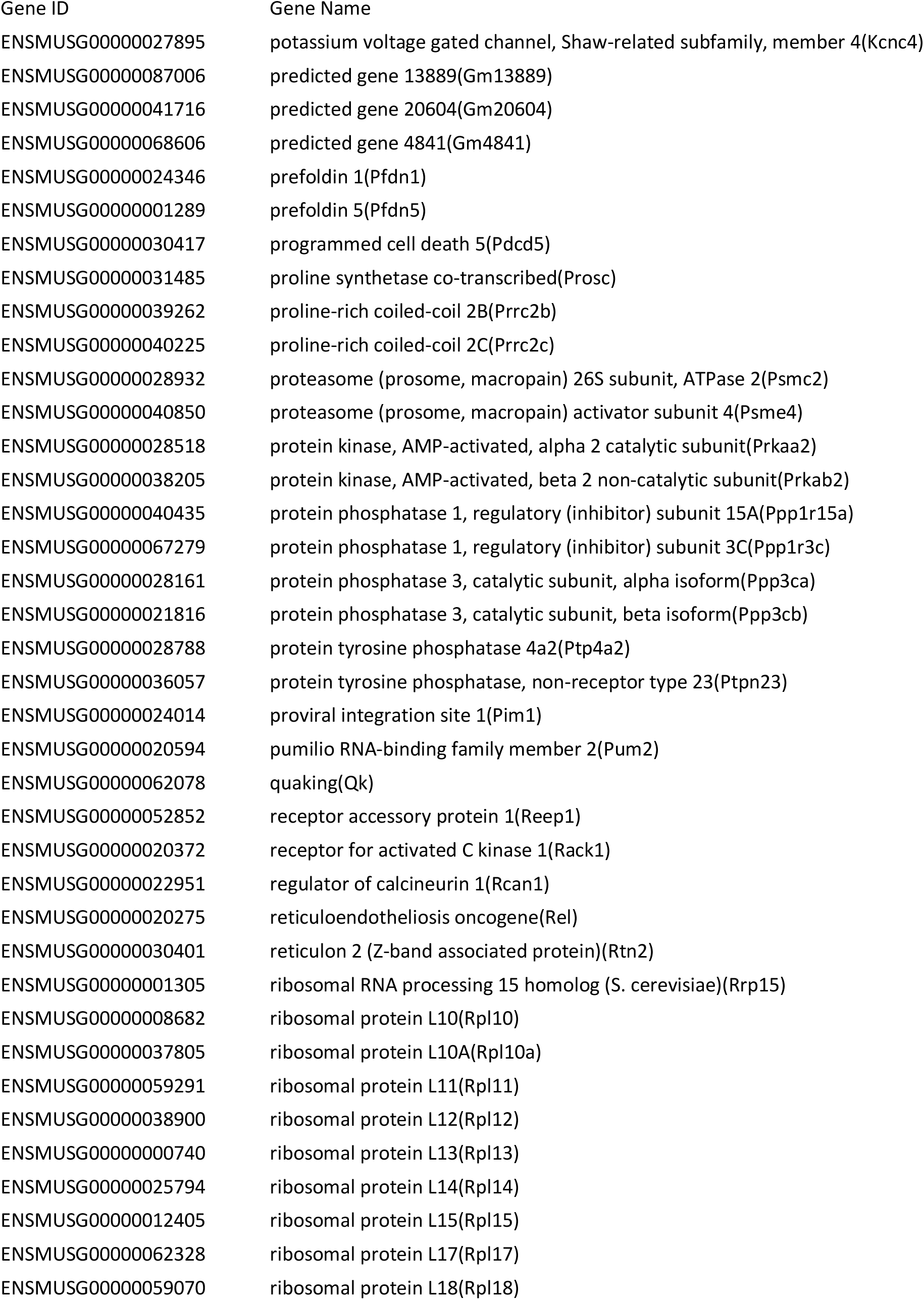
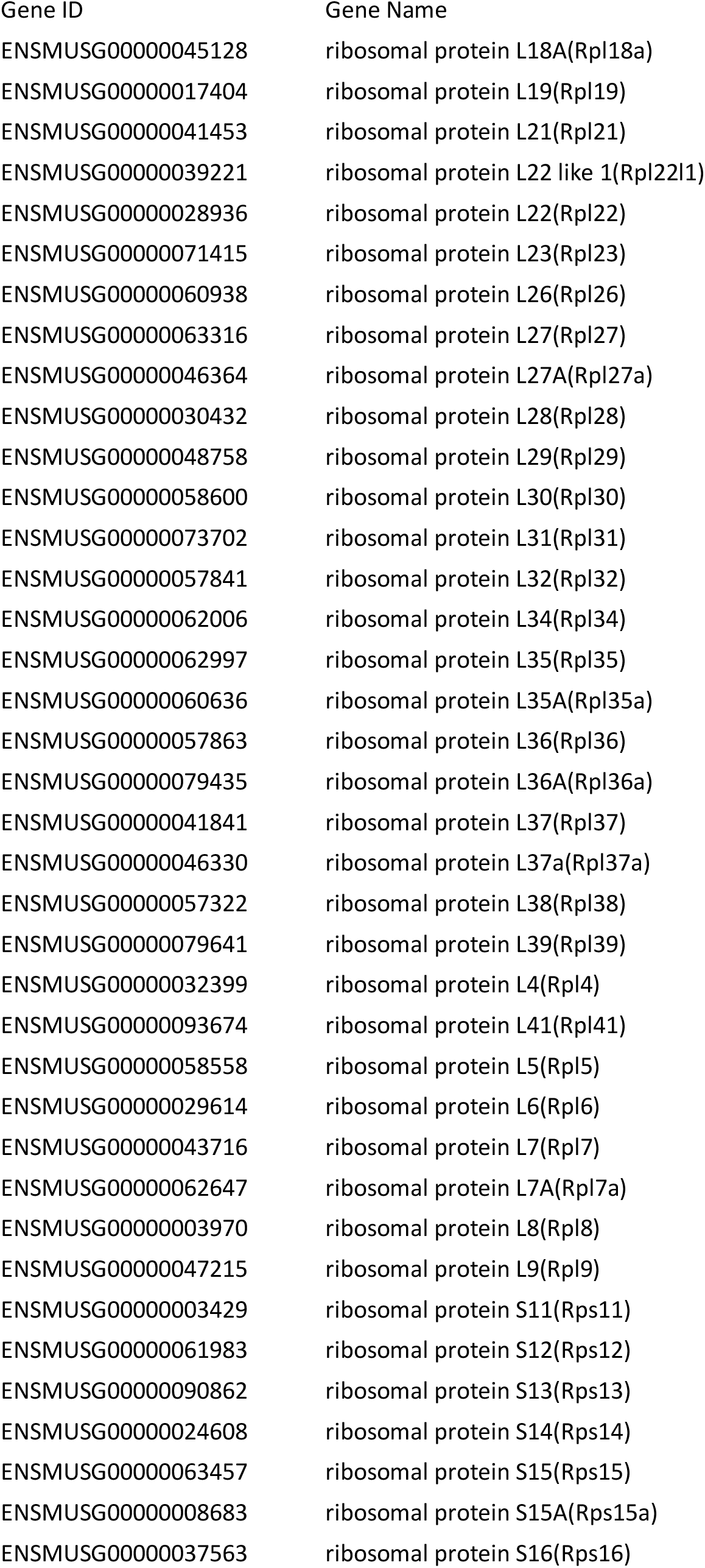
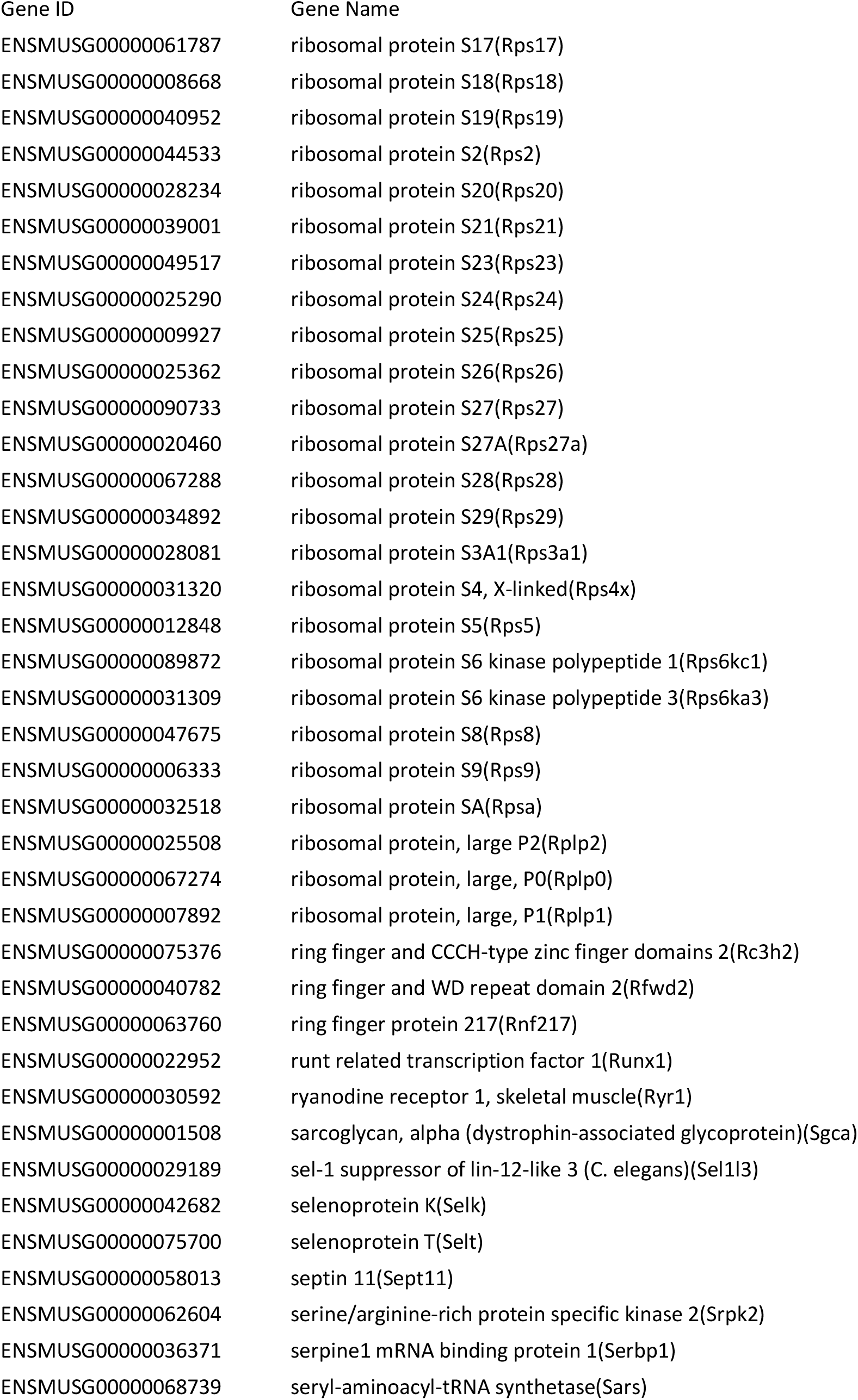
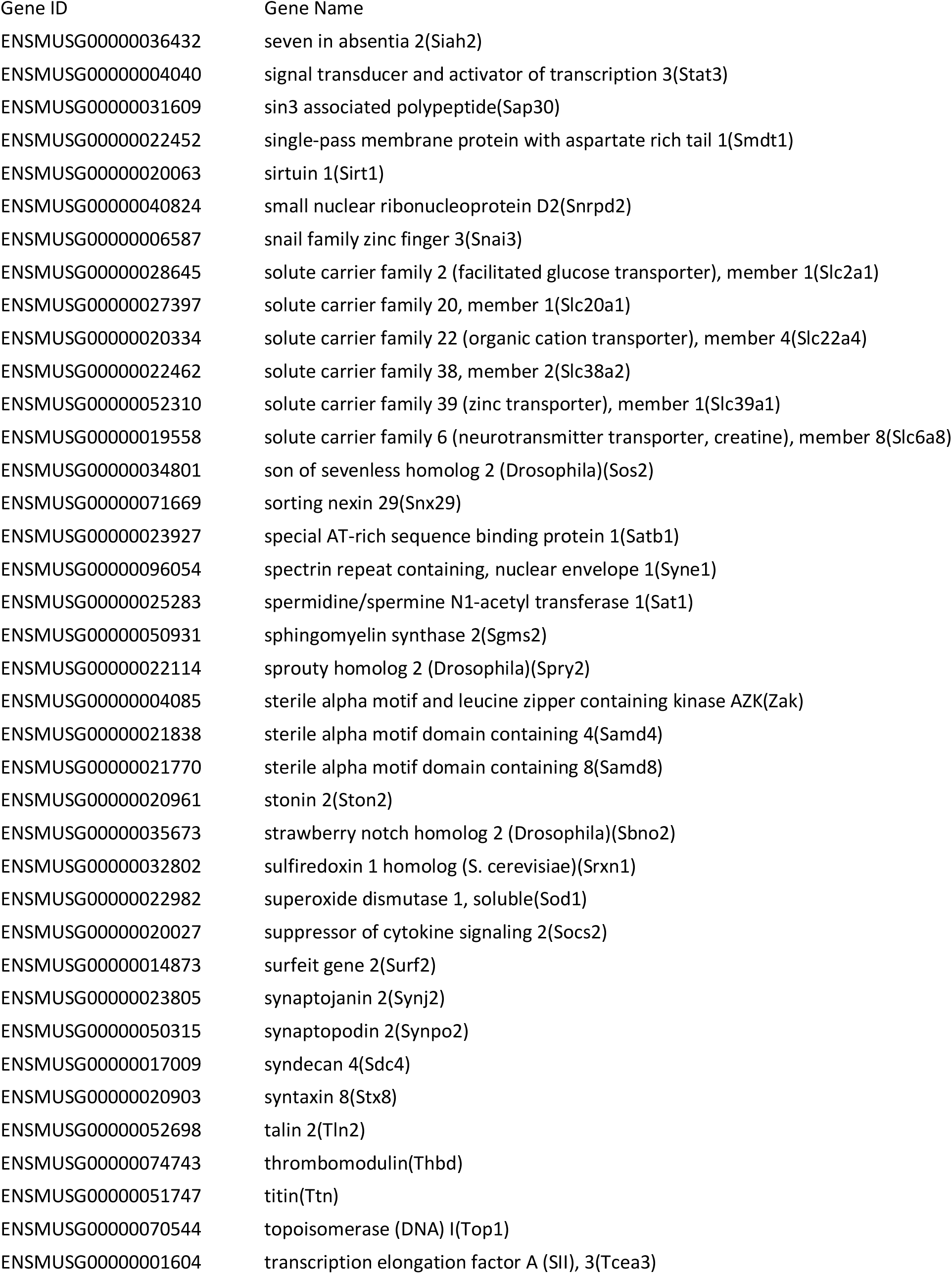
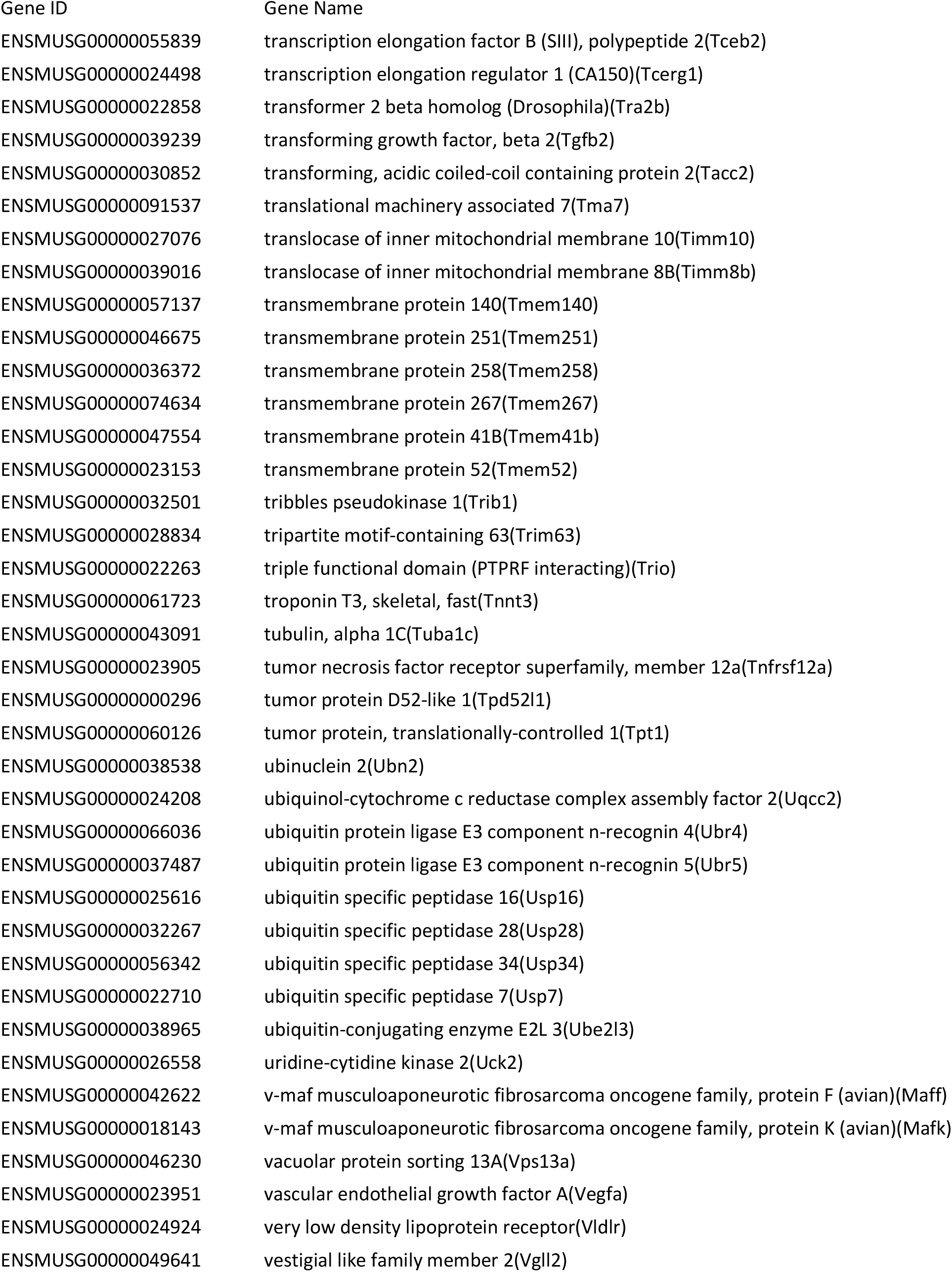
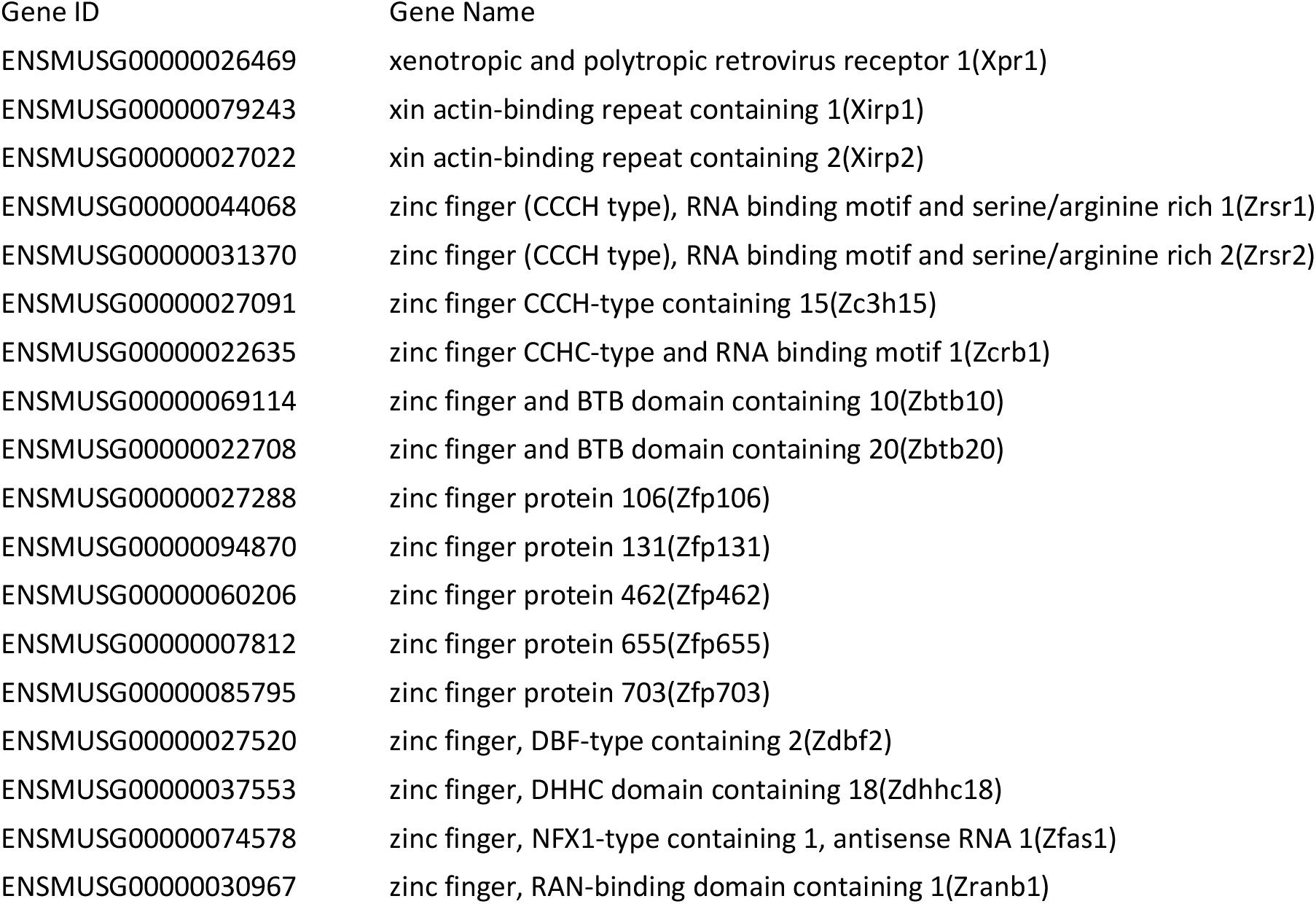
List of genes expressed exclusively in the myofiber, defined as genes with a q value lower than 0.01 between single fiber and whole muscle, more highly expressed in the single fiber with a Log2 fold change of less than -1, more highly expressed than 10 RPM and with a difference in expression between single fiber and whole muscle of at least 10 RPM.

**Table S2B:**
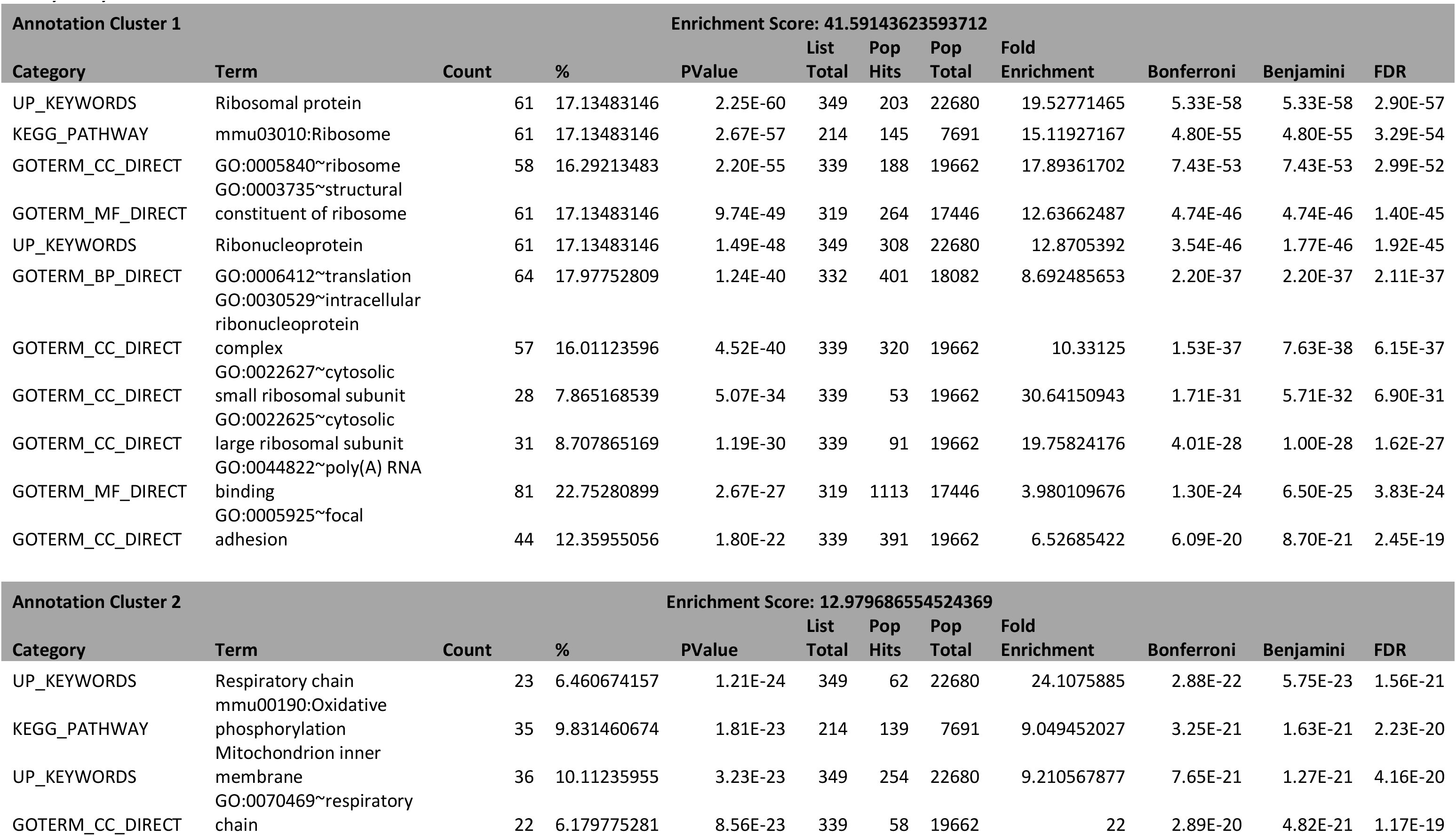
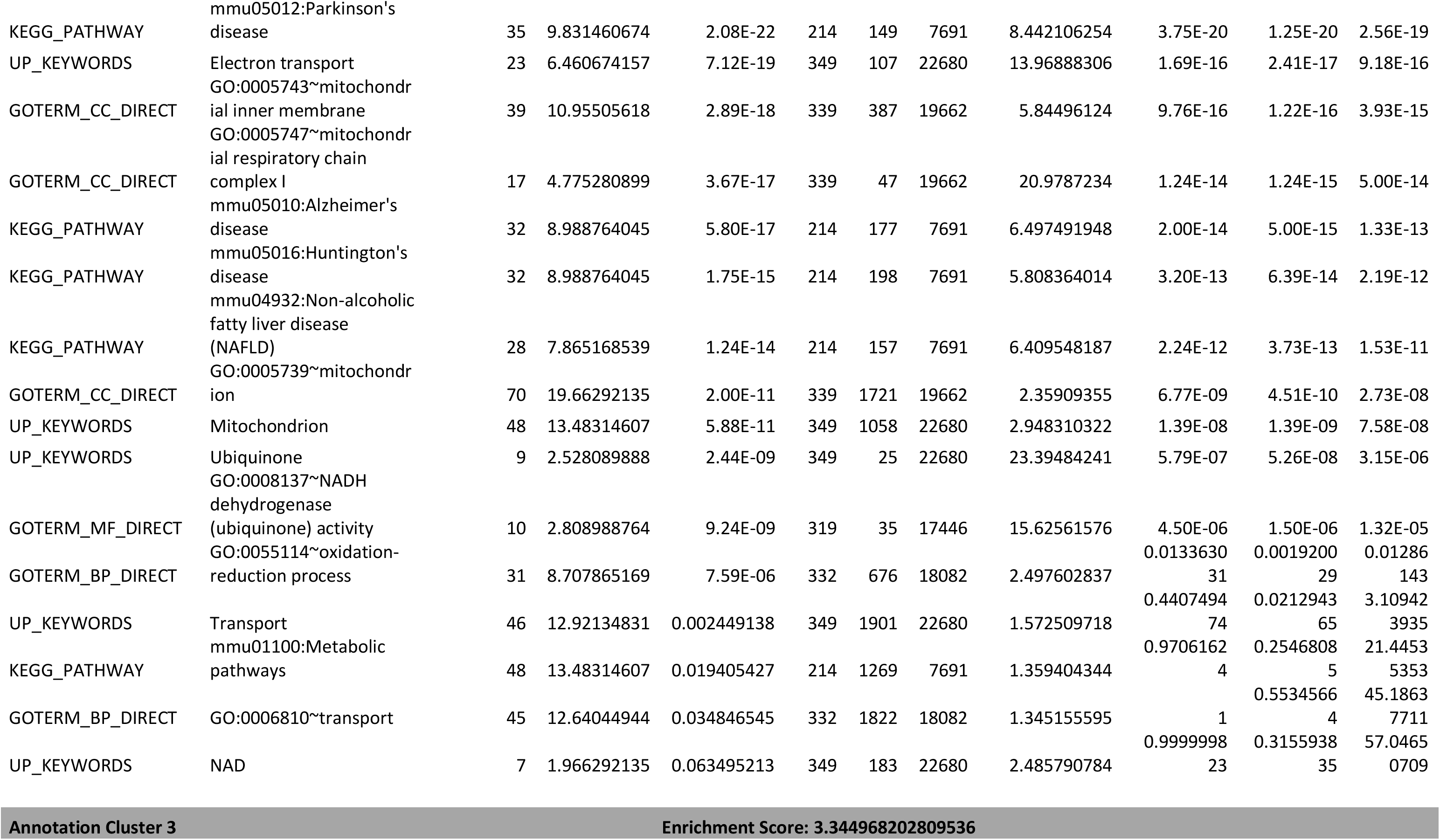
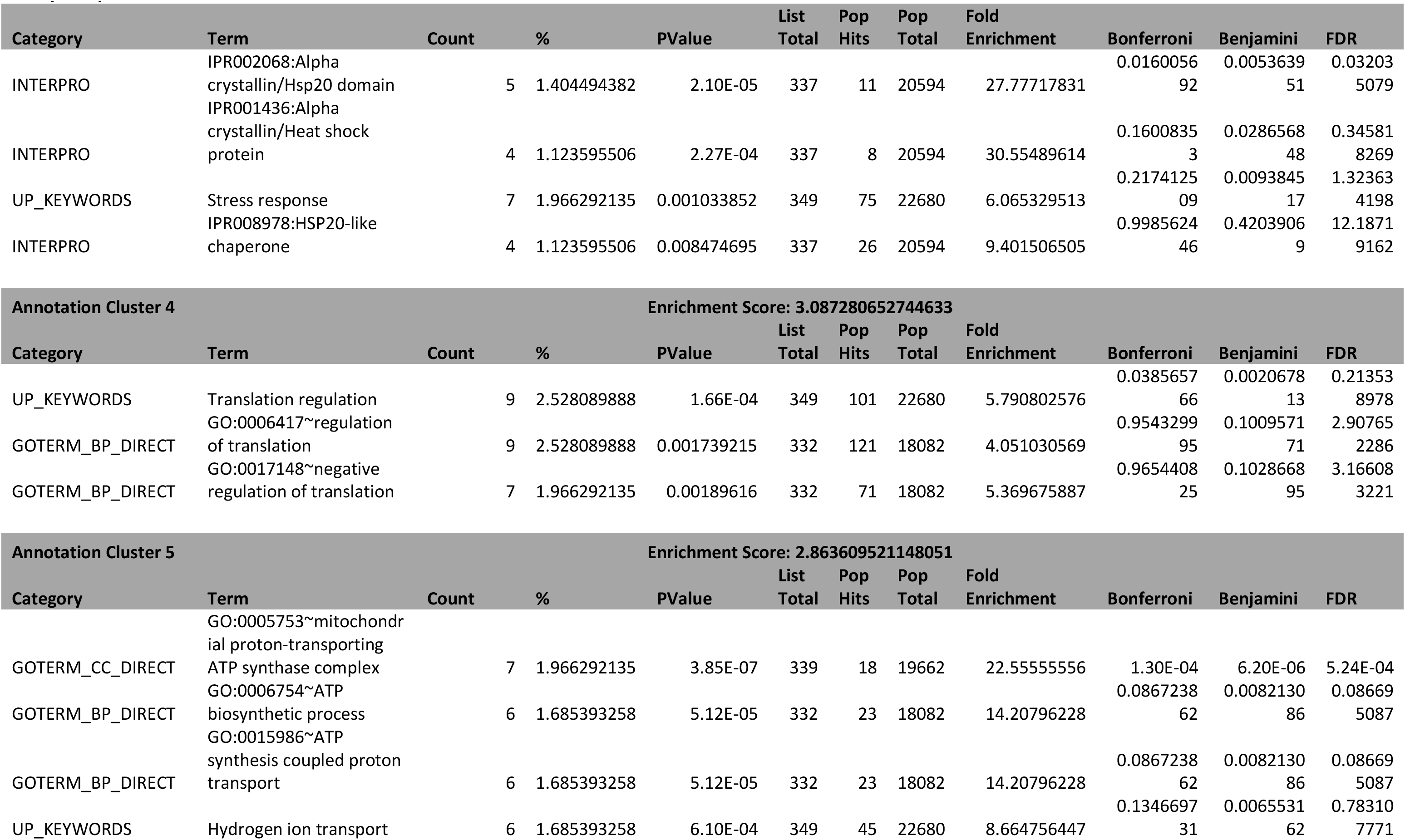
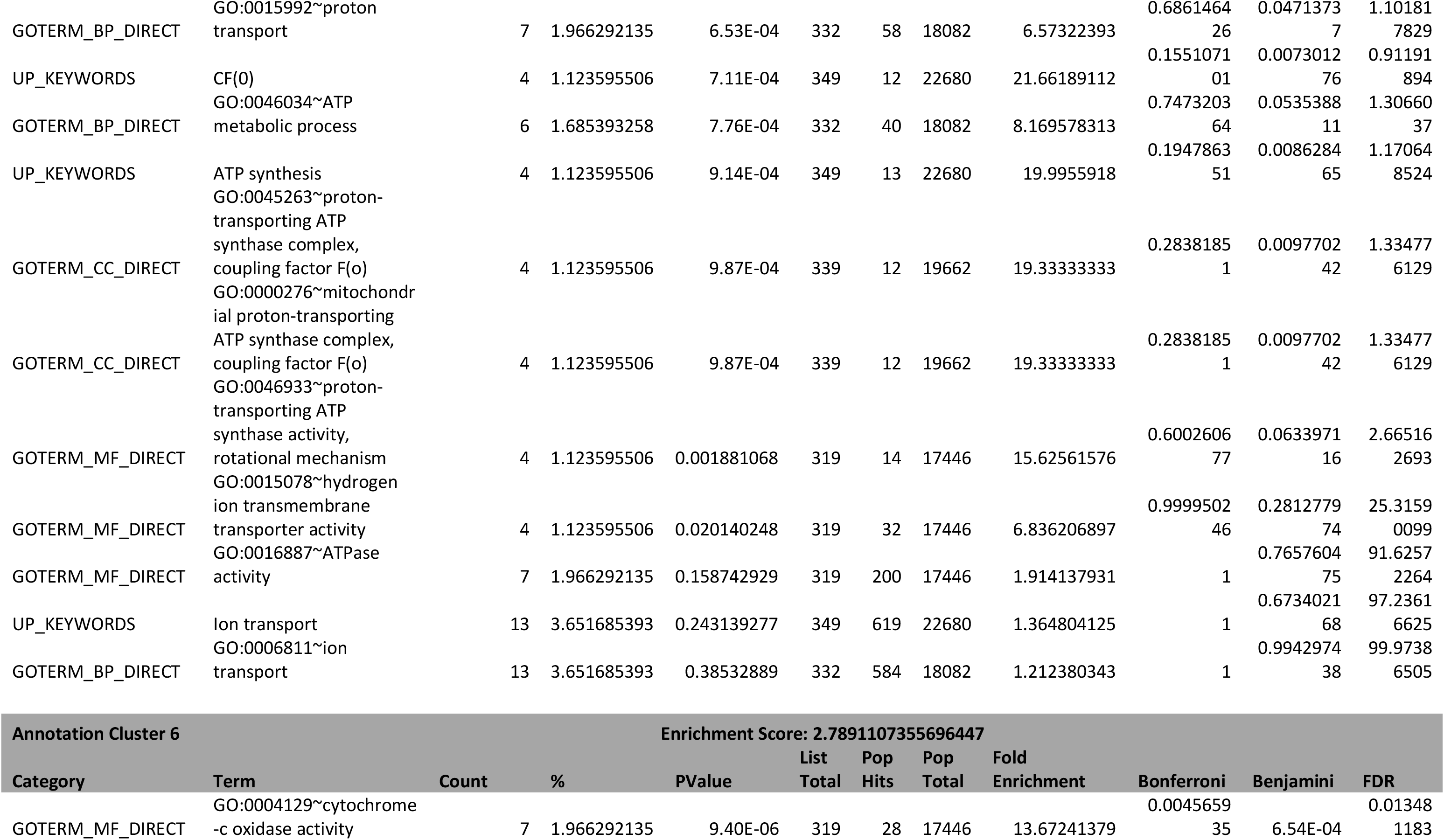
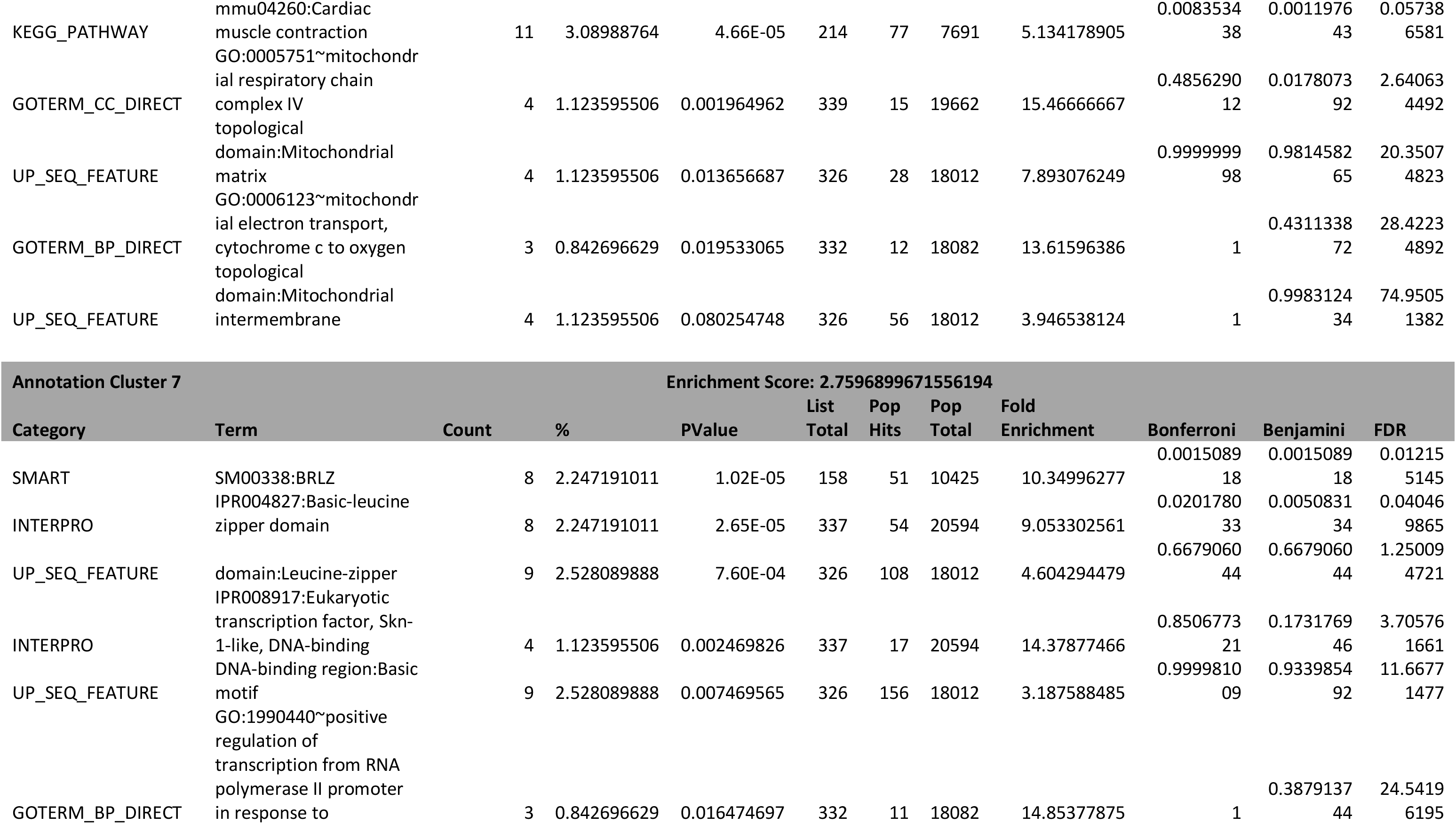
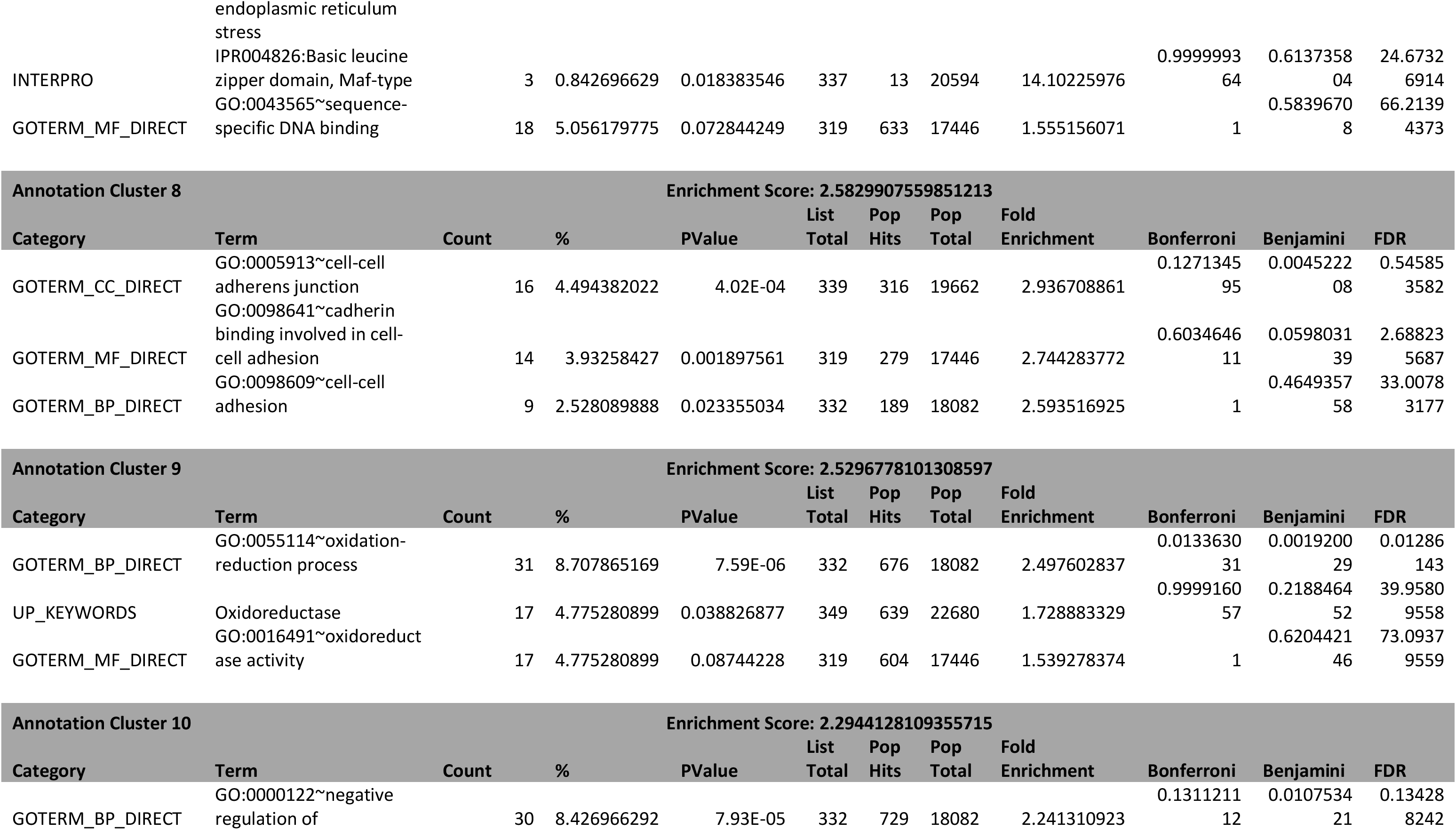
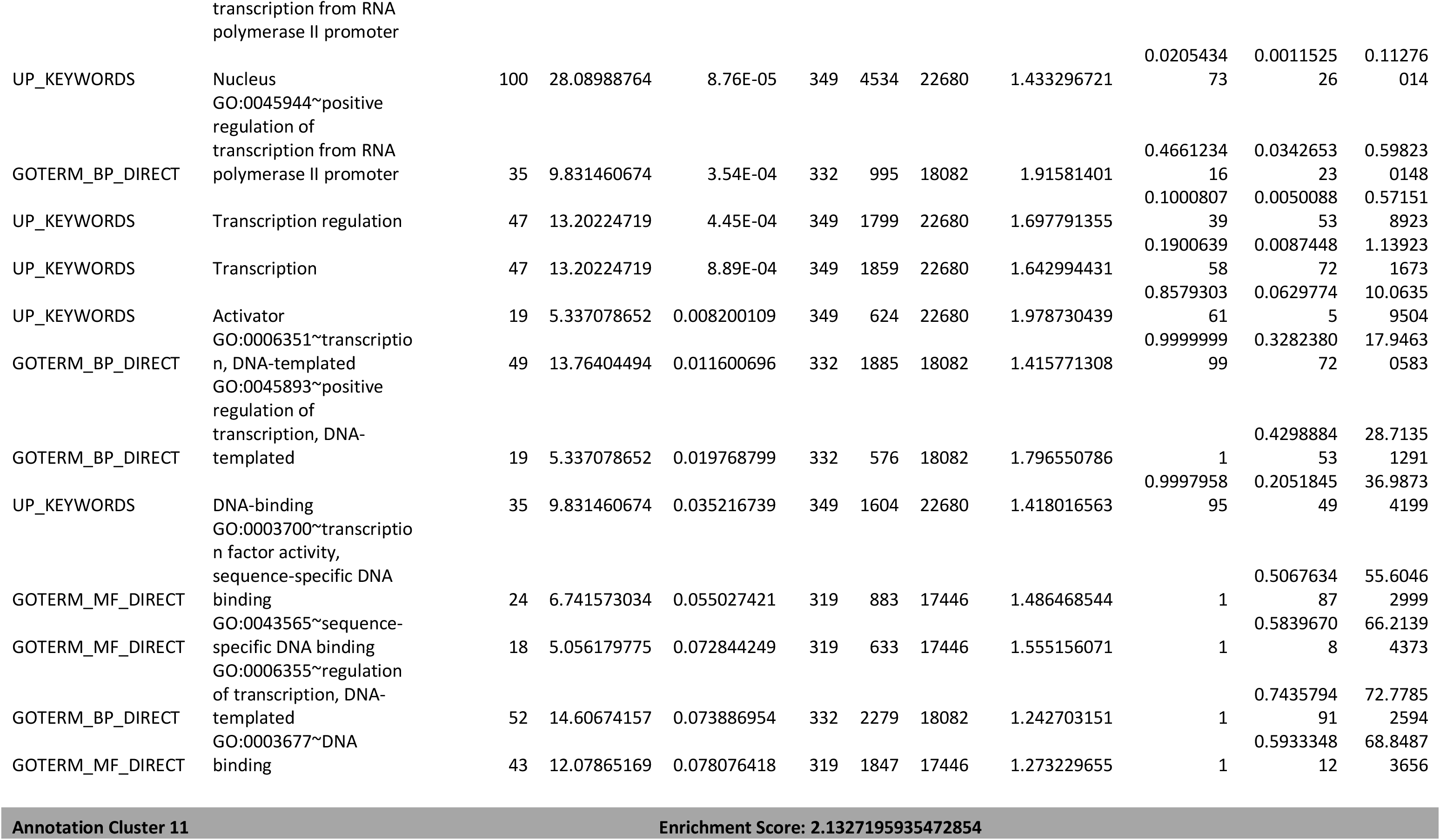
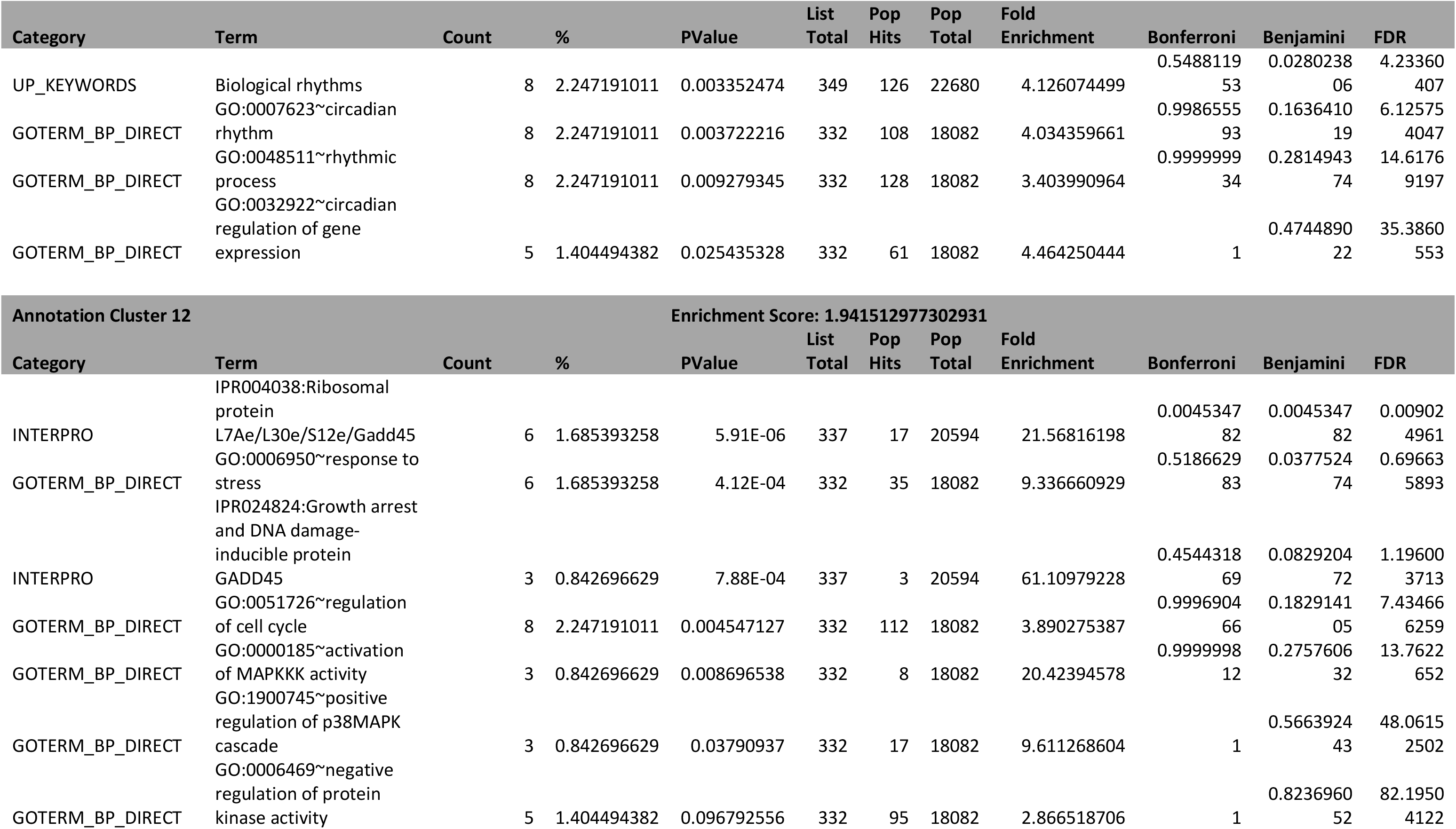
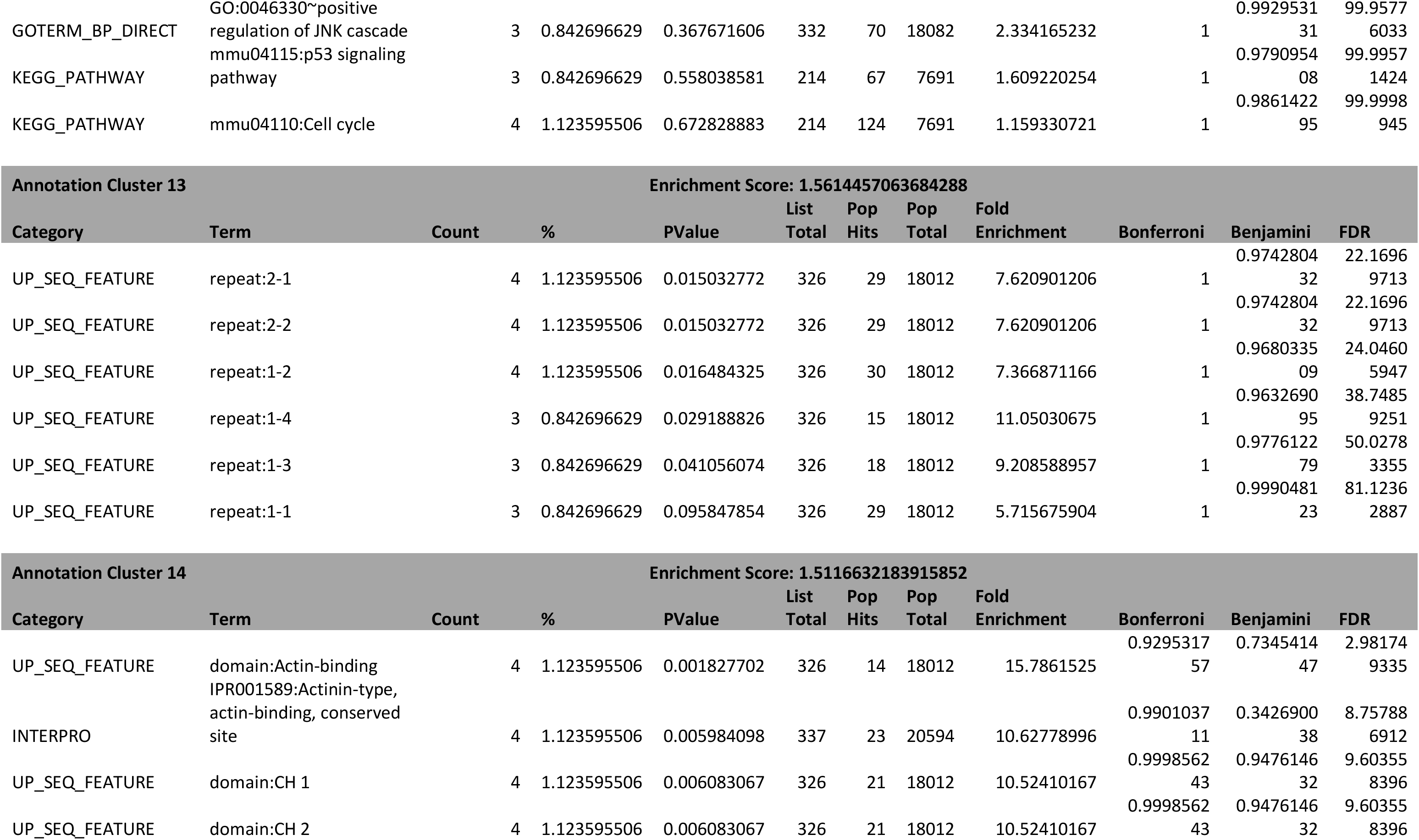
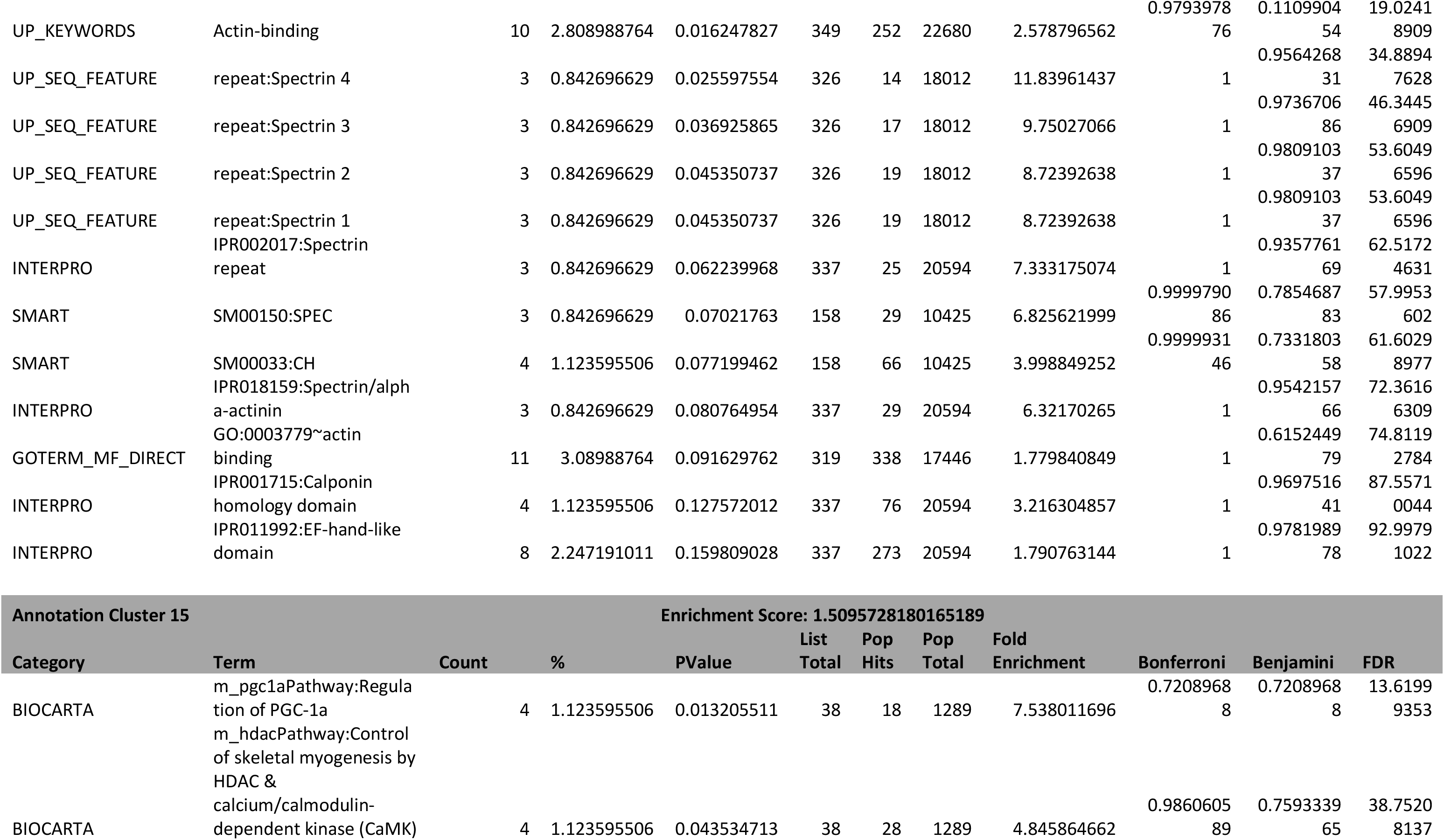
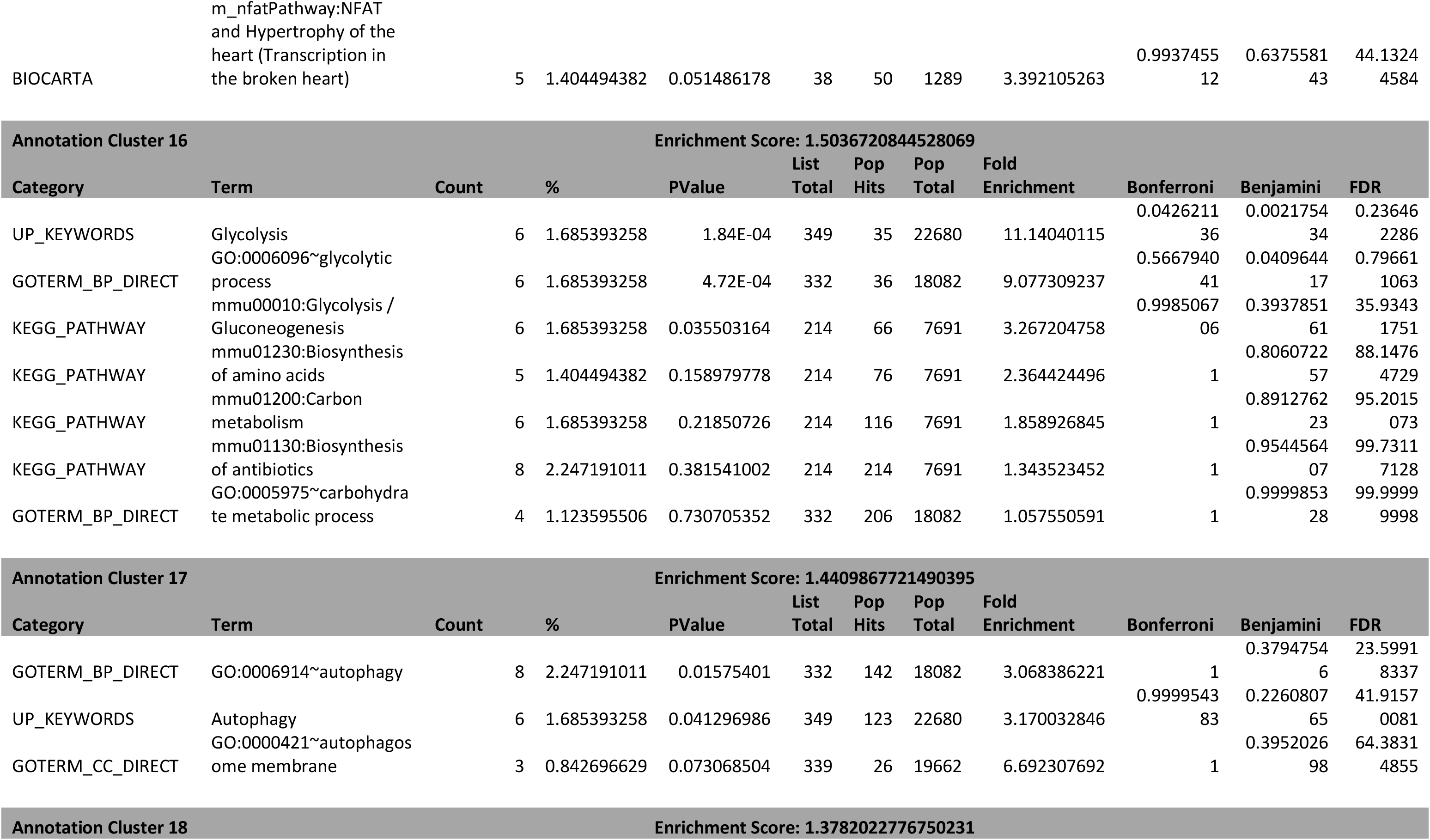
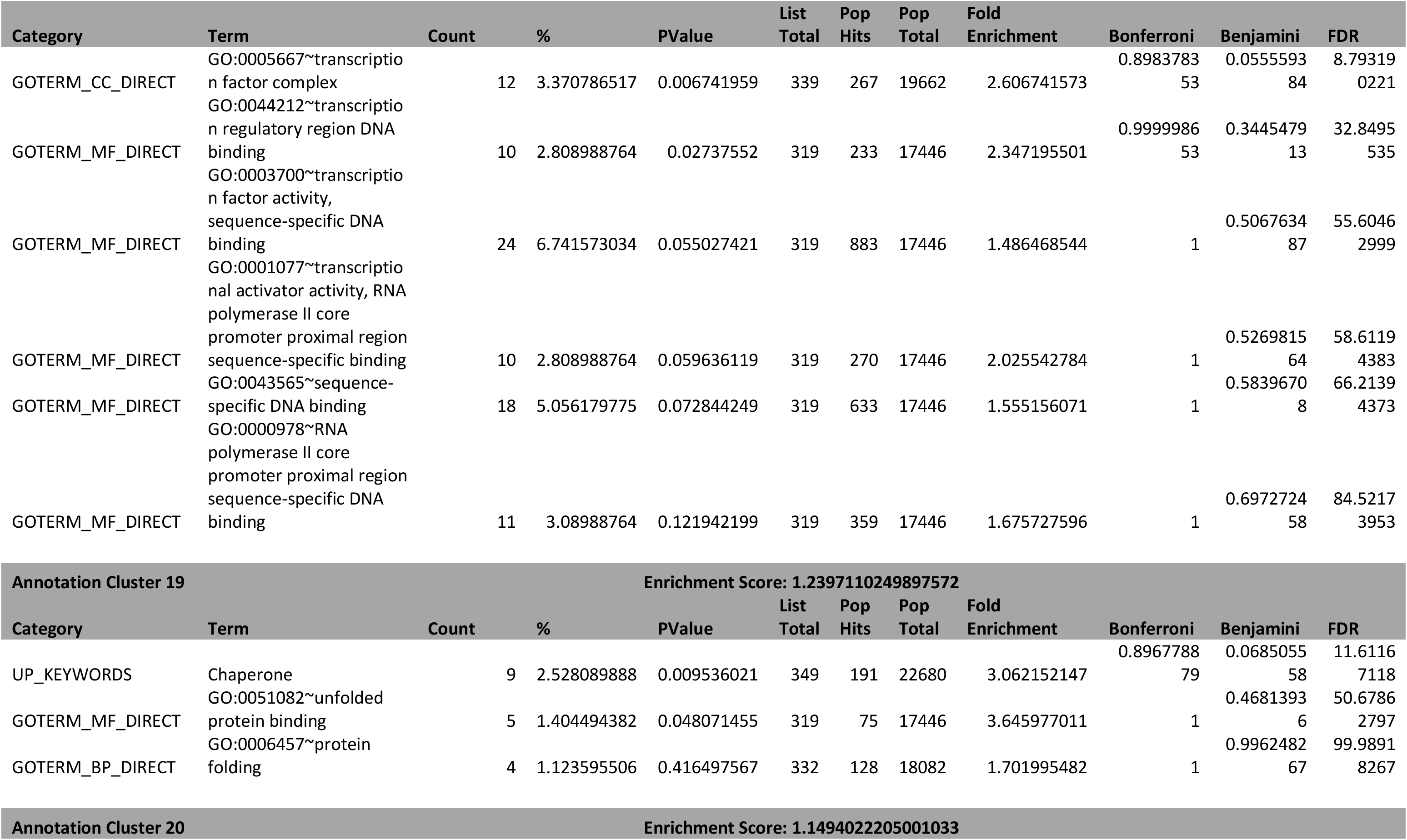
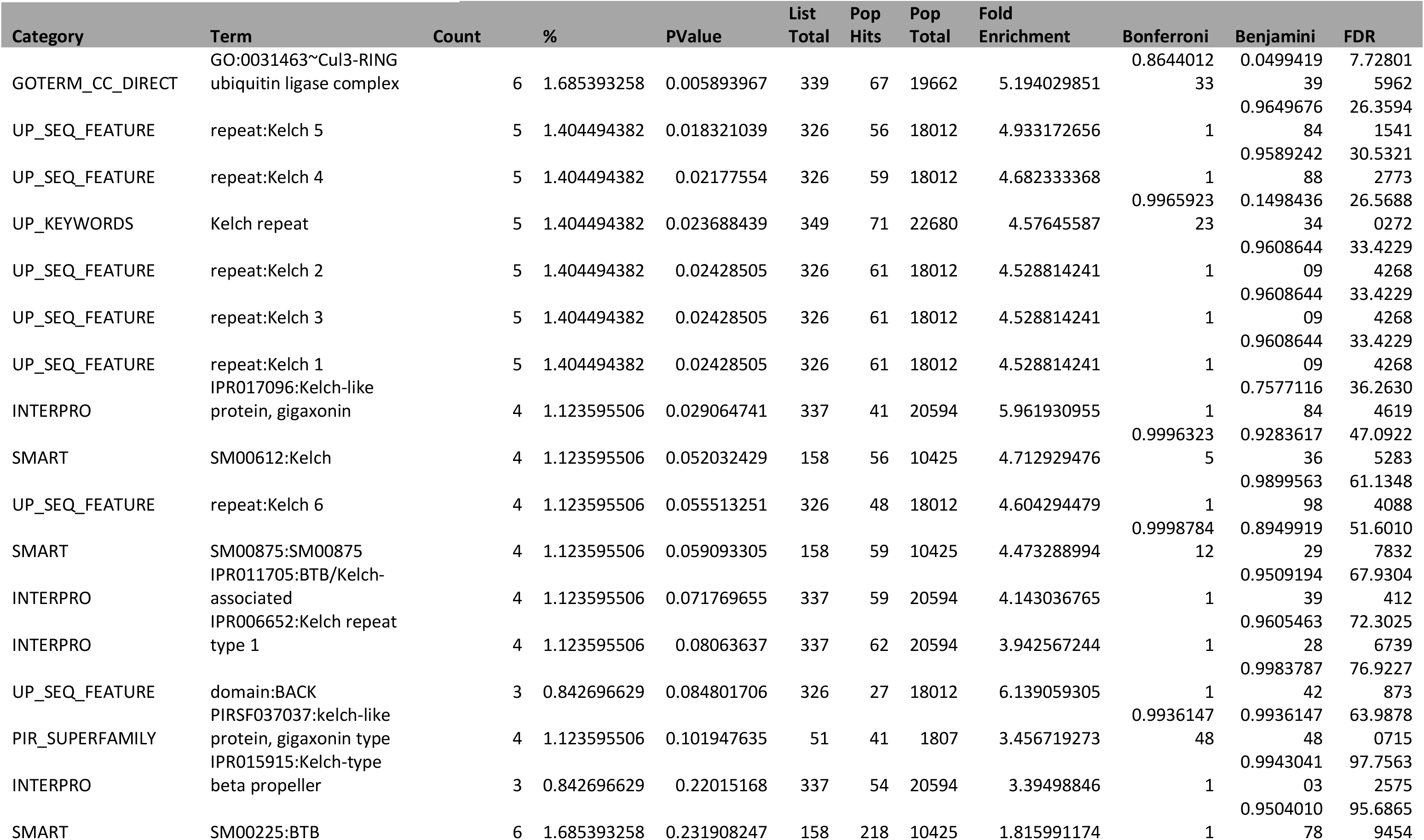
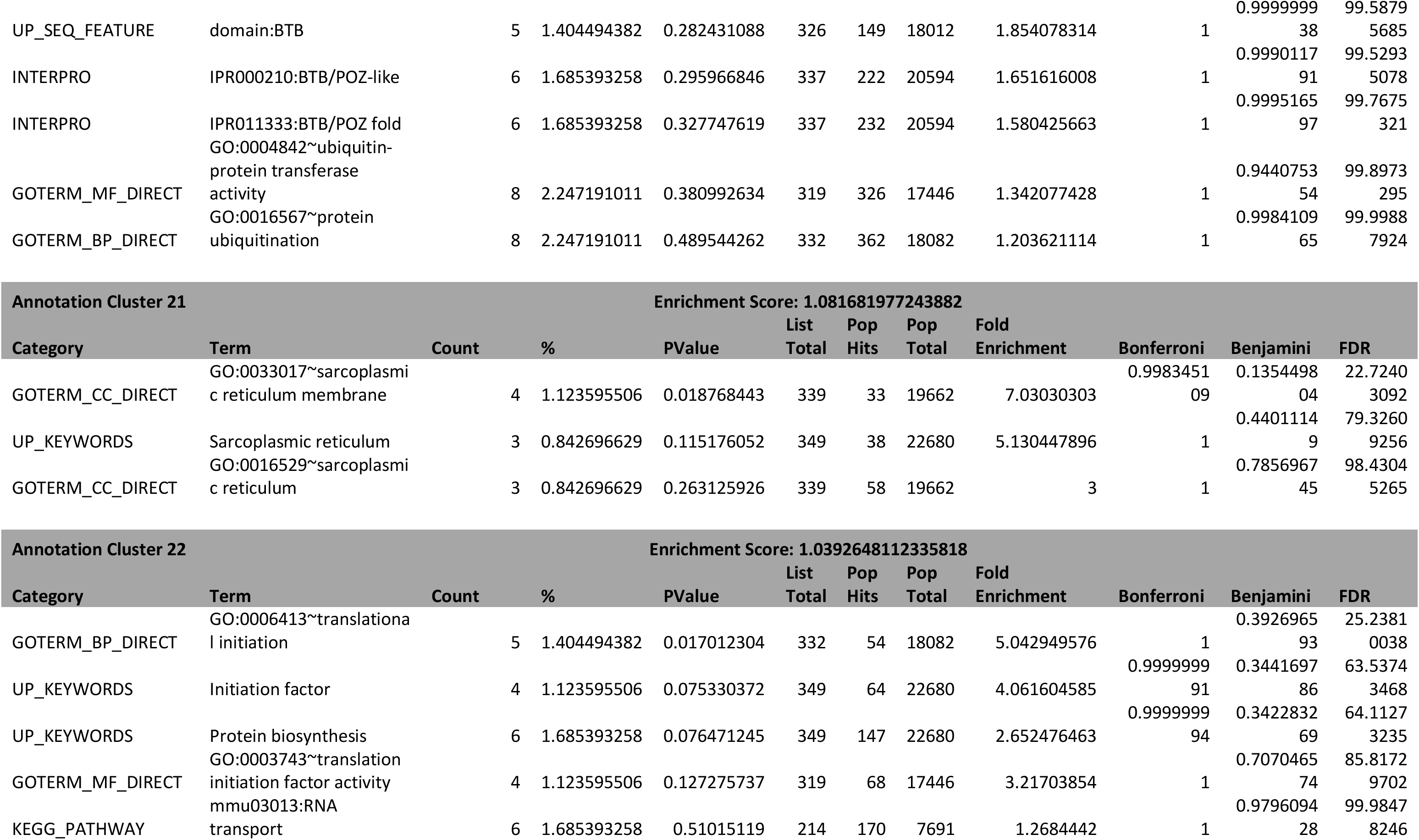
Go term analysis of genes exclusively expressed in the single myofiber, defined as genes with a q value lower than 0.01 between single fiber and whole muscle, more highly expressed in the single fiber with a Log2 fold change of less than -1, more highly expressed than 10 RPM and with a difference in expression between single fiber and whole muscle of at least 10 RPM. Used DAVID 6.8 online functional annotation tool with the ENSEMBL_GENE_ID identifier and with a classification stringency of medium.

**Table S3:** Alignment statistics of sequenced reads from young and old single myofibers.

| Sample | No Feature | Ambiguous | Unique | % overall alignment rate | % unique alignment |
| --- | --- | --- | --- | --- | --- |
| 1M SF1 | 6783051 | 6805642 | 31809772 | 85.06 | 70.07 |
| 1M SF2 | 7516309 | 8878533 | 35451356 | 85.50 | 68.38 |
| 3M SF1 | 9839266 | 5780069 | 24273800 | 75.34 | 60.85 |
| 3M SF2 | 8509055 | 13154637 | 31692716 | 84.05 | 59.40 |
| 3M SF3 | 6307020 | 3744695 | 16002158 | 75.79 | 61.42 |
| 19M SF1 | 14806391 | 15264491 | 48244515 | 81.09 | 61.60 |
| 19M SF2 | 5255899 | 5987465 | 31378173 | 87.67 | 73.62 |
| 19M SF3 | 7121590 | 11497612 | 28455416 | 84.87 | 60.45 |
| WM 1 | 2188178 | 1917476 | 9300343 | 83.68 | 69.37 |
| WM 2 | 6680813 | 4020883 | 22788368 | 80.05 | 68.05 |
| WM 3 | 7793097 | 4584231 | 24389338 | 78.80 | 66.34 |

